# Inhibition of ATR Reverses a Mitochondrial Respiratory Insufficiency

**DOI:** 10.1101/805390

**Authors:** Megan B. Borror, Milena Girotti, Adwitiya Kar, Meghan Cain, Xiaoli Gao, Vivian L. MacKay, Brent Herron, Shylesh Bhaskaran, Sandra Becerra, Nate Novy, Natascia Ventura, Thomas E. Johnson, Brian K. Kennedy, Shane L. Rea

**Author notes:** Address correspondence to: Shane Rea, PhD; University of Washington, Department of Pathology, 1959 NE Pacific St., Box 357470, Seattle WA 98195-7470. **Abbreviations: ATM**, ataxia telangiectasia mutated; **ATFS-1**, Activating Transcription Factor associated with Stress; **ATL-1**, ataxia telangectasia mutated-Like (*C. elegans* ortholog of ATR); **ATR**, ataxia telangiectasia mutated and Rad3-related; **DDR**, DNA damage response; **DHODH**, dihydroorotate dehydrogenase; **DNA pol** α, DNA polymerase alpha; **DNA polA1**, DNA polymerase alpha catalytic subunit; **ETC**, electron transport chain; **mtDNA**, mitochondrial DNA; **MAPKAP**, p38 MAPK-activated protein kinase; **mTOR**, mechanistic target of rapamycin; **PIKK**, phosphoinositide 3-kinase-related kinase; **R-loop**, DNA::RNA hybrid; **RNA pol II**; RNA polymerase II; **TUNEL**, Terminal deoxynucleotidyl transferase dUTP nick end labeling.

## Abstract

Diseases that affect the mitochondrial electron transport chain (ETC) often manifest as threshold effect disorders, meaning patients only become symptomatic once a certain level of ETC dysfunction is reached. Multiple processes work to control proximity to the critical ETC threshold and as a consequence there can be significant variability in disease presentation among patients. Identification of such control processes remains an ongoing goal. Checkpoint signaling comprises a collection of alert mechanisms activated in cells in response to nuclear DNA damage. Well-defined hierarchies of proteins are involved in both sensing and signaling DNA damage, with ATM (ataxia telangiectasia mutated) and ATR (ATM and Rad3-related) acting as pivotal signaling kinases. In the nematode *C. elegans*, severe reduction of mitochondrial ETC activity shortens life, as in humans, but mild reduction extends life as a consequence of survival strategies that are invoked under these circumstances. Here we show that removal of ATL-1, the worm ortholog of ATR, unexpectedly lessens the severity of ETC dysfunction, but removal of ATM does not. Multiple genetic and biochemical tests show no evidence for increased mutation or DNA breakage in animals exposed to ETC disruption. Instead, we find that reduced ETC function alters nucleotide ratios within both the ribo- and deoxyribo-nucleotide pools, and causes stalling of RNA polymerase, which is also known to activate ATR. Unexpectedly, *atl-1* mutants confronted with mitochondrial ETC disruption maintain normal levels of oxygen consumption and have an increased abundance of translating ribosomes. This suggests checkpoint signaling by ATL-1 normally dampens cytoplasmic translation. Taken together, our data suggests a model whereby ETC insufficiency in *C. elegans* results in nucleotide imbalances leading to stalling of RNA polymerase, activation of ATL-1, dampening of global translation and magnification of ETC dysfunction. Loss of ATL-1 effectively reverses the severity of ETC disruption so that animals become phenotypically closer to wild type.

## INTRODUCTION

The essential nature of mitochondria is underscored by the devastating effects of inherited mitochondrial diseases which afflict approximately one in 5000 children and almost always result in pathological shortening of life ^1^. Because these organelles play central roles in many cellular processes, including energy production, nucleotide metabolism and apoptotic signaling ^2, 3, 4^, their progressive, age-dependent reduction in function means almost all of us, at some time in life, will experience the negative consequences of mitochondrial dysfunction ^5^. Indeed a growing number of age-related ailments common to western societies are either associated with, or directly caused by, mitochondrial disruption ^6, 7^.

In *Caenorhabditis elegans*, as in humans, mitochondrial dysfunction is threshold dependent. That is, a critical level of electron transport chain (ETC) dysfunction must be reached before pathology is observed ^8^. Nematodes with high levels of mitochondrial dysfunction exhibit shortened lifespan and reduced fitness, while worms with moderate levels of mitochondrial dysfunction display extended lifespan ^9^. This response is evolutionarily conserved from yeast to mammals and understanding the molecular mechanisms regulating lifespan extension in worms in response to mild ETC disruption may provide insight into strategies that could be applied to help keep our own cells above their critical mitochondrial dysfunction threshold.

Reduced ETC activity disrupts many downstream cellular processes and as might be expected cells have evolved a variety of sensors and strategies to counter progression toward their critical ETC threshold. Multiple signaling pathways, collectively called retrograde responses, are activated within cells and these in turn control expression of nuclear counter-measures ^10^. For example, an increase in the unfolded protein load of the mitochondrial matrix re-directs the transcription factor ATFS-1 from the matrix to the nucleus where it stimulates expression of mitochondrial chaperones ^11^. Other signals such as changes in calcium concentration, activation of mitogen-activated protein kinases (MAPKs), and even knock-on changes in the cytoplasmic unfolded protein load, also result in retrograde response activation ^12, 13, 14^. Counter-measures that are activated by cells include increased mitochondrial biogenesis, elevated mitochondrial DNA (mtDNA) replication, increased mitophagy, activation of alternate pathways of energy production (such as glycolysis), and changes in the abundance and activity of respiratory complexes or their regulatory factors ^8, 15, 16^. The complete network of pathways that detect and respond to mitochondrial stress is, however, far from being fully understood ^17^.

Previously, we described a phenomenon in worms in which loss of CEP-1, a homolog of the human p53 checkpoint protein that recognizes identical DNA sequences and is the only p53 family member present in *C. elegans* ^18^, right shifted the mitochondrial ETC threshold, effectively allowing animals to cope with a greater degree of mitochondrial ETC disruption and consequently mitigating the lifespan shortening effects of severe mitochondrial dysfunction ^19^. This response was due to alterations in autophagy and lipid metabolism, both of which are regulated by CEP-1 ^20^. In the current study, we examined the role of other checkpoint proteins in modulating the mitochondrial threshold effect of worms since this is still a little-explored area of investigation.

Ataxia-telangiectasia mutated (ATM) and ‘ATM- and *rad3*-related’ (ATR) are two well-studied members of the phosphoinositide 3-kinase-related kinase (PIKK) family of proteins that play essential roles in transducing checkpoint signaling during the DNA damage response (DDR) ^21^. In humans, both proteins together directly phosphorylate over 900 sites on some 700 proteins, including p53 ^22^. ATM and ATR differ in the lesions to which they are recruited, with ATM being recruited to dsDNA breaks and ATR having a broader specificity underscored by the presence of ssDNA, including recessed dsDNA breaks and stalled replication forks ^23^. Depending on the level of damage and success of repair, checkpoint activation can lead to pro-survival or pro-apoptotic mechanisms ^24^. Recently, additional functions were ascribed to both ATM and ATR. For ATM, these involve regulation of a variety of processes, including insulin signaling, the pentose phosphate pathway and mitophagy. Also, in yeast, ATM/Tel1p was found to act as a specific sensor of mitochondria-generated ROS ^25, 26, 27, 28, 29^. ATR on the other hand was shown to be important in controlling an outer mitochondrial membrane (OMM)-localized apoptotic signal, autophagy, RNA polymerase activity and chromatin condensation ^30, 31, 32, 33, 34^. In this study, we uncover a new role for ATR in mitochondrial retrograde response signaling.

## RESULTS

### Loss of ATL-1/ATR desensitizes worms to mitochondrial ETC stress

In *C. elegans, atp-3* encodes the ortholog of the human ATP5O/OSCP subunit of the mitochondrial F_1_F_o_ ATP synthase, while *isp-1* encodes the Rieske Fe-S ortholog of complex III. We have previously reported that wild type worms exposed to increasing amounts of bacterial feeding RNAi targeting either *atp-3* or *isp-1* reach a critical threshold after which the concentration of RNAi causes worms to exhibit a reduction in body size and an extension of lifespan ^35^. For *isp-1* RNAi, life extension follows a monotonic function, while for *atp-3* RNAi, life is first extended then it is pathologically shortened ^35^. These ‘mean lifespan versus feeding RNAi dosage’ curves were mapped using 12 point datasets and their profiles were robust across multiple rounds of testing ^35^. This differing effect of *atp-3* and *isp-1* feeding RNAi on lifespan reflects the more potent degree of ETC inhibition that is ultimately attained following severe *atp-3* knockdown. Using these well-defined treatment regimes and reagents, we tested whether loss of ATM-1 or ATR checkpoint proteins altered the phenotypic response of worms to mitochondrial ETC disruption (see **Supplemental File S1** for a full description of mutants used in this study and the method used to the propagate lethal DDR strains). When mutant *atm-1(gk186)* worms were exposed to *atp-3* or *isp-1* RNAi (1/10^th^ strength and undiluted) animals matured and survived indistinguishably from similarly-treated wild type worms (Fig. 1A-D). In contrast, *atl-1(tm853)* mutants were differentially refractory to the effects of knockdown of either RNAi. This effect was most evident in worms treated with 1/10^th^ strength *atp-3* or *isp-1* RNAi, where animals showed obvious resistance to both size reduction (Fig. 1A, B) and lifespan extension (Fig. 1C, D). (Additional examples are presented later in Figure 2A-C). *atp-3* and *isp-1* knockdown did eventually reduce adult size and extend life in the mutant *atl-1(tm853)* background, suggesting the response to ETC disruption in these animals is right-shifted relative to wild type control (see schematic Fig. 1C, D). Surprisingly, measurement of *atp-3* and *isp-1* mRNA levels following RNAi treatment revealed that target gene knockdown was as efficacious in *atl-1(tm853)* mutants as it was in wild type worms (Fig. 1E), indicating these animals were not deficient for RNAi. Furthermore, we found that *atl-1(tm853)* mutants were also refractory to mitochondrial ETC disruption induced using chemical inhibition (**Supplemental Figure S1**). Based on these findings, we conclude that inhibition of the phosphoinositide 3-kinase-related kinase ATL-1 confers resistance to mitochondrial ETC disruption.

**Figure 1.**
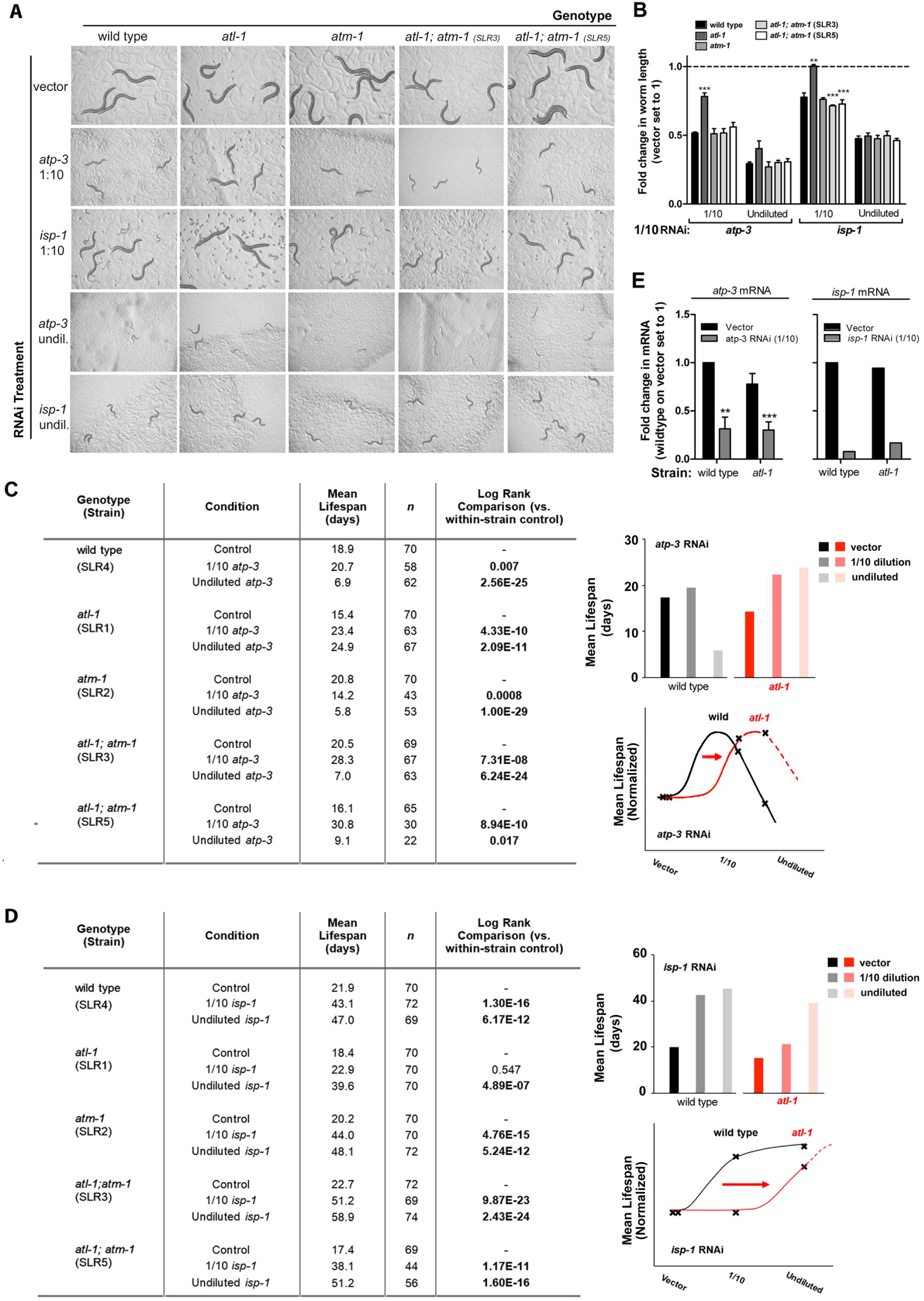
Loss of ATL-1 desensitizes worms to mitochondrial respiratory chain stress. (**A, B**) *atl-1(tm853)* knockout mutants are resistant to size reduction induced by *atp-3* or *isp-1* bacterial feeding RNAi (compare 1/10^th^ strength RNAi to vector-treated control, *panel A*). Tested worms include wild type, *atm-1(gk186)* and *atl-1(tm853)* single mutants, as well as two independent *atl-1(tm853); atm-1(gk186)* lines. Worm length is quantified in *panel B*. All worms were seeded at the same initial time. Each row was photographed at a common time interval after seeding, and this time interval increased with increasing RNAi potency. Evident across rows are differences in growth rates when strains are cultured on the same RNAi treatment. Error bars: SEM. Significance testing: Student’s t-test. Asterisks refer to difference relative to respective wild type sample: **p<0.01, ***p<0.001. (**C, D**) *Charts on Left:* Adult lifespan of worm strains tested in (A). Shown is a representative study. All conditions were tested in parallel derived from a starting population of 16,000 synchronous eggs. Replicate data for *atp-3* is presented in Figure 2. *Right panels*: Schematics illustrating the right-shifting effect of ATL-1 removal on mean lifespan following exposure to *atp-3* and *isp-1* RNAi. Curve shapes are based on the 12-point RNAi data collected in Rea *et. al*. ^35^. (**E**) RNAi knockdown efficacy is not altered in *atl-1(tm853)* mutants. mRNA levels were quantified by q-RT PCR following exposure to 1/10^th^ strength *atp-3* or *isp-1* bacterial feeding RNAi (n=3 and n=1 experimental replicates, respectively). Error bars: SEM. Significance testing: Student’s t-test. Asterisks refer to difference relative to wild type vector control **p<0.01, ***p<0.001. In (A-E), all worms are unbalanced animals (∼1,000) derived from F1 progeny of parents carrying the nT1 reciprocal chromosomal translocation. These worms were hand selected from the 16,000 eggs after they matured into adults. The wild type control line (SLR4), was deliberately moved into the nT1 genetic background as the appropriate control (see *Materials & Methods*).

**Figure 2.**
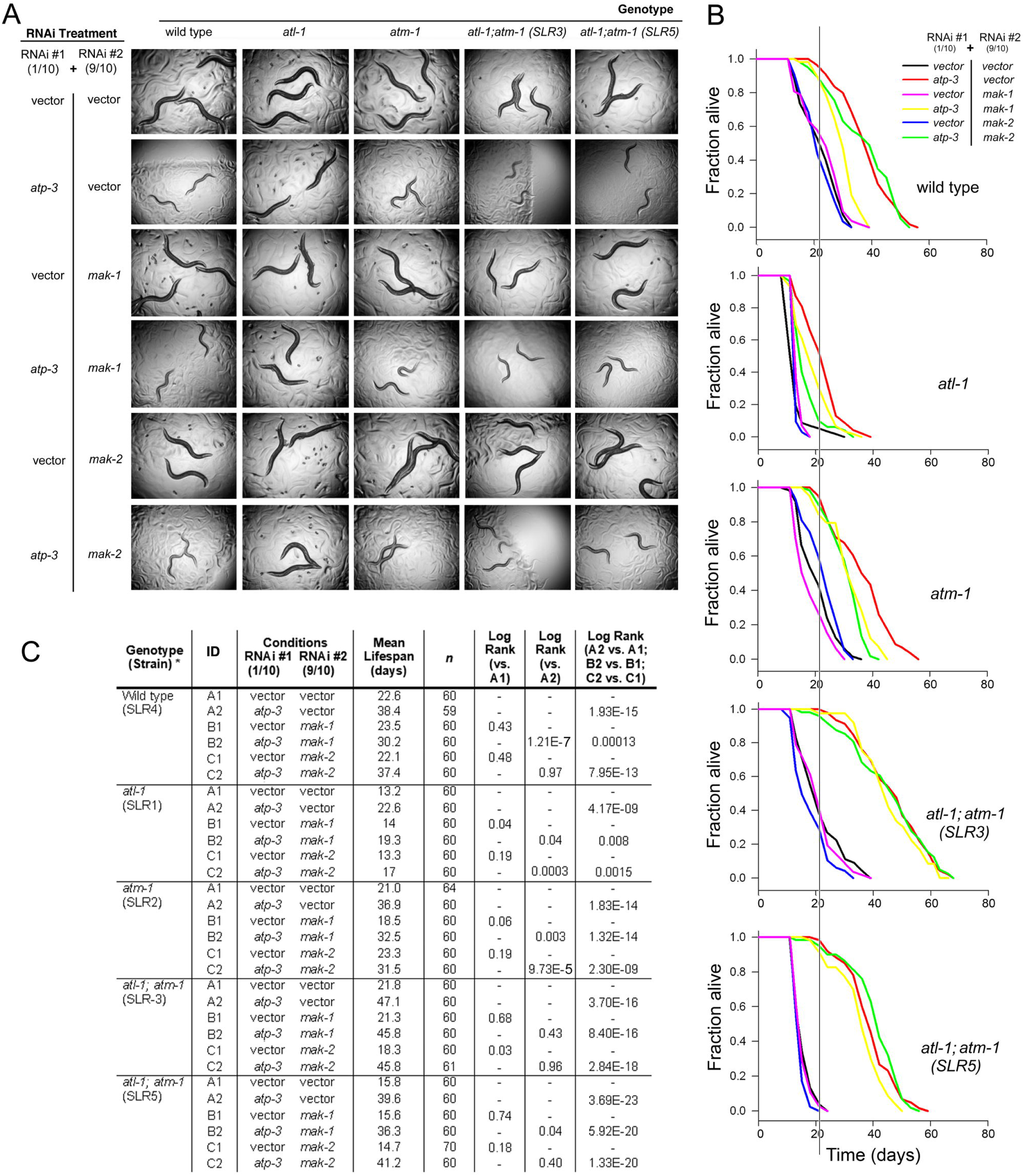
MAPKAP kinases are active following mitochondrial respiratory chain stress. (**A-C**) The MAPKAP kinases MAK-1 and MAK-2 play no role in size control of worms exposed to 1/10^th^ strength *atp-3* feeding RNAi (A), however they do control lifespan in wild type worms, *atm-1 (gk186)* mutants and *atl-1(tm853)* mutants (B, C). Worms in (A) and (B) were established independently using a total of 8,000 and 32,000 synchronized eggs, respectively, and all conditions in each panel were tested in parallel. In (A), all worms were photographed at the same chronological point. In (B), the vertical line marks median survival of wild type worms on vector control RNAi. In (A-C), all worms were the unbalanced F1 progeny of parents carrying the nT1 reciprocal chromosomal translocation.

### MAK-1 and MAK-2 participate in mitochondrial retrograde response signaling

During the course of our studies we made the unexpected observation that two independently-generated *atm-1(gk186); atl-1(tm853)* double mutants regained their sensitivity to *atp-3* and *isp-1* knockdown (Fig. 1). This observation provides additional support for the notion that the RNAi machinery is unaffected by loss of *atl-1,* but to explain this unexpected result, we have formed the following working hypothesis: ETC disruption results in DNA damage that normally activates ATR. In the absence of ATR, DNA damage accumulates, or secondary forms of DNA damage accumulate, and eventually ATM is activated. In the absence of both ATR and ATM, a third checkpoint response, with a higher threshold for activation, is triggered. This model reflects the well-established redundancy in the DNA damage response network ^22^. Several recent studies by us and others ^13, 36, 37^, have shown a role for MAPK signaling following ETC disruption in worms. Interestingly, the p38 MAPK-activated protein kinases MAPKAP-2 and MAPKAP-3 are checkpoint proteins known in humans to act in parallel to ATM and ATR ^38, 39, 40^. To explore the possibility that alternate checkpoint signaling is activated in PIKK mutant worms following ETC disruption, we exposed animals to *atp-3* knockdown and simultaneously inhibited expression of either MAK-1 or MAK-2, the worm orthologs of MAPKAP-2 and -3, respectively. The effect of these treatments on *alt-1(tm853)* and *atm-1(gk186)* single mutants, *atm-1(gk186); atl*-*1(tm853)* double mutants, and wild type worms was examined. Our results can be summarized as follows: (i) Knockdown of *mak-1* or *mak-2*, either alone or in combination with *atp-3* knockdown, had no effect on final adult size in any of the genetic backgrounds (Fig. 2A). (ii) Knockdown of *mak-1* or *mak-2* failed to reproducibly prevent life extension following *atp-3* inhibition in the two independent *atm*-*1(gk186); atl-1(tm853)* double mutant isolates. (iii) Unexpectedly, the survival of *atl-1(tm853)* mutants exposed to *atp-3* knockdown was further impaired when undertaken in conjunction with *mak-1* or *mak-2*. A similar result was also observed for the *atm-1(gk186)* mutant. (iv) For wild type worms, only *mak-1* knockdown significantly reduced *atp-3* mediated life extension (Fig. 2B, C). These findings suggest that at least two other checkpoint proteins, namely MAK-1 and MAK-2, are causally involved in lifespan extension (but not size determination) of worms experiencing mitochondrial ETC disruption. The role of different checkpoint proteins is therefore dependent upon the functional status of both ATL-1 and ATM-When the latter two proteins are disrupted simultaneously, MAK-1 and MAK-2 become dispensable, potentially implying yet another checkpoint system awaits identification in *atm-1(gk186); atl-1(tm853)* double mutants. Although clearly complicated, this is not unexpected for a response that is network-based and sensitive to a variety of DNA lesions.

### Knockdown of DNA pol α reverses the small phenotype of *atm-1(gk186); atl-1(tm853)* mutants

To further explore the role of novel checkpoint proteins in modulating the critical ETC threshold in *atm*-*1(gk186); atl-1(tm853)* double mutants, we undertook a double RNAi screen of 201 DNA damage response-related genes. We asked whether knockdown of a target gene in this genetic background could re-confer resistance to *atp-3* knockdown, similar to *atl-1(tm853)* mutants (for screen details see Fig. 3A). We identified two genes: *scc-3,* which encodes a cohesin complex subunit; and Y47D3A.29, which is orthologous to the human PolA1 catalytic subunit of DNA polymerase alpha (DNA pol α, **Figs. 43, C**). The effect of losing either gene was robust to three different post-hoc tests (each p<0.001, see **Tables S1-S3** and **Methods** for details of statistical testing; see also **File S2** for final round hit dataset). Isolation of PolA1 is particularly intriguing because it suggests that *atp-3* knockdown in *atm-1(gk186); atl-1(tm853)* double mutants could be inducing some kind of disruption to DNA, its repair, or its replication and which is by-passable by low fidelity polymerases (which presumably accommodate knockdown of PolA1). This in turn might avoid checkpoint activation and make animals appear more *atl-1(tm853*)-like in our assay.

**Figure 3.**
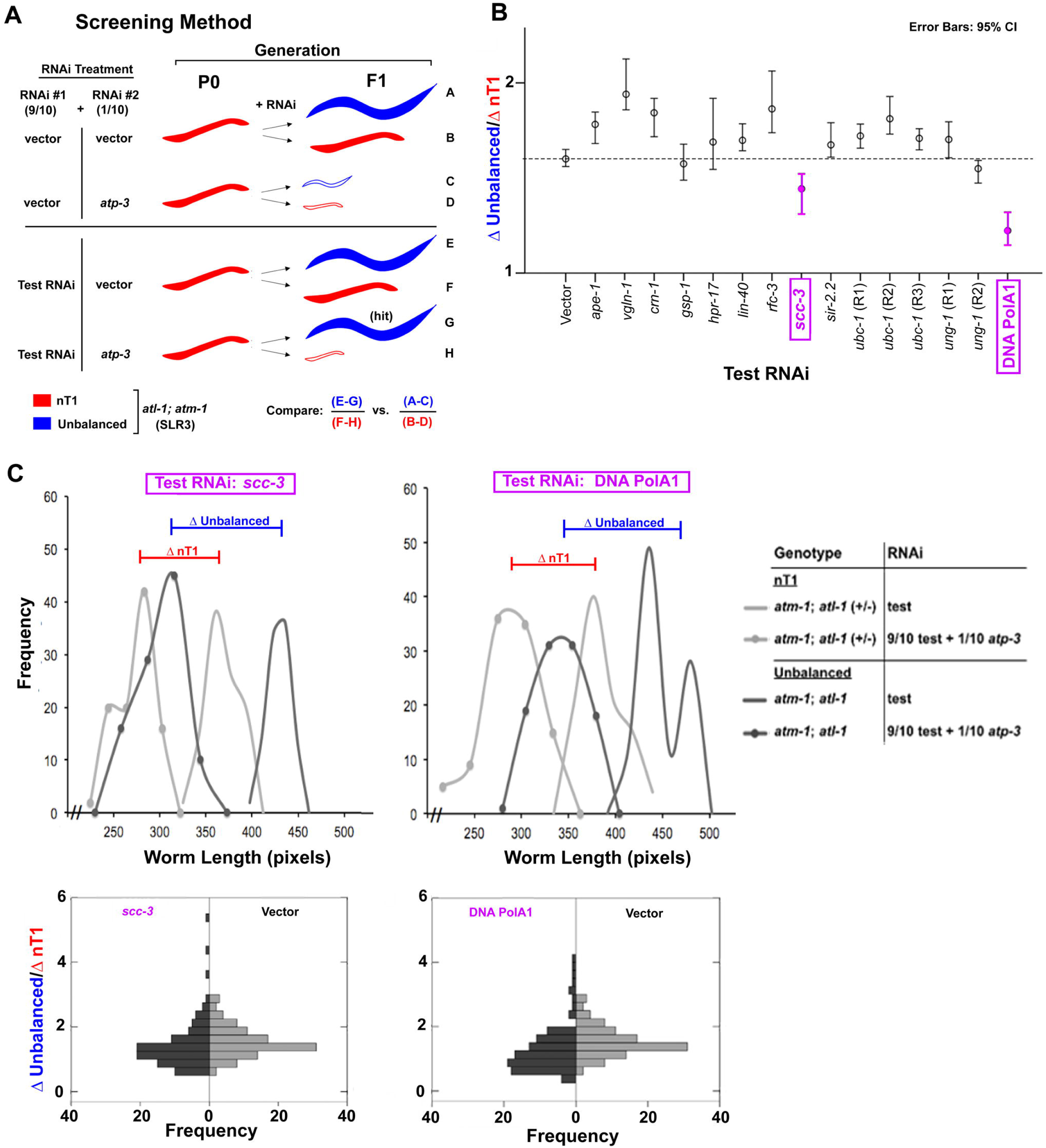
Screen for additional checkpoint proteins activated by mitochondrial ETC dysfunction. (**A**) Schematic of RNAi screen used to identify DNA damage response genes active in *atl-1(tm853); atm-1(gk186)* double mutants following *atp-3* knockdown. Test RNAi that differentially countered the size reducing effect of 1/10^th^ strength *atp-3* RNAi in unbalanced *atl-1(tm853); atm-1(gk186)* worms but not nT1-containing *atl-1(tm853); atm-1(gk186)/+* worms, were sought (hit). (**B**) Results of RNAi screen. The effect of each test RNAi on the length of unbalanced worms and nT1-containing worms in the presence and absence of 1/10^th^ strength *atp-3* RNAi is reported as a mean difference ratio. The 95% confidence interval (CI) for the ratio is shown. RNAi knockdown of *scc-3* and *Y47D3A.29* (labeled DNA PolA1) both significantly reduced the mean ratio relative to vector-only test RNAi (p< 0.001, see Methods for significance testing procedure). (**C**) *Top row:* Size distribution of worm populations treated with *scc-3 (left)* or *Y47D3A.29 (right)* feeding RNAi. *Bottom row:* Histograms of mean difference ratios for worm populations treated with *scc-3 (left)* or *Y47D3A.29 (right)* feeding RNAi and plotted relative to worms treated with vector in place of test RNAi. Mean difference ratios were generated by random sampling from the distributions in the *top row*.

### No evidence for permanent nuclear DNA damage following mitochondrial ETC disruption

Nuclear DNA (nDNA) damage is the canonical signal that results in activation of ATR. In worms, resected ends of dsDNA breaks, and stalled replication forks, have both been shown to activate ATL-1^41^. Both lesions contain single-stranded DNA (ssDNA), and binding of heterotrimeric replication protein A (RPA) to these regions in turn recruits and activates ATL-1. To test if ETC disruption results in elevated levels of dsDNA breaks we focused on *isp-1(qm150)* genetic mutants, since no viable *atp-3* mutant exists, and utilized TUNEL staining on sections of paraffin-embedded worms ^42^. We found no evidence for increased nDNA breakage (**Supplemental Fig. S2A**). DNA repair is sometimes inaccurate, depending on the repair machinery that is employed. To test if DNA mutation frequency was increased in worms exposed to ETC disruption we employed four independent genetic assays. To facilitate analysis, these studies were undertaken using *atp-3* RNAi. The eT1 (III;V) and nT1(IV;V) reciprocal chromosomal translocations each suppress recombination against the regions they balance (∼15 Mbp for both balancers). Homozygous-lethal DNA mutations therefore accumulate on balanced chromosomes and they become identifiable when eT1 and nT1 chromosomal pair are lost because unbalanced progeny that are homozygous for the mutation are inviable and hence absent. Using both assays, we found no evidence for increased nDNA mutation rate following knockdown of *atp-3* in wild type worms (Fig. 4A). Similarly, use of two genetic mutation (Unc) reversion assays yielded similar conclusions (Fig. 4B). Taken together, our data show there is neither an increase in DNA strand breakage nor is there an increase in nuclear DNA mutation frequency in worms experiencing mitochondrial ETC dysfunction.

**Figure 4.**
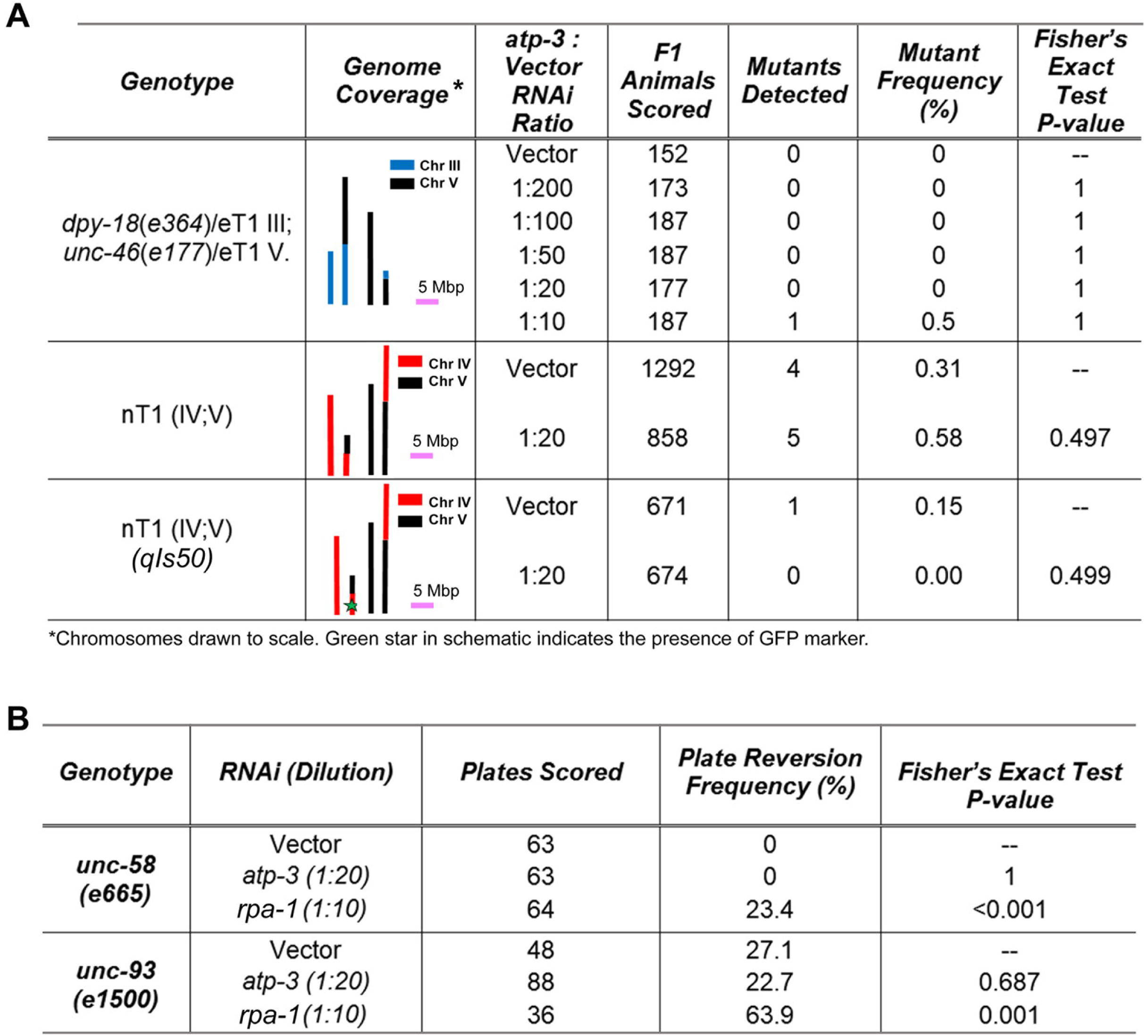
Nuclear DNA mutation rate is not increased following mitochondrial ETC disruption. (**A, B**) Multiple genetic screens reveal no significant increase in nuclear DNA mutation frequency following knockdown of *atp-3* (p>0.05, Fisher’s Exact Test). Phenotypes scored include lethal nuclear DNA mutation events covered by the eT1 and nT1 chromosomal translocations (A), and recovery of wild type movement in *unc-58(3665*) and *unc-93(e1500)* mutants (B). Knockdown of replication protein A (*rpa-1*) was included as a positive control in (B).

### Ribonucleotide and deoxyribonucleotide pools are disrupted by mitochondrial ETC dysfunction

To search for alternate mechanisms by which ATL-1 might be activated in worms experiencing mitochondrial ETC dysfunction, we examined changes in whole-worm nucleotide levels. As in humans, pyrimidine biosynthesis in *C. elegans* requires a functional mitochondrial electron transport chain. This is because formation of orotate from dihydroorotate is coupled to reduction of ubiquinone (UQ) in a reaction catalyzed by the inner mitochondrial membrane enzyme dihydroorotate dehydrogenase (DHODH, **Supplemental Fig. S3**). In mammals, processes that slow flux through the mitochondrial ETC, downstream of UQ, also slow the rate of dihydroorotate dehydrogenase resulting in measurable disruption of nucleotide pools ^43^. Several important enzymes are sensitive to changes in nucleotide abundance, including DNA polymerases which are rate-limited by availability of their deoxyribonucleotide substrates ^44^. Stalling of DNA polymerase, followed by continued DNA unwinding by the MCM helicase, results in ssDNA formation ^45, 46^, which is a known substrate for ATR activation.

To measure nucleotide levels, we re-focused our attention on the *isp-1(qm150)* genetic mutant for ease of culture. We interrogated three stages of development – the final two stages of larval development (L3 and L4) and the first day of adulthood. These stages were selected because in wild type worms there is a heavy demand for nucleotide precursors around the L3/L4 molt, and then again around the L4/adult molt. At these times the gonad arms are expanding and there is also a concomitant five-fold and a 6-fold increase in mitochondrial DNA (mtDNA), respectively ^47^. Among nine deoxyribonucleotide species that we were able to resolve and reliably detect (see **Methods**) we observed significant increases in dCDP and dTMP in L4 larvae of *isp-1(qm150)* mutants (Fig. 5A). The increase in dCDP abundance was maintained into adulthood (Fig. 5B). Among eleven ribonucleotide species that we were able to reliably resolve and detect, we observed significant increases in AMP, CMP and UMP in L4 larvae of *isp-1(qm150)* mutants (Fig. 5C). This was accompanied by a significant decrease in the level of GTP. In adult *isp-1(qm150)* worms, changes in ribonucleotide pools became more pronounced, with eight species being significantly underrepresented – ADP, ATP, GDP, GTP, CMP, CTP, UDP, UTP (Fig. 5D). Changes in nucleotide pools were specific to L4 and adult worms since we did not detect any measureable differences in any quantifiable nucleotide species in L3 larvae of *isp-1(qm150)* mutants relative to wild type worms (Fig. 5E). Finally, the absence of a corresponding increase in many of the monophosphate ribonucleotide species in adult *isp-1(qm150)* worms when their corresponding di- and triphosphate counterparts decreased, suggests ribonucleotide pool sizes are reduced in these animals. We combined the levels of each mono-, di- and triphosphate species and tested if they differed significantly between strains at each developmental stage, but they did not. Error propagation, however, severely reduced our statistical resolving power (Fig. 5F). Collectively, we conclude that disruption of the mitochondrial ETC in *C. elegans* results in measurable and significant changes in the abundance of multiple nucleotide species, notably with all ribonucleotide triphosphate species becoming underrepresented by adulthood.

**Figure 5.**
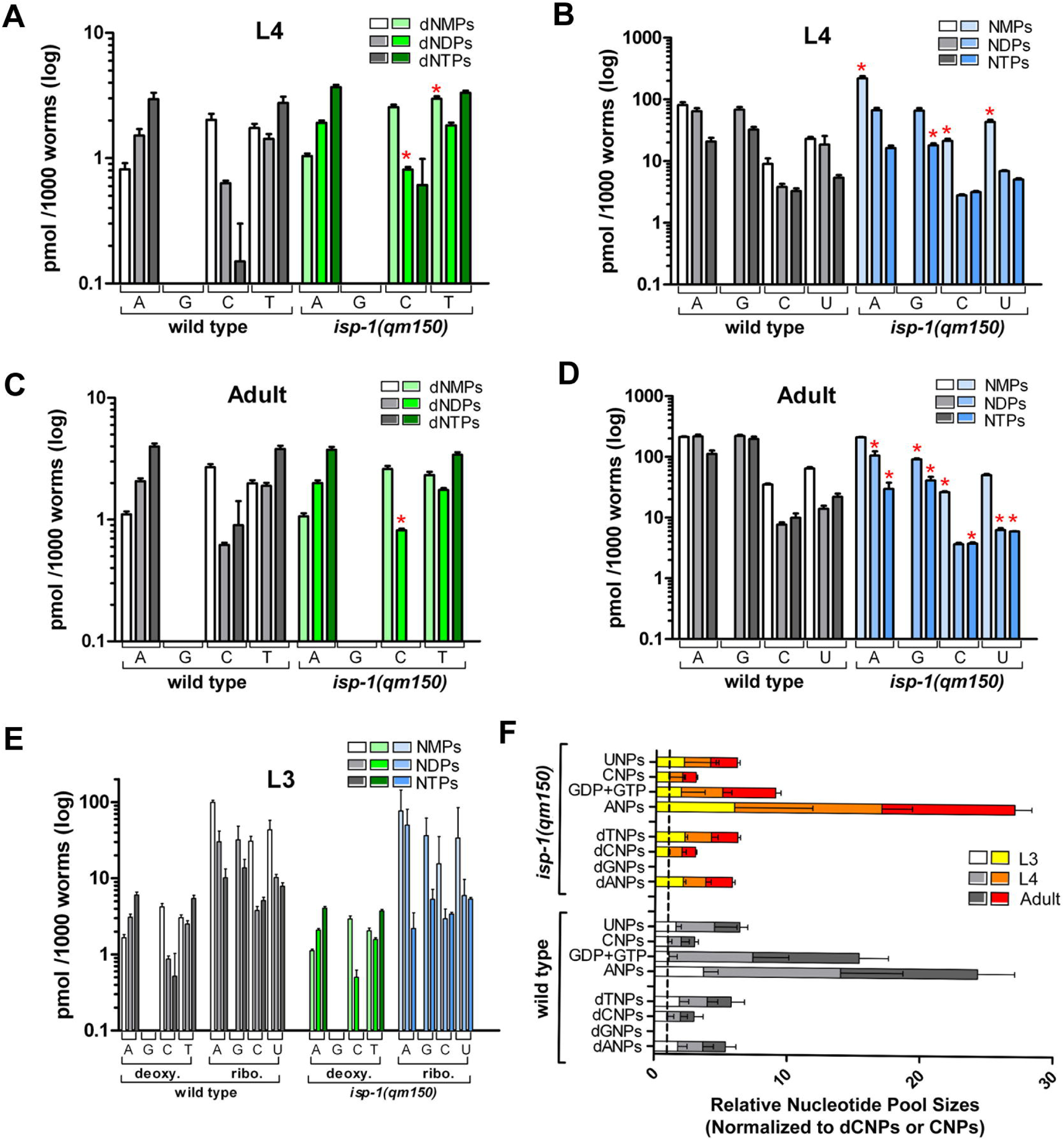
Ribonucleotide and deoxyribonucleotide pool ratios are disrupted following mitochondrial ETC disruption. (**A-E**) Absolute levels of deoxyribonucleotide (A and C) and ribonucleotide (B and D) species in wild type (N2) and *isp-1(qm150)* worms at the L4 and day-one adult stages, measured using quantitative LC-MS. Red asterisks indicate nucleotide quantities that differ significantly between strains (Student’s t-test, 5% FDR). Four nucleotides were not measurable by our assay (dGMP, dGDP, dGTP, GMP). No significant differences in any nucleotide level existed between strains at the L3 larval stage (E). (**F**) Relative nucleotide pool sizes in L3, L4 and day-one adult N2 and *isp-1(qm150)* worms, determined by summing together the mono-, di- and triphosphate species of each deoxyribonucleotide or ribonucleotide (guanosine was an exception) and normalizing to dCNP or CNP, respectively, across each larval stage. No significance difference in each respective nucleotide pool size between strains was detected (Student’s t-test, 5% FDR; N = 3, 7 and 3 experimental replicates for N2 L3, L4, and *a*dult worms; and N= 3, 5 and 3 for *isp-1(qm150)* L3, L4 and adult worms, respectively).

### Evidence for RNA polymerase stalling following mitochondrial ETC disruption

Stalling of RNA polymerase II (RNA pol II) during transcription is sufficient to activate an ATR, RPA and p53-dependent DNA damage response in human fibroblasts ^48^. Given that mitochondrial ETC dysfunction in *isp-1(qm150)* mutants leads to measurable reductions in all four ribonucleotide triphosphates, and because we have previously shown a role for p53/*cep-1* in the life extension of these worms ^19^, we hypothesized that ATL-1 activation following ETC disruption might occur as a result of RNA pol II stalling due to reduced nucleotide availability. We therefore quantified the formation of R-loops on genomic DNA ^49^. R-loops are DNA::RNA hybrids that accumulate when transcription stalls (Fig. 6A). We detected R-loops in purified, whole-worm genomic DNA fractions using slot blot analysis in conjunction with a DNA:RNA hybrid-specific antibody (αS9.6) ^50^. To obtain sufficient material for these studies we utilized a double feeding RNAi approach and tested four conditions: Wild type worms exposed to either vector control RNAi, 1/10^th^ strength *isp-1* RNAi, 9/10^th^ strength *atl-1* RNAi, or both 9/10^th^ strength *atl-1* RNAi with 1/10^th^ strength *isp-1* RNAi. Vector control feeding RNAi was added as balancer where appropriate. qRT-PCR analysis undertaken before the start of the experiment revealed 9/10^th^ strength *atl-1* RNAi reduced *atl-1* mRNA by 50% (Fig. 6B), the same amount that was observed in maternal-effect rescued *atl-1(tm853)* mutants (Fig. 6C). The results of our R-loop studies can be summarized as follows (Fig. 6D): Knockdown of *isp-1* significantly enhanced R-loop formation relative to vector control (Student’s t-test, p<0.05). As predicted, *isp-1* functioned epistatically to *atl-1,* since *atl*-*1* knockdown did not block R-loop formation by *isp-1*, as would be expected for a gene that functioned downstream of R-loop accumulation. We next tested whether any differences in DNA:RNA polymer length distinguished R-loops generated in worms exposed to *isp-1* RNAi from those exposed to *isp-1* and *atl-1* double RNAi. Genomic fractions were separated on an agarose gel, stained with ethidium bromide, then transferred to nitrocellulose and probed with αS9.6 antibody (Fig. 6E). No overt difference distinguished the two samples, however in both instances we noted some of the staining with αS9.6 localized to low molecular weight DNA:RNA fragments (< 2kb), in addition to the higher molecular weight structures that we expected. The identity of these smaller fragments remains uncharacterized (see **Discussion**). Collectively, our studies suggest that transcriptional stalling is increased in worms exposed to mitochondrial ETC disruption, identifying one route by which ATL-1 is potentially activated in these animals.

**Figure 6.**
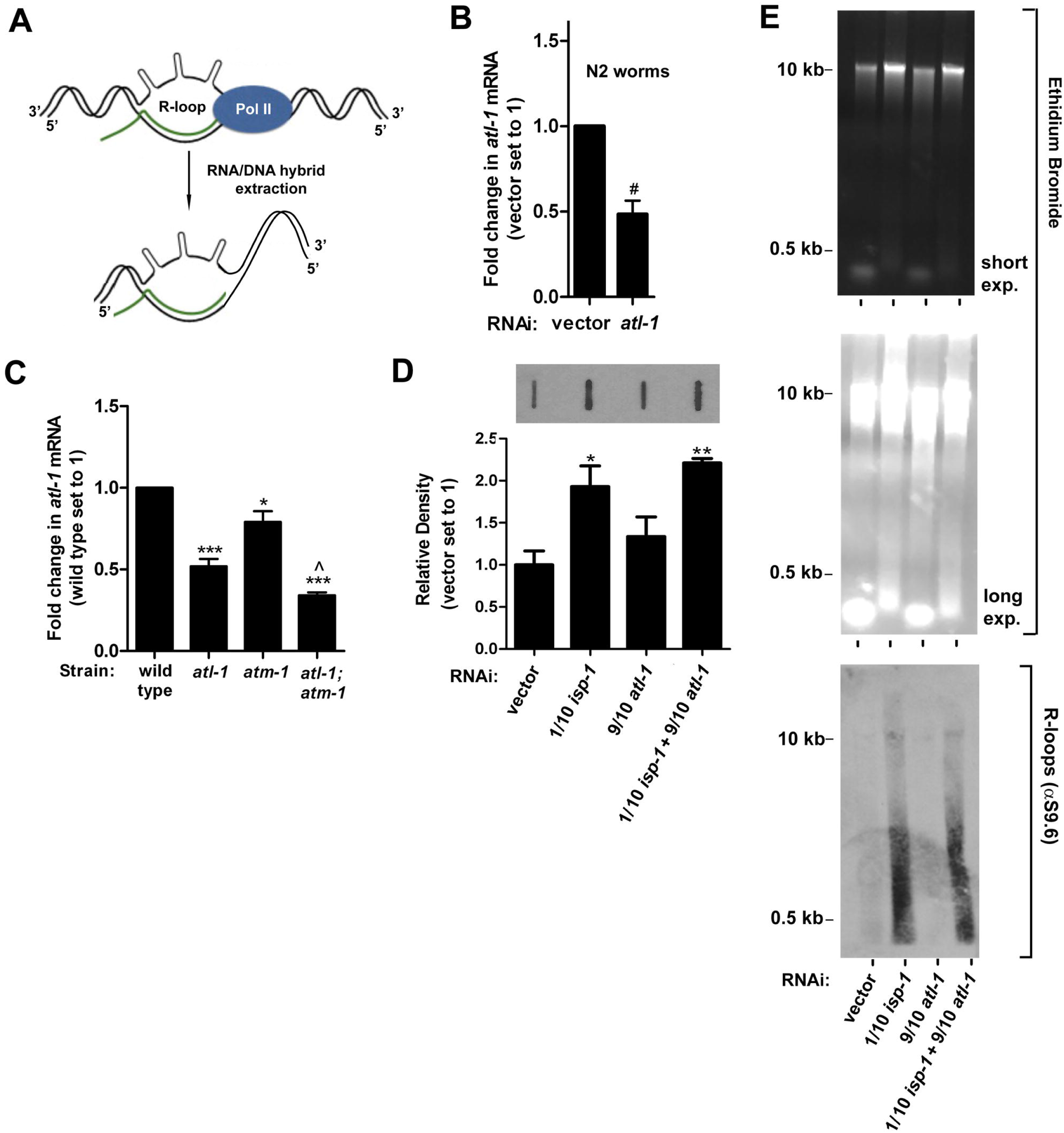
Evidence for RNA Pol II stalling following mitochondrial ETC stress. (**A**) Schematic showing genesis of R-loops following stalling of RNA Polymerase II. Modified from ^51^. (**B, C**) RNAi knockdown of *atl-1* in wild type (N2) worms reduces *atl-1* transcript abundance to the same extent as in maternal-effect *atl-1(tm853)* mutants. In (C), all worms are unbalanced F1 progeny from parents carrying the nT1 reciprocal chromosomal translocation (*N = 3*; *error bars*: SEM; *significance testing*: Student’s t-test; *symbols:* significantly different from wt: * p<0.05; *** p<0.001; significantly different from *atl-1(tm853): ^ p < 0.05)*. In (B), *N=2* and *#* marks *range.* (**D**) R-loops are significantly increased in N2 worms following knockdown of *isp-1* (1/10^th^ RNAi strength). Combined knockdown of *isp-1* (1/10^th^ RNAi strength) and *atl-1* (9/10^th^ RNAi strength) does not abrogate R-loop accumulation. *Top panel:* representative immunoblot of R-loops, *Bottom panel:* Immunoblot quantitation: *N = 3; error bars*: SEM; Student’s t-test (vs. vector): * p≤0.05, ** p≤0.01. (**E**) Samples in (D) were further analyzed by agarose gel electrophoresis (*top two panels)* and then western blotting using an antibody that cross reacts with DNA::RNA hybrids (αS9.6, *bottom panel*). Knockdown of *isp-1* results in an increase of gDNA associated R-loops, and unknown small molecular weight DNA::RNA hybrids.

### RNA splicing is not altered by mitochondrial ETC disruption

Having identified a possible mechanism by which ATL-1 is activated in worms experiencing mitochondrial ETC stress, we next sought to determine the mode by which ATL-1 acts to further disrupt mitochondrial function. It is well-established that pre-mRNA splicing is a co-transcriptional process ^52^ and it has also been reported that *isp-1(qm150)* mutants are sensitive to knockdown of two spliceosome factors SFA-1 and REPO-1, fully ablating their life extension ^53^. We therefore tested whether stalling of RNA pol II altered either splicing efficiency or splicing fidelity in worms exposed to ETC disruption, and if ATL-1 played a role in this process. We used two synthetic-gene reporter assays developed by Kuroyanagi and colleagues ^54, 55^ designed to measure alternate splicing at the *egl-15* and *ret-1* loci, respectively. The latter reporter was shown to be a sensitive marker of age-related splicing dysfunction ^53^. Each assay has the potential to produce two fluorescent outputs that record changes in the nature of a differential splicing event and the tissues in which these events occur (**Supplemental Fig. S4 A, B**). We exposed worms carrying the relevant reporter cassette to double feeding RNAi targeting *atp-3* or *isp-1* (1/10^th^ strength) in conjunction with either *atl-1* or vector control (9/10^th^ RNAi strength) (**Supplemental Fig. S4C**). Animals were followed three days into adulthood. None of the conditions that disrupted the mitochondrial ETC, neither in the presence or absence of *atl-1,* resulted in alteration of *egl-15* or *ret-1* splicing relative to vector control treated animals.

### Reduced ATL-1 activity does not enhance hormetic stress response activation

We next tested whether loss of *atl-1* affected the activity of major stress response pathways known to be activated by *atp-3* or *isp-1* depletion ^3, 13, 35, 56^. Loss of *atl-1* did not result in constitutive activation of P*hsp-6::GFP*, P*gst-4::GFP* or P*tbb-6::GFP*, which are markers of ATFS-1, SKN-1/NRF-2 and PMK-3/p38 activation, respectively (**Supplemental Fig. S5A-D**). Neither did loss of *atl-1* increase gene expression of *sod-3, ugt-61* or *hsp-6* (**Supplemental Fig. S5E**). These results indicate that reduction of *atl-1* does not affect typical mitochondrial stress response pathways.

### Oxygen consumption remains surprisingly unaltered in *atl-1* mutants exposed to ETC disruption

We next tested whether mitochondria in *atl-1(tm853)* mutants were inherently different from those of wild type worms. We also analyzed mitochondria from *atm-1(gk186)* single mutant worms, as well as *atl-1(tm853); atm-1(gk186)* double mutants. Properties that we examined included mitochondrial DNA (mtDNA) content, nuclear-encoded ETC transcript abundance, mitochondrial morphology and whole-worm oxygen consumption. Our findings can be summarized as follows: relative to wild type worms, mtDNA copy number is halved in *atl-1(tm853)* mutants, consistent with an earlier report ^57^. mtDNA content was also halved in *atl-1(tm853); atm-1(gk186)* double mutants (Fig. 7A). To assess changes in nuclear-encoded ETC transcript abundance, we focused on five genes: *nuo-6, mev-1, isp-1, cco-1* and *atp-3*, representing subunits from mitochondrial complexes I through V, respectively. We found that all transcripts except *nuo-6* were significantly decreased relative to wild type worms in all four mutant strains studied. The only exception was *mev-1,* for which mRNA levels remained unchanged in *atm*-*1(gk186)* mutants (Fig. 7B).

**Figure 7.**
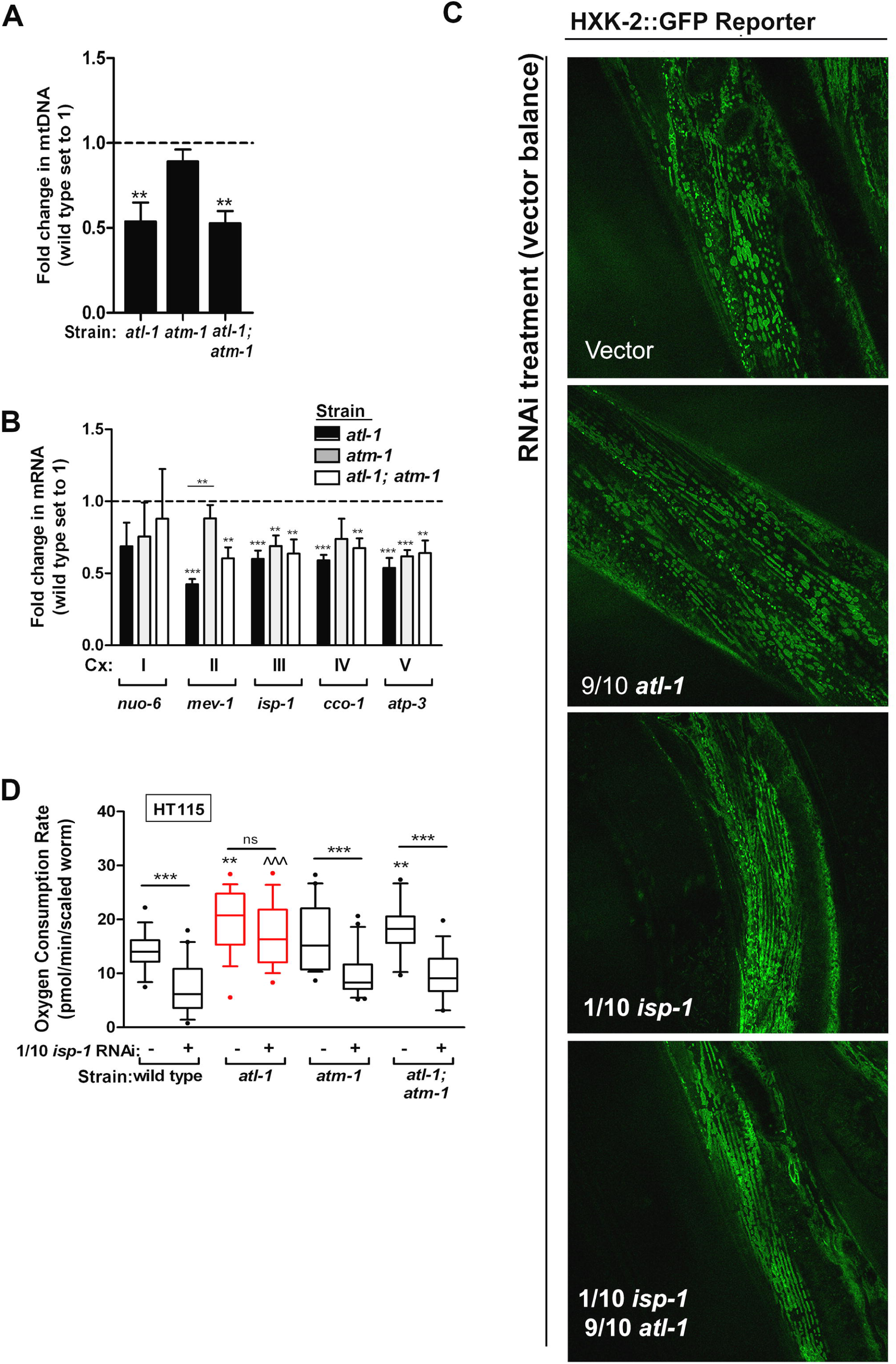
*atl-1(tm853)* mutants maintain normal oxygen consumption when confronted with disruption of their mitochondrial ETC. (**A**) Mitochondrial DNA abundance is significantly decreased in *atl-1(tm853)* mutants compared to wild type worms (N=3). (**B**) Transcript abundance of nuclear-encoded ETC genes is significantly decreased in *atl*-*1(tm853)* mutants relative to wild type worms (N=3; Cx: complex). (**C**) Morphology of body-wall muscle mitochondria assessed using HXK-2::GFP ^58^, following *atl-1* knockdown in the absence or presence of mitochondrial ETC disruption (1/10^th^ strength *isp-1* RNAi). Both *atl-1* and *isp-1* knockdown significantly alter mitochondrial morphology relative to vector-control (p<0.0024, refer to Methods for significance testing). (**D**) *atl-1(tm853)* mutants are resistant to changes in oxygen consumption following *isp-1* knockdown (1/10^th^ strength RNAi). All other tested strains experienced a significant decrease in oxygen consumption (N=4). Oxygen consumption was measured using a Seahorse XFe24 Analyzer. In (A) and (D), all worms are the unbalanced F1 progeny of parents carrying the nT1 reciprocal chromosomal translocation. In (A), (B) and (D), asterisks indicate significantly different from wild type (Student’s t-test) unless otherwise noted by a bar for the relevant comparison group: * p<0.05. ** p<0.01. *** p<0.001; ns, not significant. In (D), ^^^ p<0.001 in comparison to wild type on *isp-1* RNAi. In (A) and (B), error bars are SEM; in (D), box and whisker plots show 10-90 percentile, with outliers marked (dots).

To quantify changes in mitochondrial morphology we employed a hexokinase-2::GFP translational reporter (HXK2::GFP) that is localized to the outer mitochondrial membrane ^58^. We observed a significant increase in the number of fused mitochondria in reporter worms treated with *isp-1* RNAi (p<0.0024, see **Methods** for details of statistical testing). Co-knockdown of *atl-1* did not significantly affect this result. Mitochondria in worms treated only with *atl-1* RNAi were indistinguishable from those of wild type animals (Fig. 7C). Finally, the most significant finding that we uncovered related to alterations in oxygen consumption by *atl-1(tm853)* mutants. When wild type worms, *atl-1(tm853)* and *atm-1(gk186)* single mutants, as well as *atl-1(tm853); atm-1(gk186)* double mutants were cultured on *isp-1* RNAi, the basal oxygen consumption rate of *atl-1(tm853)* mutants, as well as *atl-1(tm853); atm*-*1(gk186)* double mutants, significantly increased relative to wild type worms (Fig. 7D). Moreover, while *isp-1* knockdown uniformly decreased oxygen consumption in wild type worms, *atm-1(gk186)* single mutants and *atl-1(tm853); atm-1(gk186)* double mutants, and by the same extent, oxygen consumption in *atl-1(tm853)* surprisingly remained unchanged following this treatment. We conclude that while *atl*-*1(tm853)* mutants contain half as much mtDNA as wild type worms and have reduced abundance of several nuclear-encoded ETC transcripts, these worms surprisingly maintain normal oxygen consumption, even when confronted with disruption of their mitochondrial electron transport chain.

### Loss of ATR leads to recovery of translational activity in worms exposed to ETC disruption

Several mechanisms could explain how *atl-1(tm853)* mutants maintain oxygen consumption despite ETC disruption. Any process that ultimately increases flux or efficiency of the respiratory chain could be involved. Enhanced supercomplex formation, elevated matrix Ca^2+^, modification of ETC subunits, or simply enhanced translational output of respiratory subunits, could all be behind the unique respiratory response seen in *atl-1(tm853)* mutants. We focused on translational output because there is evidence from human 293T cells that ATR might directly regulate the ribosomal machinery by affecting phosphorylation of the nascent polypeptide-associated complex (NAC) ^22^. In worms, ICD-1 is the sole βNAC isoform ^59, 60^. Although the ATR phosphorylation site of human BTF3/βNAC is not conserved in ICD-1, we observed by western analysis that ICD-1 protein levels are selectively elevated in *atl*-*1(tm853)* mutants following *atp-3* knockdown (*isp-1* knockdown was not tested due to limited quantities of antisera, see Materials & Methods) (Fig. 8A). Moreover, early analyses involving several hundred microarray datasets ^61^ showed *icd-1* formed part of a tightly clustered gene co-expression group (Fig. 8B) that is significantly enriched (hypergeometric distribution, p<0.05) for genes encoding ribosomal and mitochondrial ETC subunits, as well as key translational regulatory factors (Fig. 8C). We therefore tested whether global translational activity was downregulated by mitochondrial ETC disruption and if loss of *atl-1* permitted its recovery. We employed polysome profiling in conjunction with RNAi targeting both *isp-1* (1/10^th^ RNAi strength) and either *atl-1* or vector (both at 9/10^th^ RNAi strength). We found that the actively translating polysome fraction of worms treated with *isp-1* RNAi was decreased by half relative to vector control-treated worms. Strikingly, simultaneous removal of both *atl-1* and *isp-1* restored polysome activity to wild type levels (Fig. 8D). Loss of *atl-1* in the absence of ETC disruption did not significantly impact polysome abundance relative to wild type worms. These data imply that ATL-1, when activated by mitochondrial ETC disruption, normally acts to dampen global ribosomal translation. Whether this is through a novel phosphorylation site on ICD-1, downregulation of factors that normally control *icd-1* expression, or via some other means that globally regulates translational activity, remains unclear. There must be some specificity in the translational targets of ATR-1, however, because simply elevating global rRNA production by inhibiting *ncl-1*, a negative regulator of FIB-1/fibrillarin, a nucleolar protein involved in the regulation and maturation of rRNA, is not sufficient to block the size reduction caused by *isp-1* bacterial feeding RNAi (Fig. 6E, F). Collectively, our results lead to a model for how ATL-1 acts to modulate the mitochondrial threshold effect (Fig. 8G).

**Figure 8.**
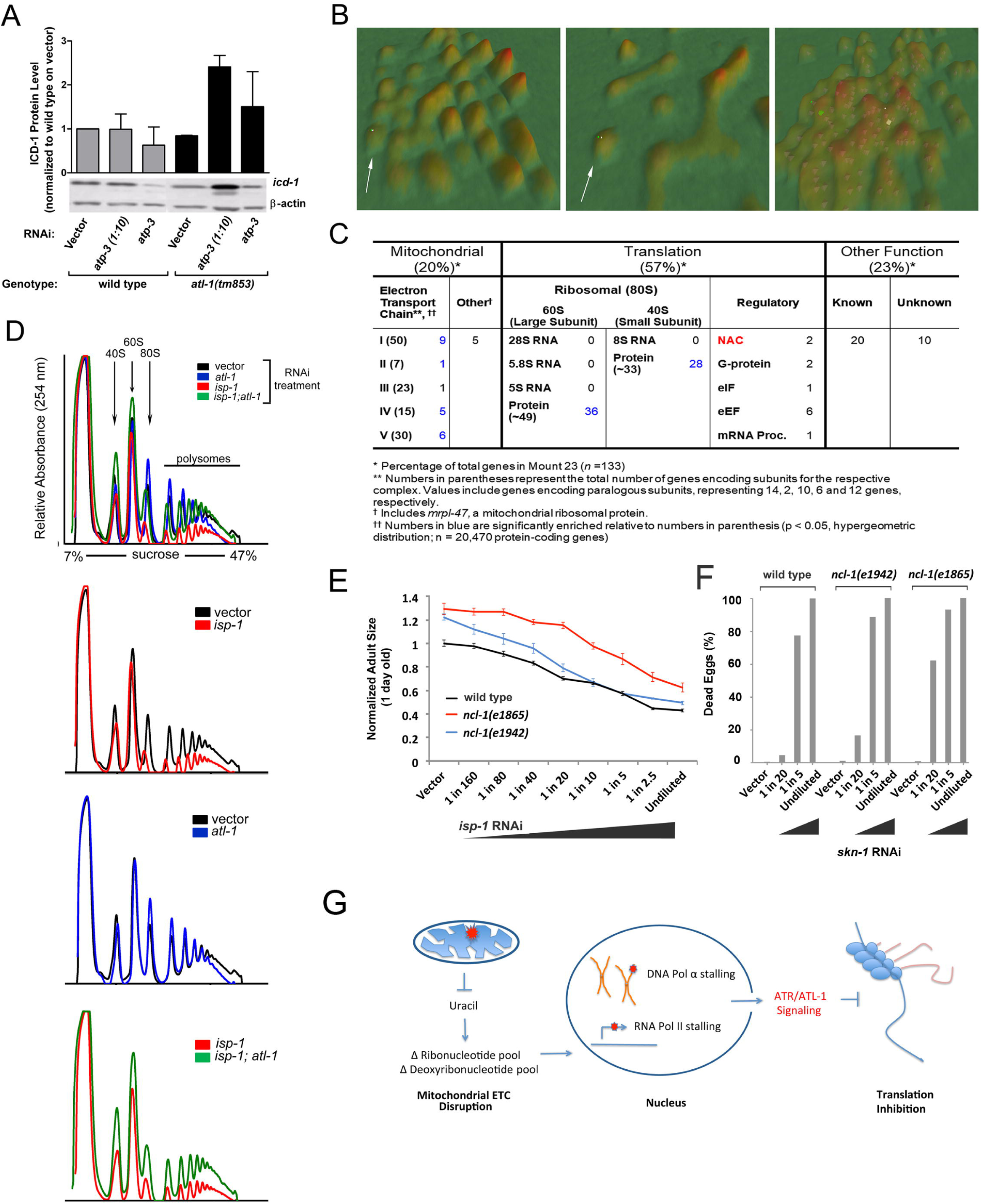
Knockdown of *atl-1* prevents global translation reduction caused by mitochondrial ETC disruption. (**A**) *Top panel:* ICD-1 protein abundance in *atl-1(tm853)* mutants relative to wild type animals following *atp-3* knockdown (N=2, bars indicate *range*). *Bottom panel:* Representative western blot. All worms are the unbalanced F1 progeny of parents carrying the nT1 reciprocal chromosomal translocation. (**B**) *C. elegans* gene co-expression terrain map generated using VxInsight ^61^ showing *icd-1-*containing ‘Mount 23’ *(white arrow).* Progressive zoom shots of Mount 23 are shown (*top to bottom*). The position of *icd-1* is marked by a *white square*. (**C**) 88 of 133 (66%) genes comprising Mount 23 of the *C. elegans* gene co-expression terrain map encode either mitochondrial ETC proteins or ribosomal proteins, including both α- and β-NAC (ICD-1). (**D**) Polysome profiling shows knockdown of *atl-1* (9/10^th^ RNAi strength) restores the level of actively translating ribosomes to control levels in worms grown on *isp-1* RNAi (1/10^th^ strength). (**E**) *ncl-1(e1865) and ncl-1(e1942)* mutants do not confer resistance to the size reducing effects of bacterial feeding RNAi targeting *isp-1* relative to wild type (N2) worms. *ncl-1* mutants are naturally larger that N2 worms. (mean +/- SEM) (**F**) RNAi remains efficacious in *ncl-1(e1865)* and *ncl-1(e1942)* mutants. Shown is survival following exposure to increasing doses of a lethal feeding RNAi (*skn-1*). Data from a single experiment is shown that was established simultaneously with common reagents and using 20 synchronous one day old adult worms per test condition. (**G**) Model showing how removal of ATL-1 counters loss of mitochondrial electron transport chain disruption. ETC insufficiency results in nucleotide imbalances, stalling of nucleotide polymerases, activation of ATL-1, and dampening of global translation, leading to magnification of ETC dysfunction.

## DISCUSSION

In this study, we discovered that the PIKK kinase ATR/ATL-1 plays a specific role in modulating the mitochondrial threshold effect of *C. elegans*. When flux through the ETC was reduced, ATL-1 was activated. We presented data suggesting stalling of RNA polymerase could be the trigger for ATL-1 activation. We recorded measurable reductions in many nucleotide species in adult worms, including substrates for both RNA and DNA polymerase, and we presented biochemical evidence showing RNA polymerase stalling following ETC disruption. In addition to these findings we showed that ATL-1 paradoxically exacerbates mitochondrial dysfunction during times of mitochondrial stress. Specifically, when worms containing reduced levels of *atl-1* were challenged with ETC disruption, unlike wild type animals, these showed no signs of impairment in global translational output or respiratory oxygen consumption. Phenotypically, loss of *atl-1* effectively reversed a mitochondrial ETC insufficiency in *C. elegans*. The survival benefits of further dampening ETC activity following an ETC insult are not entirely clear, but dampening of mitochondrial respiratory activity in conjunction with global translational dampening might represent an effort to slow overall metabolism or to maintain stoichiometric balance between nuclear- and mitochondria-encoded ETC proteins ^62^.

### Mode of ATL-1 Activation Following Mitochondrial ETC Disruption

In our search for signals that triggered ATL-1 activation in response to ETC disruption we found no evidence for elevated rates of nuclear DNA mutation or strand breakage. This does not preclude the possibility that DNA damage occurred and was accurately repaired. Such damage would have escaped detection in our assays. Nonetheless, we investigated other potential mechanisms of ATL-1 activation and found that nucleotide pools were significantly changed when analyzing *isp-1(qm150)* mutants. This is significant because ATR activation has been linked to changes in nucleotide pools in other model systems. Specifically, in *S. cerevisiae,* glucose starvation induces Snf1p (AMPK) at the mitochondrial surface following reduction of adenosine ribonucleotide energy charge. In turn, Snf1p was shown to phosphorylate Mec1p (ATR) ^34^. Whether a similar mode of ATR activation functions in *C. elegans* remains unclear, but Ggc1p, the protein that initially recruits Mec1p to the yeast mitochondrial membrane under low glucose conditions has no worm ortholog. We therefore decided to pursue an alternate hypothesis for nucleotide-dependent activation of ATR in worms, namely stalling of RNA and/or DNA polymerases.

Previously it had been shown that blocking RNA polymerase during transcriptional elongation is sufficient to activate ATR and induce a DNA damage response (DDR) ^48^. Indeed, we found evidence that DNA::RNA R-loop hybrids are increased in the presence of mitochondrial ETC disruption, which is indicative of transcriptional stalling. However, it is unclear whether all the R-loops we detected were genomic DNA (gDNA)-derived transcription intermediates. When examining samples on an agarose gel in conjunction with an antibody targeting these structures (S9.6), R-loops are expected to appear at a high molecular weight in association with the gDNA band. We clearly detected an elevated signal in this region in our *isp-1* RNAi treated samples, consistent with our hypothesis. Also but, we observed an additional smear of low molecular weight nucleic acids. At this point it is unknown if these low molecular weight molecules are truly DNA::RNA hybrids or a contaminating nucleic acid. The S9.6 antibody has a 5-fold higher affinity for DNA::RNA hybrids over RNA::RNA hybrids ^50^, so in anticipation of the latter our samples were treated with RNase A ruling out this possibility. One idea is that the low molecular weight DNA::RNA hybrids are derived from mtDNA fragments that have leaked from dysfunctional mitochondria. Consistent with this idea, in mouse embryonic fibroblasts, mitochondrial stress results in mtDNA leaking into the cytoplasm and initiating an immune response ^63^. On the other hand, worms use a rolling circle replication method for their mtDNA, so precisely how R-loops would form in the absence of D-loop (theta) replication, as is used by mammalian cells, remains enigmatic ^64^. Another idea is that the low molecular weight species we detected are cytoplasmic DNA::RNA hybrids produced by DNA polymerase α (DNA pol α). Starokadomsky and colleagues ^65^, showed that a small fraction of DNA pol α localizes to the cytoplasm in human cells where it generates DNA::RNA hybrids. These molecules directly modulate production of type I interferons and possibly exist to control the activation threshold of the innate immune response by foreign nucleic acids. In patients with X-linked reticulate pigmentary disorder (XLPDR), a primary immunodeficiency with autoinflammatory features, levels of the catalytic subunit of DNA pol α (DNA PolA1) are decreased enough to disrupt cytoplasmic DNA::RNA hybrid formation, but not enough to affect DNA replication ^65^. Intriguingly, in our screen for DDR proteins that when inhibited reversed the small size phenotype of *atl-1(tm853)*; *atm-1(gk186)* double mutants experiencing mitochondrial ETC stress, we identified an RNAi clone targeting DNA PolA1. Although not discussed earlier, we also isolated a second DNA pol α clone in that screen that targeted a different region of DNA PolA1 but instead resulted in synthetic lethality. This dose-dependent phenotypic response is remarkably similar to what was just discussed for humans. Obviously, more work is needed to determine the identity, provenance and significance of the low molecular weight DNA::RNA species in worms experiencing ETC stress and if and how they might relate to ATR activation.

### Downstream targets of ATL-1 that Modulate Mitochondrial ETC Activity

One important finding from our study is the observation that *atl-1* knockdown mitigates the effects of mitochondrial ETC dysfunction. This is an intriguing finding as it suggests the normal cellular response to ETC disruption is ostensibly further detrimental. Two possible explanations for how loss of *atl-1* could result in enhanced ETC function are, *atl-1* knockdown triggers active restoration of mitochondrial function, or alternatively, *atl-1* knockdown precludes propagation of whatever negative downstream effects are normally elicited by ETC disruption. According to our data, *atl-1(tm853)* mutants contain half the amount of mtDNA as wild type worms and have significantly reduced abundance of nuclear-encoded ETC transcripts. These findings argue against the first hypothesis. Surprisingly, despite their shortcomings, we find that *atl-1(tm853)* mutants are still able to maintain oxygen consumption at near normal levels when confronted with ETC disruption. The precise mechanism involved is yet to be determined, but it may be as simple as *atl-1(tm853)* mutants being able to maintain the translation of a factor that enhances supercomplex formation, for example ^66^. Alternatively *atl-1(tm853)* mutants may be able to keep their cells fortified with sufficient ETC proteins to counteract any rapid turnover that is likely to occur when subunits become rate-limiting during the assembly process. Along these lines, when we examined polysome formation under conditions of ETC dysfunction, we found that animals with functional ATL-1 had reduced polysome formation but removal of *atl-1* restored this number back to pre-ETC disruption levels. Future studies will be aimed at determining the precise mRNA species present in these polysomes.

Another way in which *atl-1(tm853)* mutants might be able to counter ETC disruption involves the nascent protein folding machinery. We showed that ICD-1/βNAC, a protein that acts as a ribosome-associated chaperone, is upregulated in *atl-1* mutants upon ETC disruption. In yeast and plants, βNAC plays a role in targeting proteins to the mitochondria and is required for efficient mitochondrial function ^67,68,69,70,71^. βNAC was also one of the few proteins found to be specifically regulated by ATR-dependent phosphorylation in mammals ^22^. The ATR-dependent phosphorylation site in mammals is not conserved in *C. elegans*, however there are several other potential phosphorylation sites in the ICD-1 protein sequence ^72^. It will be of particular interest in future to ascertain if ATR directly regulates βNAC abundance on ribosomes to modulate translation of ETC transcripts in worms.

### Mitochondrial Dysfunction in Mammals

Johnson and colleagues ^73^ successfully extended the survival of a mouse model of Leigh syndrome, which was deficient in Ndufs4 of complex I, by chronic administration of the mTOR inhibitor rapamycin. mTOR is a positive regulator of p70S6 kinase and a negative regulator of 4E-BP, both of which work to elevate translation. At first this seems surprising in light of our findings suggesting global translation needs to be elevated to counter ETC dysfunction in worms. However, while acute inhibition of mTOR indeed results in global dampening of translation, it also results in the specific translational upregulation of mitochondrial ETC components ^74, 75^. Also, it has been reported that in contrast to acute rapamycin administration, chronic rapamycin administration does not lead to reduced ribosomal activity *in vivo* ^76^. Whether the increase in polysomes that we observed in *atl-1* animals exposed to ETC disruption represented upregulation of global mRNA translation or just a discrete subset of mRNAs only encoding ETC proteins, remains to be determined.

In conclusion, we have shown that blockade of *atl-1* effectively reverses ETC disruption in *C. elegans*. Preventing mitochondria from transitioning into their critical ETC threshold appears, quite literally, to be the difference between life and death.

## METHODS

A detailed description of all methods is provided in **Supplemental File S1**.

### Nematode Strains and Maintenance

Genotypic information for all strains used in this study is provided in **Table S4**. Lines were maintained at 20°C on standard NGM agar plates seeded with *E. coli* (OP50) ^77^. To control for possible untoward effects derived from the *nT1(IV:V)* chromosomal pair in maternally-rescued *atl-1(tm853)* mutants, all lines, including the wild type control, were moved into the *nT1(IV:V)* background and then unbalanced F1 animals freshly collected for each relevant experiment. Further details are provided in **Supplemental File S1**.

### Bacterial Feeding RNAi

All bacterial feeding RNAi constructs were designed in the pL4440 vector backbone and maintained in HT115 bacteria ^78^. Growth rate, life span, RNAi efficacy assays and RNAi dilution methods are described in **Supplemental File S1**.

### Microscopy

Images of fluorescent worms were captured using an Olympus SZX16 fluorescence microscope connected to an Olympus DP71 CCD camera. In Fig. 7C, to determine whether differences existed in the morphology of mitochondria between the four tested conditions, four randomly selected images from each condition were mixed and then grouped by a scorer who was blind to image identity. In a set of 16 images, 10 were co-grouped correctly. The probability that this grouping occurred by chance was modeled using an Excel VBA macro that randomly binned the images into four groups of four, 10,000 times (**File S3**). The final *p-value* was <0.0024 and deemed significant.

### mRNA and mtDNA Quantitation

For quantitation of mRNA, fold change in each mRNA of interest was determined using the ΔΔCt method normalized to the geometric mean of *cdc-42*, *pmp-3*, and *Y45F10D.4* ^79^. For quantitation of mitochondrial DNA (mtDNA), PCR primer pairs targeting mtDNA-encoded *ctb-1* and intron 4 of genomic DNA-encoded *ama-1* were used as described ^80^. The ratio of mitochondrial DNA to nuclear DNA was then calculated using Real Time PCR, using *ama-1* as the normalizing quantity ^81^. A complete list of qPCR primers is provided in **Table S5**.

### Mutation Screening Assays

Full methodological details of the *unc-93(e1500)* and *unc-58(e665)* reversion assays, the eT1(III;V) and nT1(IV:V) lethal mutation assays, the Lac-Z Frameshift Assay, and the ‘terminal deoxynucleotidyl transferase dUTP nick end labeling’ (TUNEL) assay are presented in **Supplemental File S1**.

#### Nucleotide Extraction and Mass Spectrometry

Quantitation of deoxyribonucleotide and ribonucleotide species in whole-worm extracts of *isp-1(qm150)* and wild-type (N2) animals was undertaken using HPLC-MS at the UTHSCA Mass Spectrometry Facility. Methods for nucleotide extraction, separation, quantitation and statistical analysis are provided in **Supplemental File S1**.

#### R-loop Quantification

R-loop formation in worms was quantified using αS9.6 primary antibody (Kerafast, 1:2000 in 5% Blotto, 16 hours, 4°C), previously shown to be selective for DNA::RNA hybrids ^50^. A list of western reagents and methods are provided in **Supplemental File S1**.

#### Spliceosome Reporter Assays

*egl-15* and *ret-1* alternate splicing reporter genes were engineered by Kuroyanagi and colleagues ^55, 82^. Details of reporter construction and their use is provided in **Supplemental File S1**.

### Nematode Oxygen Consumption

Nematode oxygen consumption measurements were undertaken using a Seahorse XFe24 Analyzer Approximately 30 worms were added to each Seahorse well, with 5 replicate wells employed per test condition. We followed the procedure of Luz and colleagues ^83^, and only averaged oxygen consumption rates across the final four measurement phases for each well to provide a single rate value for that well. A total of four independent experiments was performed. Significance testing was undertaken using the Student’s t-test, with *p<0.05* considered significant.

#### Polysome Profiling

Quantitative polysome profiling was undertaken using the procedure of Steffen and colleagues ^84^ using a Brandel BR-188 Density Gradient Fractionation System. Several modifications were included to optimize for *C. elegans* polysome extraction and these are described in detail in **Supplemental File S1**. Two fully independent experimental replicates were collected.

#### Western Analysis

Western blotting reagents and suppliers are described in **Supplemental File S1**.

#### DNA Damage Response (DDR) Screen

The *atm-1(gk186); atl-1(tm853)* suppressor screen was undertaken using a bacterial feeding RNAi targeting 201 DDR-related genes ^85^. A detailed protocol of our screening method, including quantitation of the RNAi Effect Size and Significance Testing, is provided in **Supplemental File S1** and **Tables S1-S3**).

## FUNDING

Financial support was provided by the National Institutes of Health [AG-047561 (SLR), AG025207 (SLR) and AG-055820 (SLR). Funding sources had no role in study design, data collection and analysis, the decision to publish, or preparation of the manuscript.

## ACKNOWLEDGEMENTS

We thank the following people: Dr. Hidehito Kuroyanagi (Tokyo Medical and Dental University) for the splice reporters; Dr. Joel Rothman (UCSB) for the *icd-1* antibody; Alison Kell (CU, Boulder) for assistance with DDR library construction; Oxana Radetskaya (UW) for critical reading of the manuscript. Technical assistance was provided by Rebecca Lane, Carlos Galeano, Yvonne Penrod, Haley Beam, Bill Swan and Melissa Ligon (all UT Health San Antonio). Some strains were provided by the *Caenorhabditis* Genetic Stock Center, which is funded by NIH Office of Research Infrastructure Programs (P40 OD010440). We also thank Wormbase.

## SUPPLEMENTAL MATERIAL

### SUPPLEMENTAL FIGURES

**Figure S1.** Loss of ATL-1 desensitizes worms to mitochondrial respiratory chain stress induced by ethidium bromide.

**Figure S2.** DNA mutation detection in *C. elegans* using β-Gal frame-shift- and TUNEL assays.

**Figure S3.** Nucleotide biosynthesis in *C. elegans* highlighting the role of the mitochondrial ETC.

**Figure S4.** mRNA splicing remains unaltered in worms experiencing ETC disruption, both in the absence or presence of *atl-1* knockdown.

**Figure S5.** *atl-1* knockdown does not alter retrograde response pathways normally induced by mitochondrial ETC stress.

### SUPPLEMENTAL TABLES

**Table S1.** Summary of descriptive statistics for final-round DDR test RNAi screen hits. (Related to Fig. 3).

**Table S2.** Mean Ranks of final-round DDR RNAi hits used for Kruskal-Wallis test. (Related to Fig. 3).

**Table S3.** Post-hoc tests for each final-round DDR RNAi hit versus vector. (Related to Fig. 3).

**Table S4.** List of *C. elegans* strains employed in current study.

**Table S5.** qPCR primer list.

### OTHER SUPPLEMENTAL FILES

**File S1.** Supplemental Methods & Supplemental Figure Legends

**File S2.** Mean difference ratios of all hits from DNA Damage Response (DDR) screen. (Related to Fig. 3)

**File S3.** Significance testing for differences in HXK2::GFP fluorescence. (related to Fig. 7C)

## Notes

#### Summary of Updates

Added missing Supplemental Figure Files S1-S5. Amended label on Figure 6D

## REFERENCES

1. Schaefer AM, Taylor RW, Turnbull DM, Chinnery PF. The epidemiology of mitochondrial disorders--past, present and future. Biochim Biophys Acta 2004, 1659(2-3): 115–120.

2. Li MX, Dewson G. Mitochondria and apoptosis: emerging concepts. F1000Prime Rep 2015, 7: 42.

3. Liu Y, Samuel BS, Breen PC, Ruvkun G. Caenorhabditis elegans pathways that surveil and defend mitochondria. Nature 2014, 508(7496): 406–410.

4. Khutornenko AA, Roudko VV, Chernyak BV, Vartapetian AB, Chumakov PM, Evstafieva AG. Pyrimidine biosynthesis links mitochondrial respiration to the p53 pathway. Proc Natl Acad Sci U S A 2010, 107(29): 12828–12833.

5. Wallace DC. Mitochondrial DNA mutations in disease and aging. Environ Mol Mutagen 2010, 51(5): 440–450.

6. Wallace DC. Mitochondria and cancer. Nat Rev Cancer 2012, 12(10): 685–698.

7. Lane RK, Hilsabeck T, Rea SL. The role of mitochondrial dysfunction in age-related diseases. Biochim Biophys Acta 2015.

8. Rossignol R, Faustin B, Rocher C, Malgat M, Mazat JP, Letellier T. Mitochondrial threshold effects. Biochem J 2003, 370(Pt 3): 751–762.

9. Munkácsy E, Rea SL. The paradox of mitochondrial dysfunction and extended longevity. Exp Gerontol 2014, 56(0): 221–233.

10. Arnould T, Michel S, Renard P. Mitochondria Retrograde Signaling and the UPR(mt): Where Are We in Mammals? Int J Mol Sci 2015, 16(8): 18224–18251.

11. Nargund AM, Pellegrino MW, Fiorese CJ, Baker BM, Haynes CM. Mitochondrial import efficiency of ATFS-1 regulates mitochondrial UPR activation. Science 2012, 337(6094): 587–590.

12. Biswas G, Adebanjo OA, Freedman BD, Anandatheerthavarada HK, Vijayasarathy C, Zaidi M, et al. Retrograde Ca2+ signaling in C2C12 skeletal myocytes in response to mitochondrial genetic and metabolic stress: a novel mode of inter-organelle crosstalk. The EMBO Journal 1999, 18(3): 522–533.

13. Munkacsy E, Khan MH, Lane RK, Borror MB, Park JH, Bokov AF, et al. DLK-1, SEK-3 and PMK-3 Are Required for the Life Extension Induced by Mitochondrial Bioenergetic Disruption in C. elegans. PLoS Genet 2016, 12(7): e1006133.

14. Heo JM, Livnat-Levanon N, Taylor EB, Jones KT, Dephoure N, Ring J, et al. A stress-responsive system for mitochondrial protein degradation. Mol Cell 2010, 40(3): 465–480.

15. Quiros PM, Mottis A, Auwerx J. Mitonuclear communication in homeostasis and stress. Nat Rev Mol Cell Biol 2016.

16. Schiavi A, Maglioni S, Palikaras K, Shaik A, Strappazzon F, Brinkmann V, et al. Iron-Starvation-Induced Mitophagy Mediates Lifespan Extension upon Mitochondrial Stress in C. elegans. Curr Biol 2015, 25(14): 1810–1822.

17. Jiang P, Jin X, Peng Y, Wang M, Liu H, Liu X, et al. The exome sequencing identified the mutation in YARS2 encoding the mitochondrial tyrosyl-tRNA synthetase as a nuclear modifier for the phenotypic manifestation of Leber’s hereditary optic neuropathy-associated mitochondrial DNA mutation. Hum Mol Genet 2016, 25(3): 584–596.

18. Huyen Y, Jeffrey PD, Derry WB, Rothman JH, Pavletich NP, Stavridi ES, et al. Structural Differences in the DNA Binding Domains of Human p53 and Its C. elegans Ortholog Cep-1. Structure 2004, 12(7): 1237–1243.

19. Ventura N, Rea SL, Schiavi A, Torgovnick A, Testi R, Johnson TE. p53/CEP-1 increases or decreases lifespan, depending on level of mitochondrial bioenergetic stress. Aging Cell 2009, 8(4): 380–393.

20. Schiavi A, Torgovnick A, Kell A, Megalou E, Castelein N, Guccini I, et al. Autophagy induction extends lifespan and reduces lipid content in response to frataxin silencing in C. elegans. Experimental Gerontology 2013, 48(2): 191–201.

21. Lovejoy CA, Cortez D. Common mechanisms of PIKK regulation. DNA Repair (Amst) 2009, 8(9): 1004–1008.

22. Matsuoka S, Ballif BA, Smogorzewska A, McDonald ER, III, Hurov KE, Luo J, et al. ATM and ATR Substrate Analysis Reveals Extensive Protein Networks Responsive to DNA Damage. Science 2007, 316(5828): 1160–1166.

23. Marechal A, Zou L. DNA damage sensing by the ATM and ATR kinases. Cold Spring Harb Perspect Biol 2013, 5(9).

24. Vousden KH, Lane DP. p53 in health and disease. Nat Rev Mol Cell Biol 2007, 8(4): 275–283.

25. Yang DQ, Kastan MB. Participation of ATM in insulin signalling through phosphorylation of eIF-4E-binding protein 1. Nat Cell Biol 2000, 2(12): 893–898.

26. Schneider JG, Finck BN, Ren J, Standley KN, Takagi M, Maclean KH, et al. ATM-dependent suppression of stress signaling reduces vascular disease in metabolic syndrome. Cell Metab 2006, 4(5): 377–389.

27. Cosentino C, Grieco D, Costanzo V. ATM activates the pentose phosphate pathway promoting anti-oxidant defence and DNA repair. EMBO J 2011, 30(3): 546–555.

28. Schroeder EA, Raimundo N, Shadel GS. Epigenetic silencing mediates mitochondria stress-induced longevity. Cell Metabolism 2013, 17(6): 954–964.

29. Fang EF, Bohr VA. NAD+: The convergence of DNA repair and mitophagy. Autophagy 2017, 13(2): 442–443.

30. Chiolo I, Minoda A, Colmenares SU, Polyzos A, Costes SV, Karpen GH. Double-strand breaks in heterochromatin move outside of a dynamic HP1a domain to complete recombinational repair. Cell 2011, 144(5): 732–744.

31. Gupta A, Sharma S, Reichenbach P, Marjavaara L, Nilsson AK, Lingner J, et al. Telomere length homeostasis responds to changes in intracellular dNTP pools. Genetics 2013, 193(4): 1095–1105.

32. Kumar A, Mazzanti M, Mistrik M, Kosar M, Beznoussenko GV, Mironov AA, et al. ATR mediates a checkpoint at the nuclear envelope in response to mechanical stress. Cell 2014, 158(3): 633–646.

33. Hilton BA, Li Z, Musich PR, Wang H, Cartwright BM, Serrano M, et al. ATR Plays a Direct Antiapoptotic Role at Mitochondria, which Is Regulated by Prolyl Isomerase Pin1. Mol Cell 2015, 60(1): 35–46.

34. Yi C, Tong J, Lu P, Wang Y, Zhang J, Sun C, et al. Formation of a Snf1-Mec1-Atg1 Module on Mitochondria Governs Energy Deprivation-Induced Autophagy by Regulating Mitochondrial Respiration. Dev Cell 2017, 41(1): 59–71 e54.

35. Rea SL, Ventura N, Johnson TE. Relationship Between Mitochondrial Electron Transport Chain Dysfunction, Development, and Life Extension in Caenorhabditis elegans. PLoS Biol 2007, 5(10): e259.

36. Melo JA, Ruvkun G. Inactivation of conserved C. elegans genes engages pathogen- and xenobiotic-associated defenses. Cell 2012, 149(2): 452–466.

37. Bennett CF, Kwon JJ, Chen C, Russell J, Acosta K, Burnaevskiy N, et al. Transaldolase inhibition impairs mitochondrial respiration and induces a starvation-like longevity response in Caenorhabditis elegans. PLoS Genet 2017, 13(3): e1006695.

38. Satsuka A, Mehta K, Laimins L. p38MAPK and MK2 pathways are important for the differentiation-dependent human papillomavirus life cycle. J Virol 2015, 89(3): 1919–1924.

39. Reinhardt HC, Aslanian AS, Lees JA, Yaffe MB. p53-Deficient Cells Rely on ATM- and ATR-Mediated Checkpoint Signaling through the p38MAPK/MK2 Pathway for Survival after DNA Damage. Cancer Cell 2007, 11(2): 175–189.

40. Manke IA, Nguyen A, Lim D, Stewart MQ, Elia AE, Yaffe MB. MAPKAP kinase-2 is a cell cycle checkpoint kinase that regulates the G2/M transition and S phase progression in response to UV irradiation. Mol Cell 2005, 17(1): 37–48.

41. Garcia-Muse T, Boulton SJ. Distinct modes of ATR activation after replication stress and DNA double-strand breaks in Caenorhabditis elegans. Embo J 2005, 24(24): 4345–4355.

42. Dmitrieva NI, Celeste A, Nussenzweig A, Burg MB. Ku86 preserves chromatin integrity in cells adapted to high NaCl. Proc Natl Acad Sci U S A 2005, 102(30): 10730–10735.

43. Desler C, Lykke A, Rasmussen LJ. The effect of mitochondrial dysfunction on cytosolic nucleotide metabolism. J Nucleic Acids 2010, 2010.

44. Kuchta RD, Stengel G. Mechanism and evolution of DNA primases. Biochim Biophys Acta 2010, 1804(5): 1180–1189.

45. Guilliam TA, Keen BA, Brissett NC, Doherty AJ. Primase-polymerases are a functionally diverse superfamily of replication and repair enzymes. Nucleic Acids Res 2015, 43(14): 6651–6664.

46. Zou L, Elledge SJ. Sensing DNA damage through ATRIP recognition of RPA-ssDNA complexes. Science 2003, 300(5625): 1542–1548.

47. Tsang WY, Lemire BD. Mitochondrial Genome Content Is Regulated during Nematode Development. Biochemical and Biophysical Research Communications 2002, 291(1): 8–16.

48. Derheimer FA, O’Hagan HM, Krueger HM, Hanasoge S, Paulsen MT, Ljungman M. RPA and ATR link transcriptional stress to p53 10.1073/pnas.0705317104. PNAS 2007: 0705317104.

49. Hamperl S, Cimprich KA. The contribution of co-transcriptional RNA:DNA hybrid structures to DNA damage and genome instability. DNA Repair (Amst) 2014, 19: 84–94.

50. Zhang ZZ, Pannunzio NR, Hsieh CL, Yu K, Lieber MR. Complexities due to single-stranded RNA during antibody detection of genomic rna:dna hybrids. BMC Res Notes 2015, 8: 127.

51. Sollier J, Cimprich KA. Breaking bad: R-loops and genome integrity. Trends Cell Biol 2015, 25(9): 514–522.

52. Proudfoot NJ, Furger A, Dye MJ. Integrating mRNA processing with transcription. Cell 2002, 108(4): 501–512.

53. Heintz C, Doktor TK, Lanjuin A, Escoubas C, Zhang Y, Weir HJ, et al. Splicing factor 1 modulates dietary restriction and TORC1 pathway longevity in C. elegans. Nature 2017, 541(7635): 102–106.

54. Kuroyanagi H, Kobayashi T, Mitani S, Hagiwara M. Transgenic alternative-splicing reporters reveal tissue-specific expression profiles and regulation mechanisms in vivo. Nat Methods 2006, 3(11): 909–915.

55. Kuroyanagi H, Watanabe Y, Suzuki Y, Hagiwara M. Position-dependent and neuron-specific splicing regulation by the CELF family RNA-binding protein UNC-75 in Caenorhabditis elegans. Nucleic Acids Res 2013, 41(7): 4015–4025.

56. Ventura N, Rea SL. Caenorhabditis elegans mitochondrial mutants as an investigative tool to study human neurodegenerative diseases associated with mitochondrial dysfunction. Biotechnology Journal 2007, 2(5): 584–595.

57. Mori C, Takanami T, Higashitani A. Maintenance of mitochondrial DNA by the Caenorhabditis elegans ATR checkpoint protein ATL-1. Genetics 2008, 180(1): 681–686.

58. Henderson ST, Bonafe M, Johnson TE. daf-16 protects the nematode Caenorhabditis elegans during food deprivation. J Gerontol A Biol Sci Med Sci 2006, 61(5): 444–460.

59. Bloss TA, Witze ES, Rothman JH. Suppression of CED-3-independent apoptosis by mitochondrial [beta]NAC in Caenorhabditis elegans. Nature 2003, 424(6952): 1066–1071.

60. Arsenovic PT, Maldonado AT, Colleluori VD, Bloss TA. Depletion of the C. elegans NAC engages the unfolded protein response, resulting in increased chaperone expression and apoptosis. PLoS ONE 2012, 7(9): e44038.

61. Kim SK, Lund J, Kiraly M, Duke K, Jiang M, Stuart JM, et al. A gene expression map for Caenorhabditis elegans. Science 2001, 293(5537): 2087–2092.

62. Houtkooper RH, Mouchiroud L, Ryu D, Moullan N, Katsyuba E, Knott G, et al. Mitonuclear protein imbalance as a conserved longevity mechanism. Nature 2013, 497(7450): 451–457.

63. West AP, Khoury-Hanold W, Staron M, Tal MC, Pineda CM, Lang SM, et al. Mitochondrial DNA stress primes the antiviral innate immune response. Nature 2015, 520(7548): 553–557.

64. Lewis SC, Joers P, Willcox S, Griffith JD, Jacobs HT, Hyman BC. A rolling circle replication mechanism produces multimeric lariats of mitochondrial DNA in Caenorhabditis elegans. PLoS Genet 2015, 11(2): e1004985.

65. Starokadomskyy P, Gemelli T, Rios JJ, Xing C, Wang RC, Li H, et al. DNA polymerase-alpha regulates the activation of type I interferons through cytosolic RNA:DNA synthesis. Nat Immunol 2016, 17(5): 495–504.

66. Milenkovic D, Blaza JN, Larsson NG, Hirst J. The Enigma of the Respiratory Chain Supercomplex. Cell Metab 2017, 25(4): 765–776.

67. George R, Beddoe T, Landl K, Lithgow T. The yeast nascent polypeptide-associated complex initiates protein targeting to mitochondria in vivo. Proc Natl Acad Sci U S A 1998, 95(5): 2296–2301.

68. George R, Walsh P, Beddoe T, Lithgow T. The nascent polypeptide-associated complex (NAC) promotes interaction of ribosomes with the mitochondrial surface in vivo. FEBS Lett 2002, 516(1-3): 213–216.

69. Yogev O, Karniely S, Pines O. Translation-coupled translocation of yeast fumarase into mitochondria in vivo. J Biol Chem 2007, 282(40): 29222–29229.

70. Yang K-S, Kim H-S, Jin U-H, Lee S, Park J-A, Lim Y, et al. Silencing of NbBTF3 results in developmental defects and disturbed gene expression in chloroplasts and mitochondria of higher plants. Planta 2007, 225(6): 1459–1469.

71. del Alamo M, Hogan DJ, Pechmann S, Albanese V, Brown PO, Frydman J. Defining the specificity of cotranslationally acting chaperones by systematic analysis of mRNAs associated with ribosome-nascent chain complexes. PLoS Biology 2011, 9(7): e1001100.

72. Amanchy R, Periaswamy B, Mathivanan S, Reddy R, Tattikota SG, Pandey A. A curated compendium of phosphorylation motifs. Nat Biotechnol 2007, 25(3): 285–286.

73. Johnson SC, Yanos ME, Kayser EB, Quintana A, Sangesland M, Castanza A, et al. mTOR inhibition alleviates mitochondrial disease in a mouse model of Leigh syndrome. Science 2013, 342(6165): 1524–1528.

74. Zid BM, Rogers AN, Katewa SD, Vargas MA, Kolipinski MC, Lu TA, et al. 4E-BP extends lifespan upon dietary restriction by enhancing mitochondrial activity in Drosophila. Cell 2009, 139(1): 149–160.

75. Bonawitz ND, Chatenay-Lapointe M, Pan Y, Shadel GS. Reduced TOR Signaling Extends Chronological Life Span via Increased Respiration and Upregulation of Mitochondrial Gene Expression. Cell Metab 2007, 5(4): 265–277.

76. Garelick MG, Mackay VL, Yanagida A, Academia EC, Schreiber KH, Ladiges WC, et al. Chronic rapamycin treatment or lack of S6K1 does not reduce ribosome activity in vivo. Cell Cycle 2013, 12(15): 2493–2504.

77. Wood WB (ed). The Nematode Caenorhabditis elegans. Cold Spring Harbor Laboratory: New York, 1988.

78. Timmons L, Fire A. Specific interference by ingested dsRNA. Nature 1998, 395(6705): 854.

79. Hoogewijs D, Houthoofd K, Matthijssens F, Vandesompele J, Vanfleteren JR. Selection and validation of a set of reliable reference genes for quantitative sod gene expression analysis in C. elegans. BMC Mol Biol 2008, 9: 9.

80. Sugimoto T, Mori C, Takanami T, Sasagawa Y, Saito R, Ichiishi E, et al. Caenorhabditis elegans par2.1/mtssb-1 is essential for mitochondrial DNA replication and its defect causes comprehensive transcriptional alterations including a hypoxia response. Experimental Cell Research 2008, 314(1): 103–114.

81. Rooney JP, Ryde IT, Sanders LH, Howlett EH, Colton MD, Germ KE, et al. PCR based determination of mitochondrial DNA copy number in multiple species. Methods Mol Biol 2015, 1241: 23–38.

82. Kuroyanagi H, Ohno G, Sakane H, Maruoka H, Hagiwara M. Visualization and genetic analysis of alternative splicing regulation in vivo using fluorescence reporters in transgenic Caenorhabditis elegans. Nat Protoc 2010, 5(9): 1495–1517.

83. Luz AL, Rooney JP, Kubik LL, Gonzalez CP, Song DH, Meyer JN. Mitochondrial Morphology and Fundamental Parameters of the Mitochondrial Respiratory Chain Are Altered in Caenorhabditis elegans Strains Deficient in Mitochondrial Dynamics and Homeostasis Processes. PLoS One 2015, 10(6): e0130940.

84. Steffen KK, MacKay VL, Kerr EO, Tsuchiya M, Hu D, Fox LA, et al. Yeast life span extension by depletion of 60s ribosomal subunits is mediated by Gcn4. Cell 2008, 133(2): 292–302.

85. Torgovnick A, Schiavi A, Shaik A, Kassahun H, Maglioni S, Rea SL, et al. BRCA1 and BARD1 mediate apoptotic resistance but not longevity upon mitochondrial stress in Caenorhabditis elegans. EMBO Rep 2018, 19(12).

