## Supplementary figures and images for "Inhibition of ATR Reverses a Mitochondrial Respiratory Insufficiency"

### Supplemental Figure S1

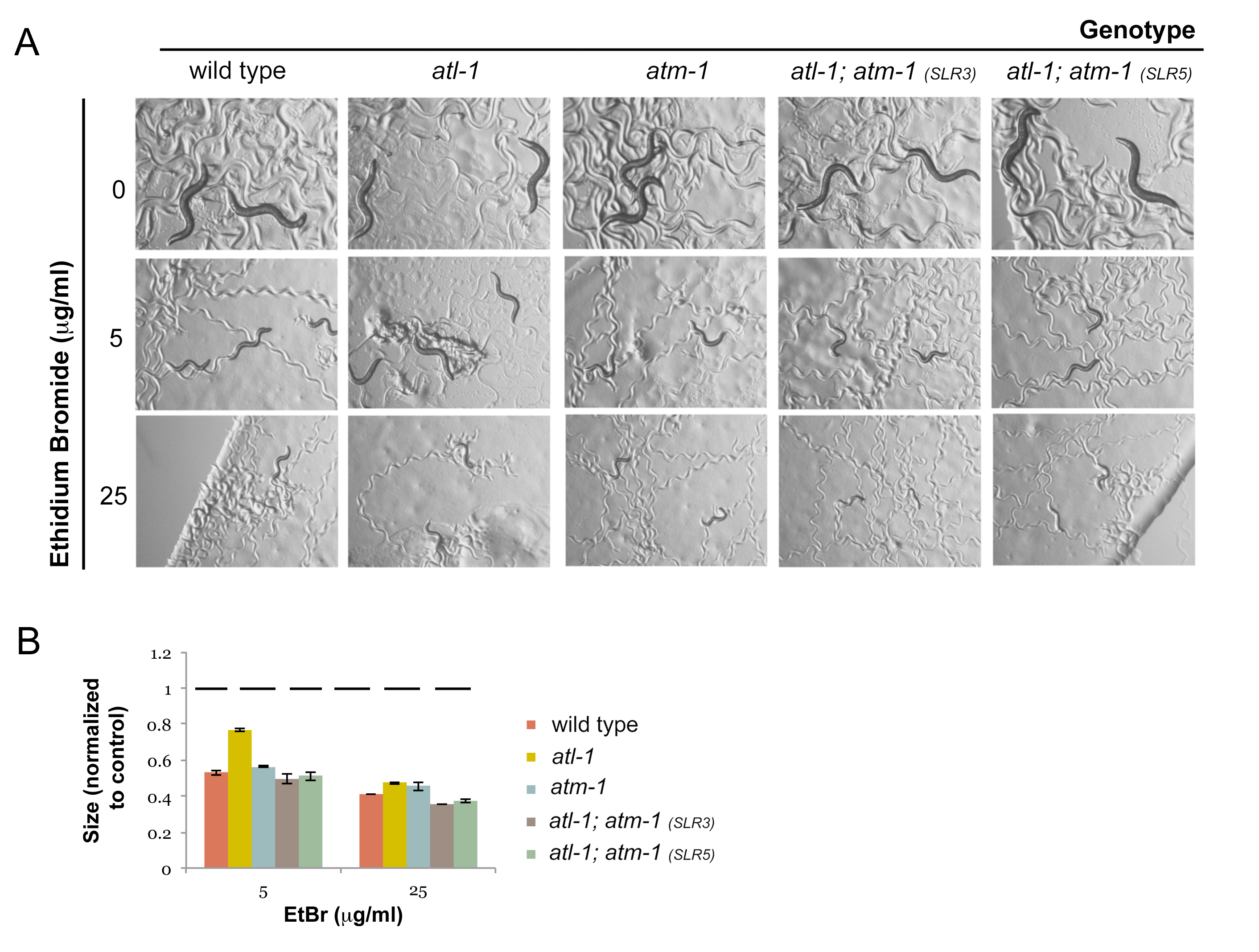

### Supplemental Figure S2

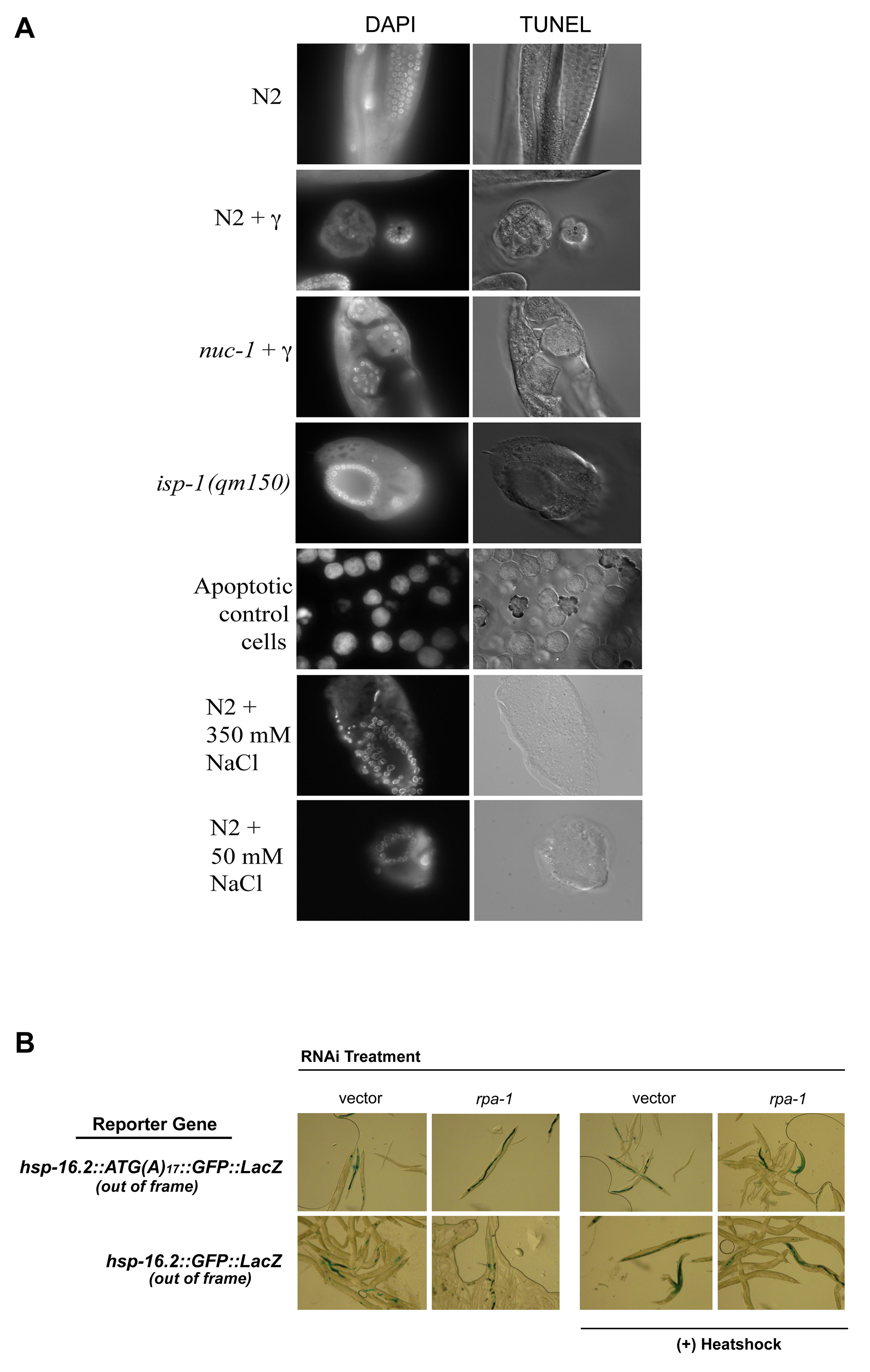

### Supplemental Figure S3

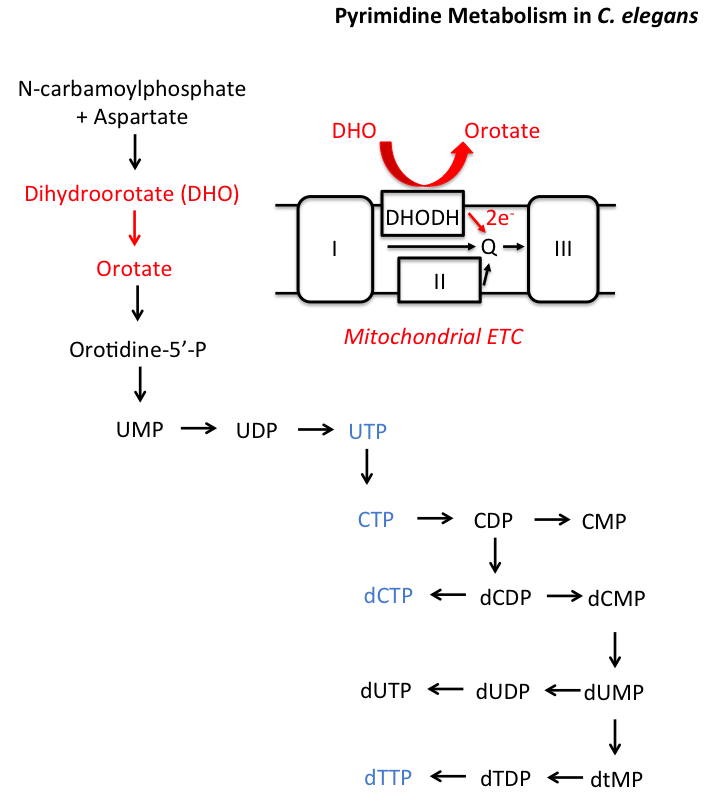

### Supplemental Figure S4

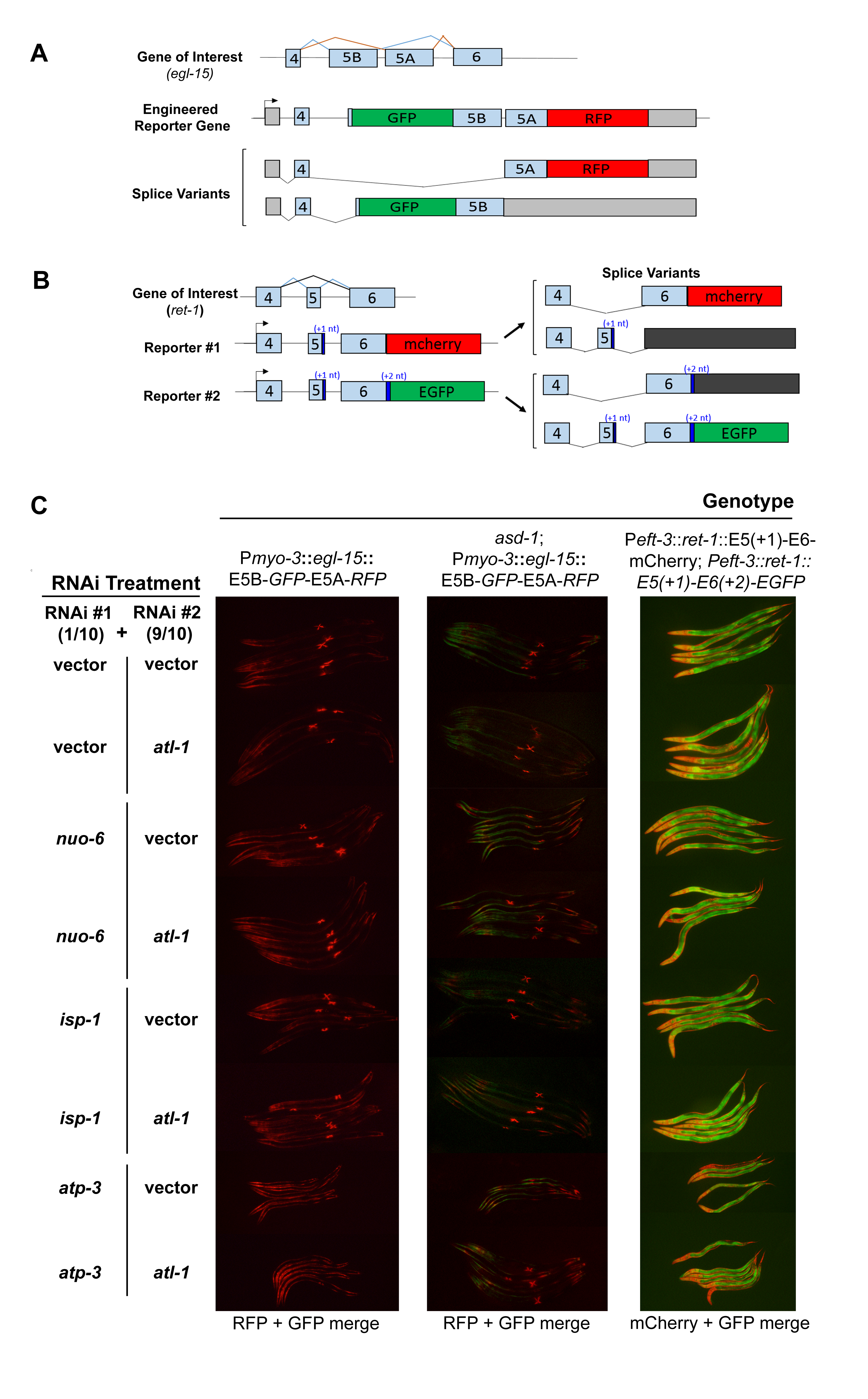

### Supplemental Figure S5

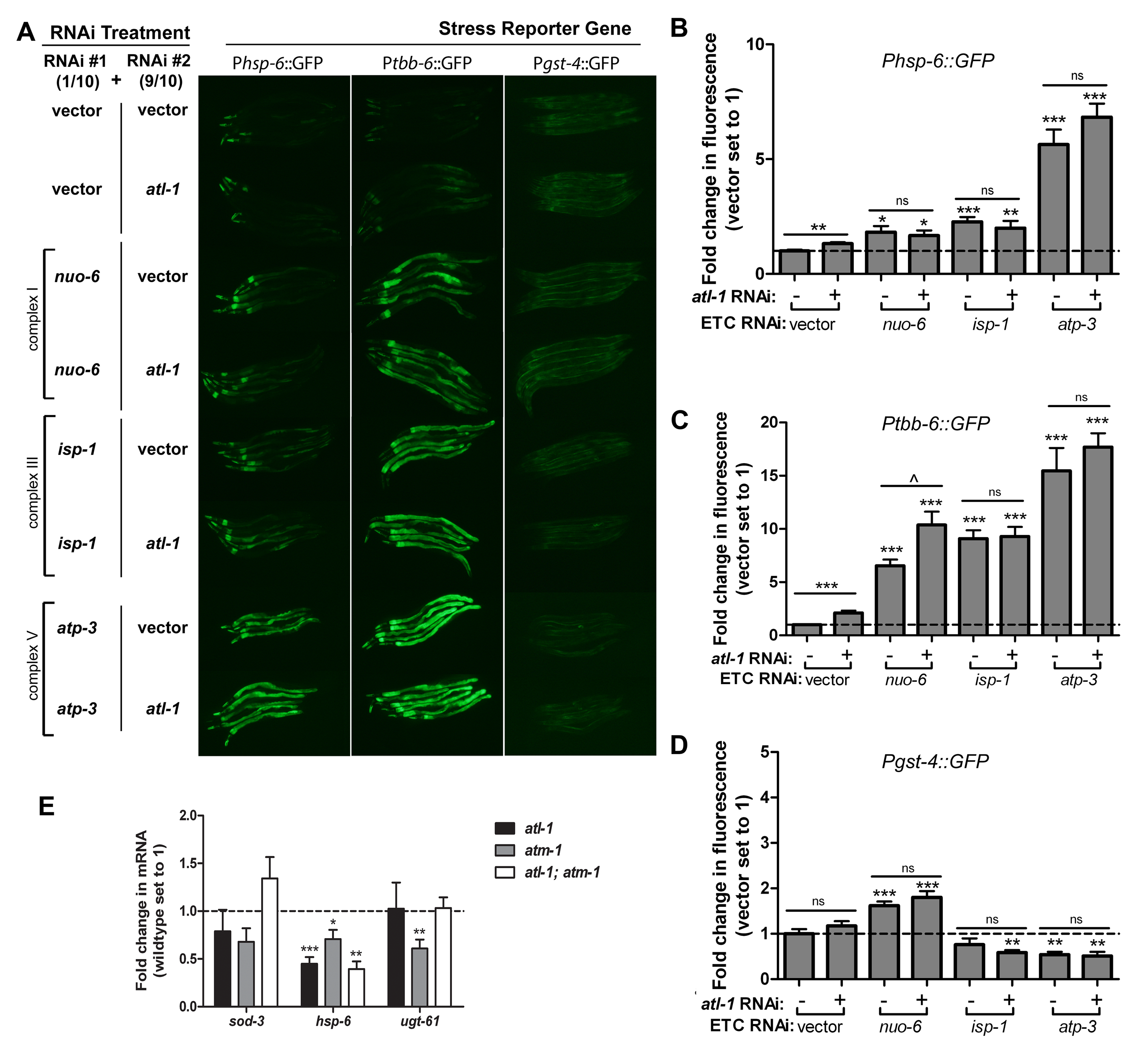

### Supplemental File S2

#### Slide 1
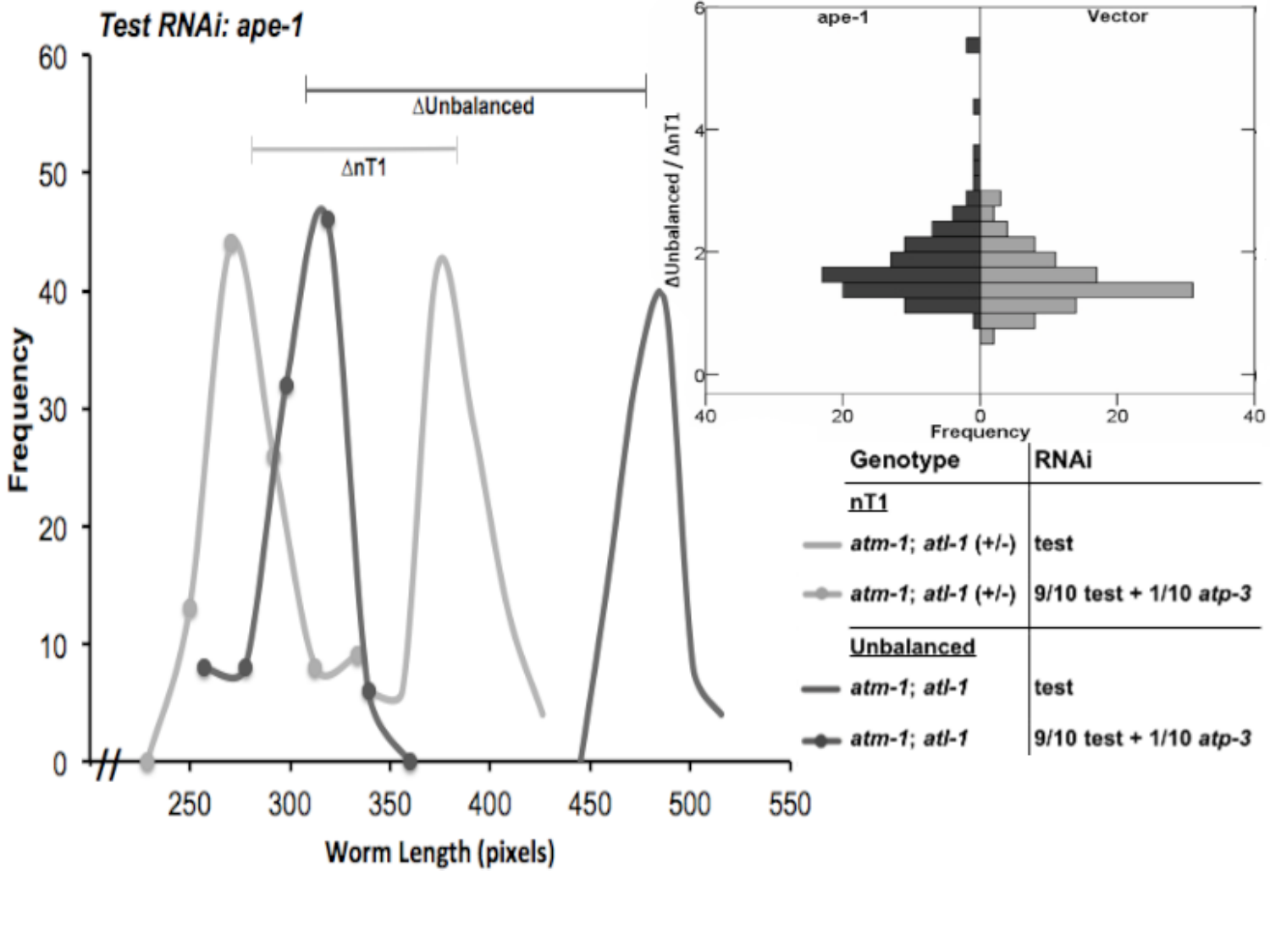

#### Slide 2
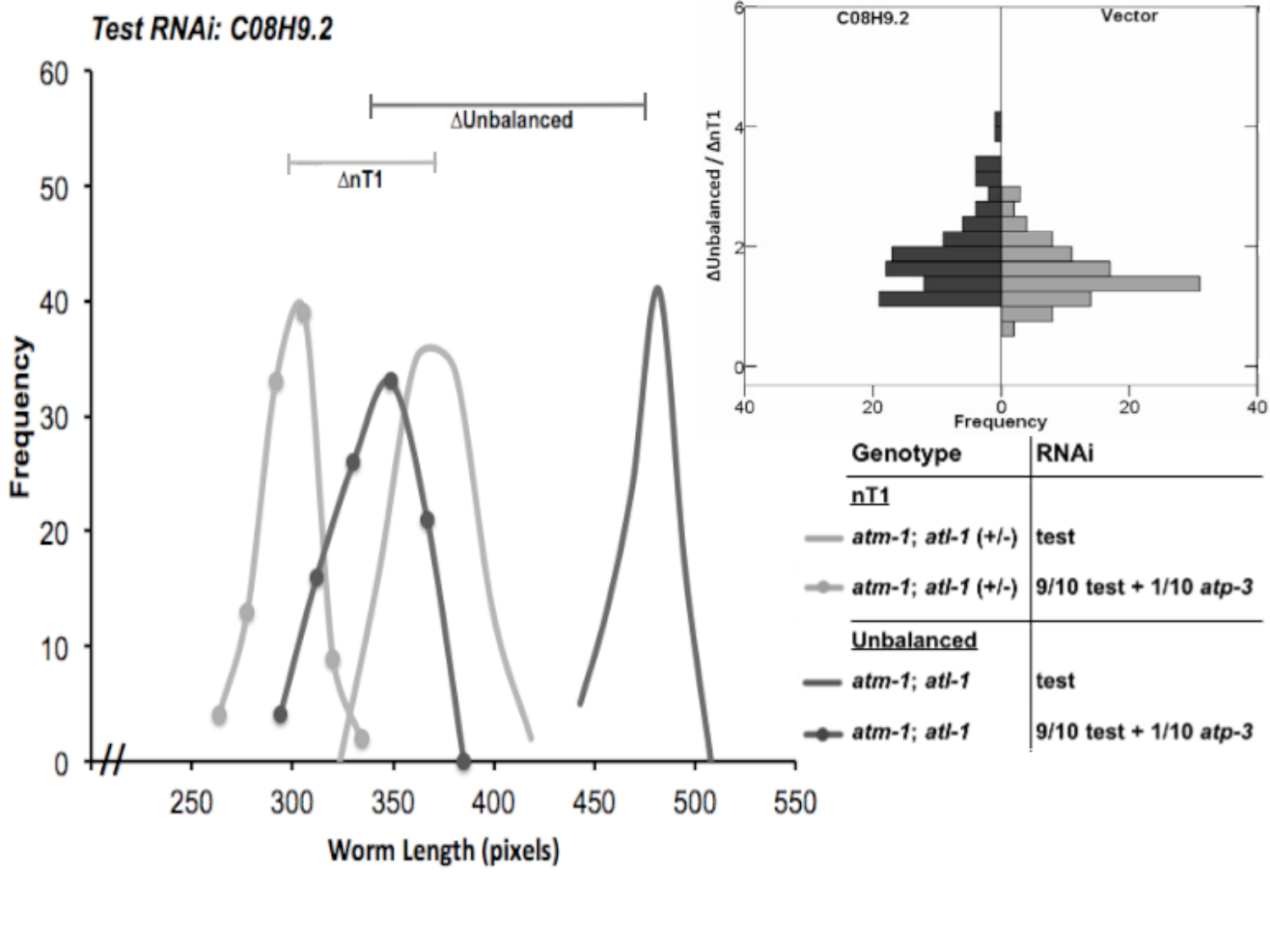

#### Slide 3
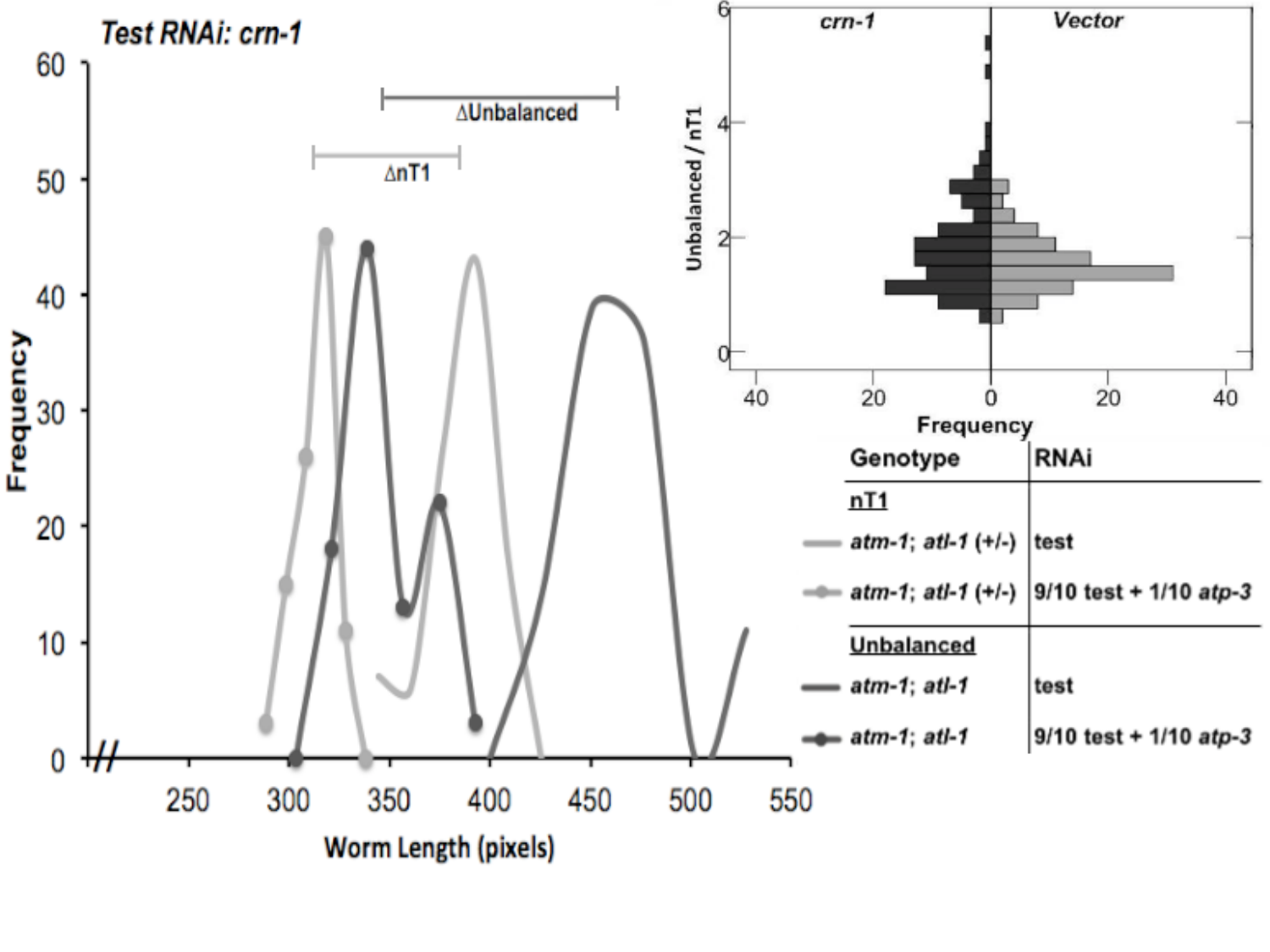

#### Slide 4
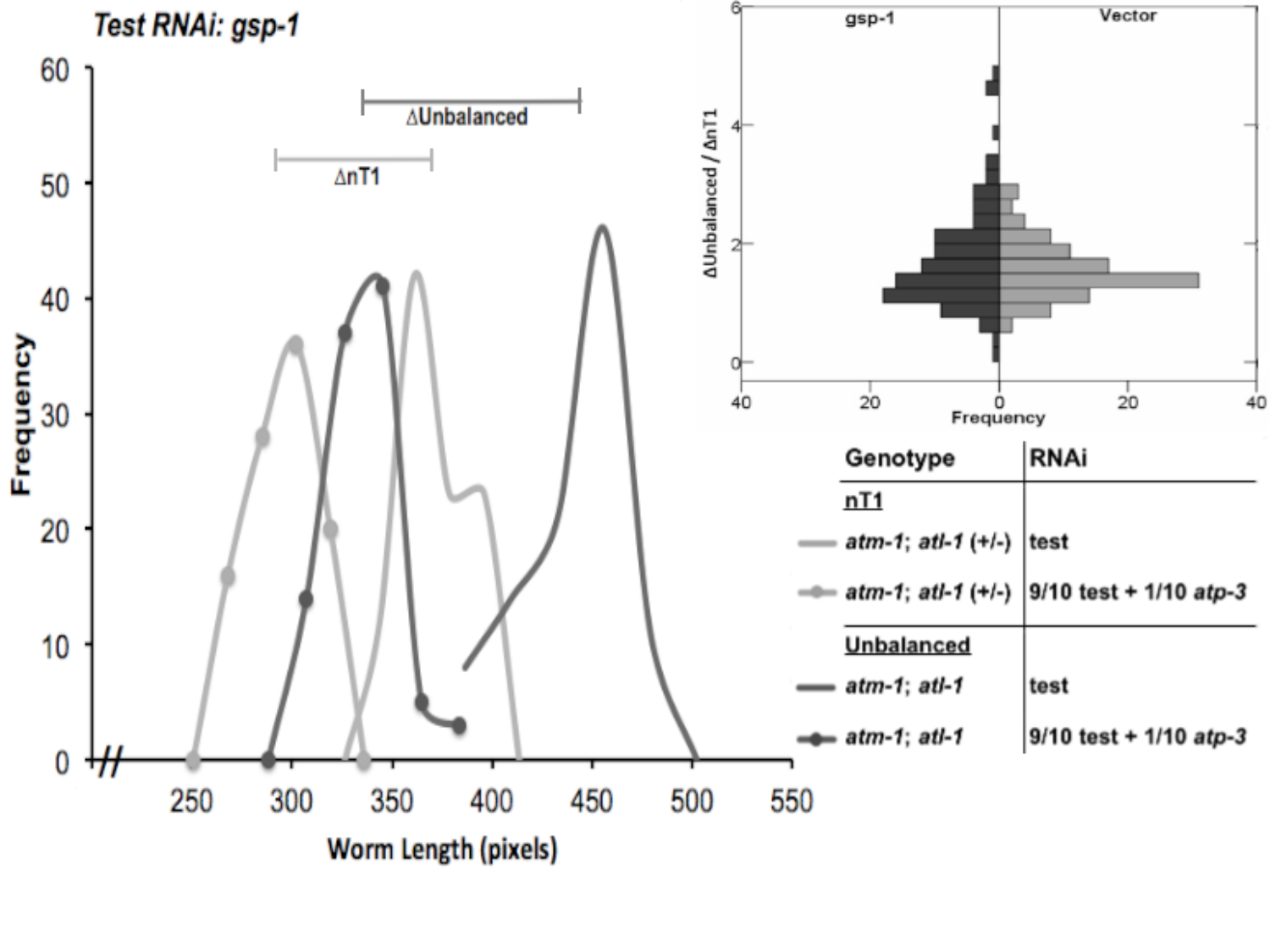

#### Slide 5
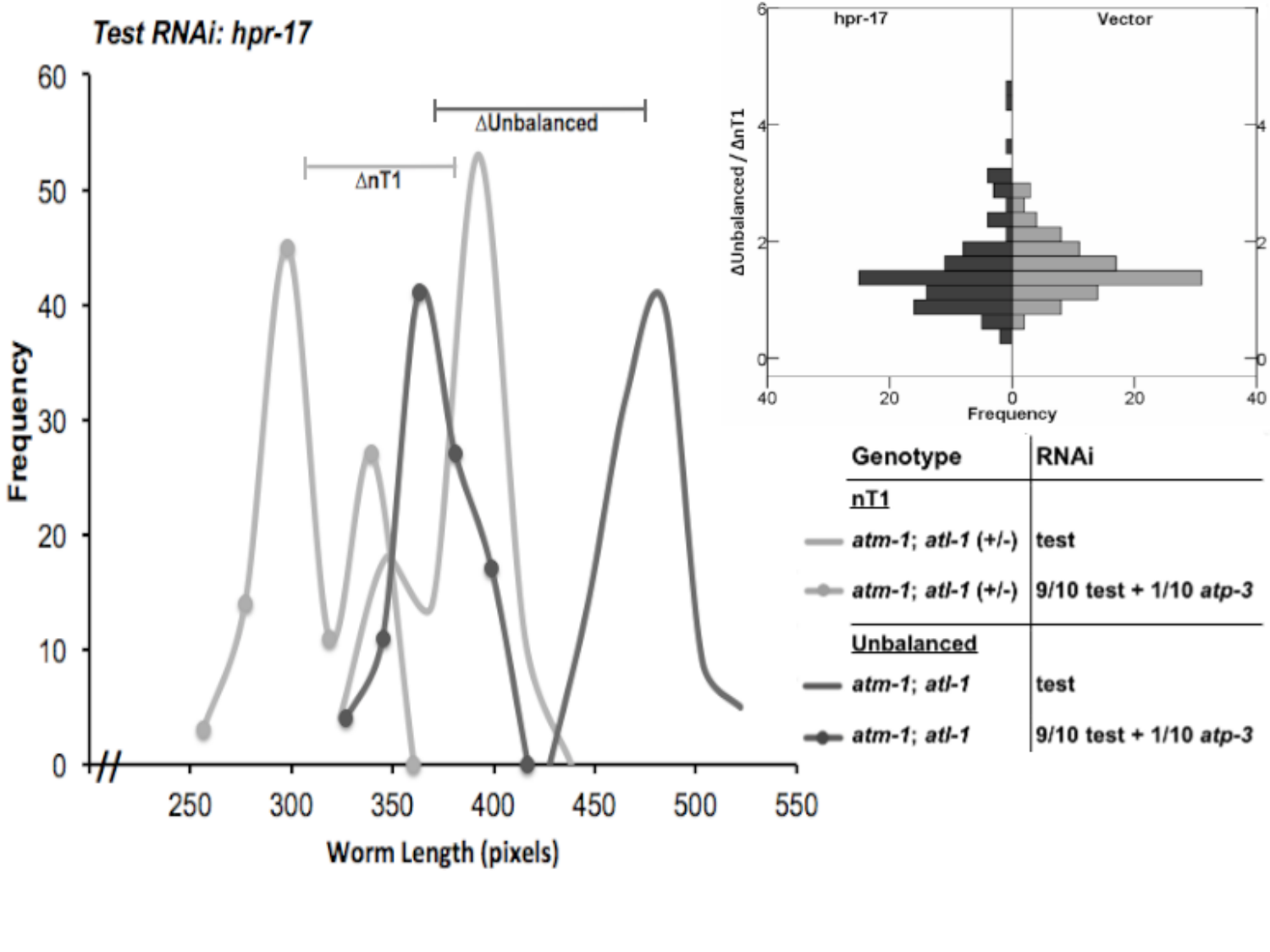

#### Slide 6
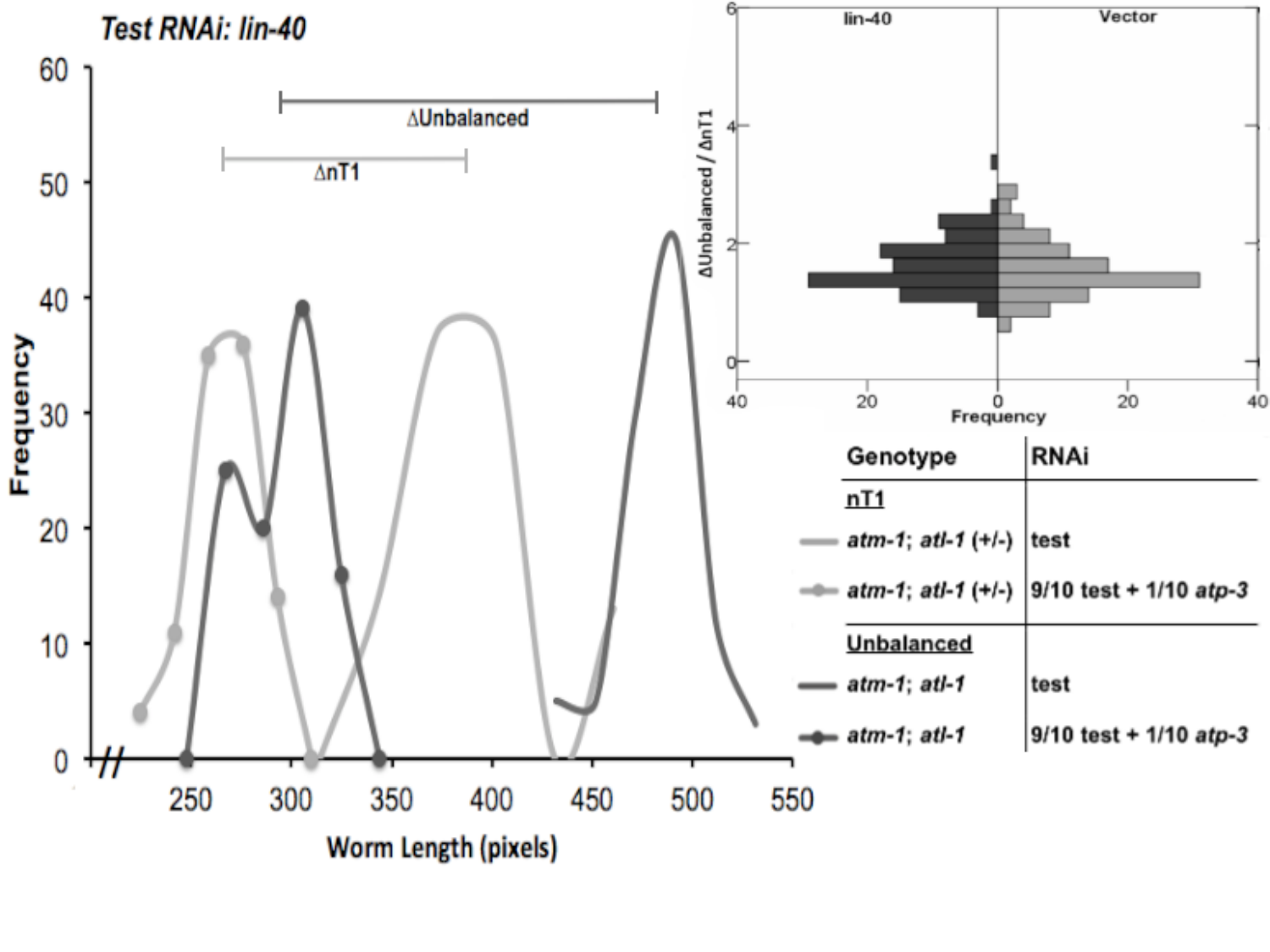

#### Slide 7
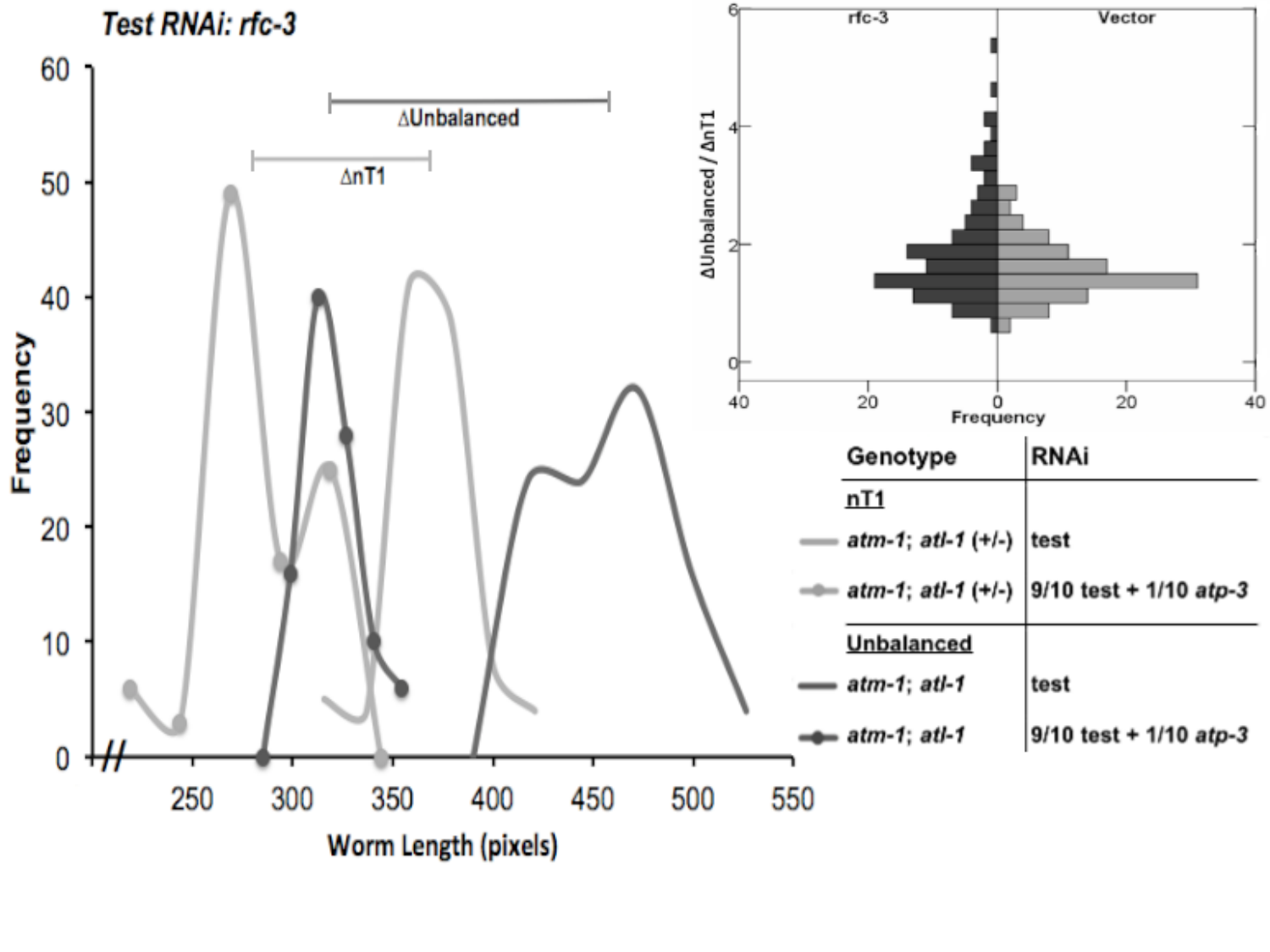

#### Slide 8
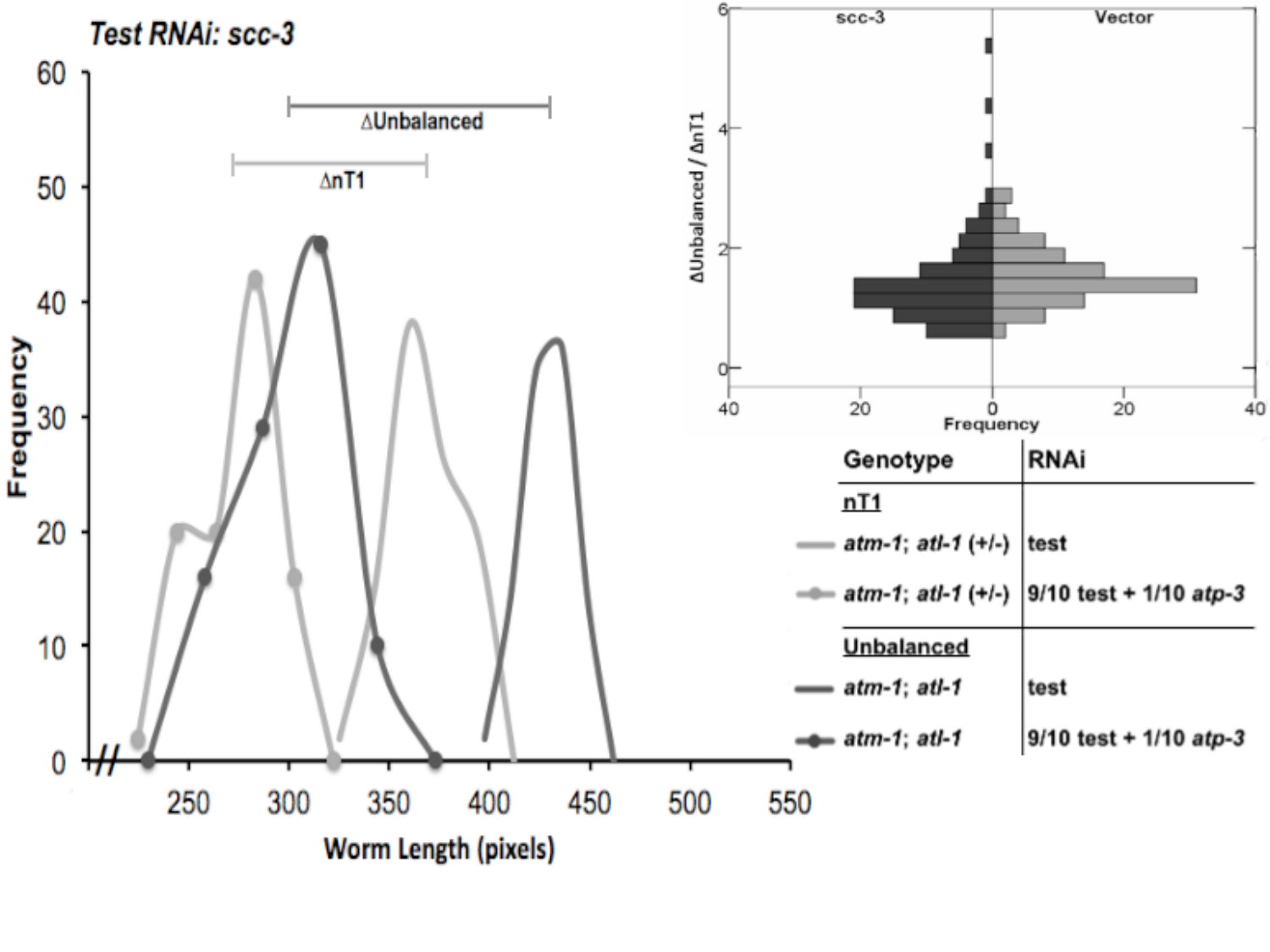

#### Slide 9
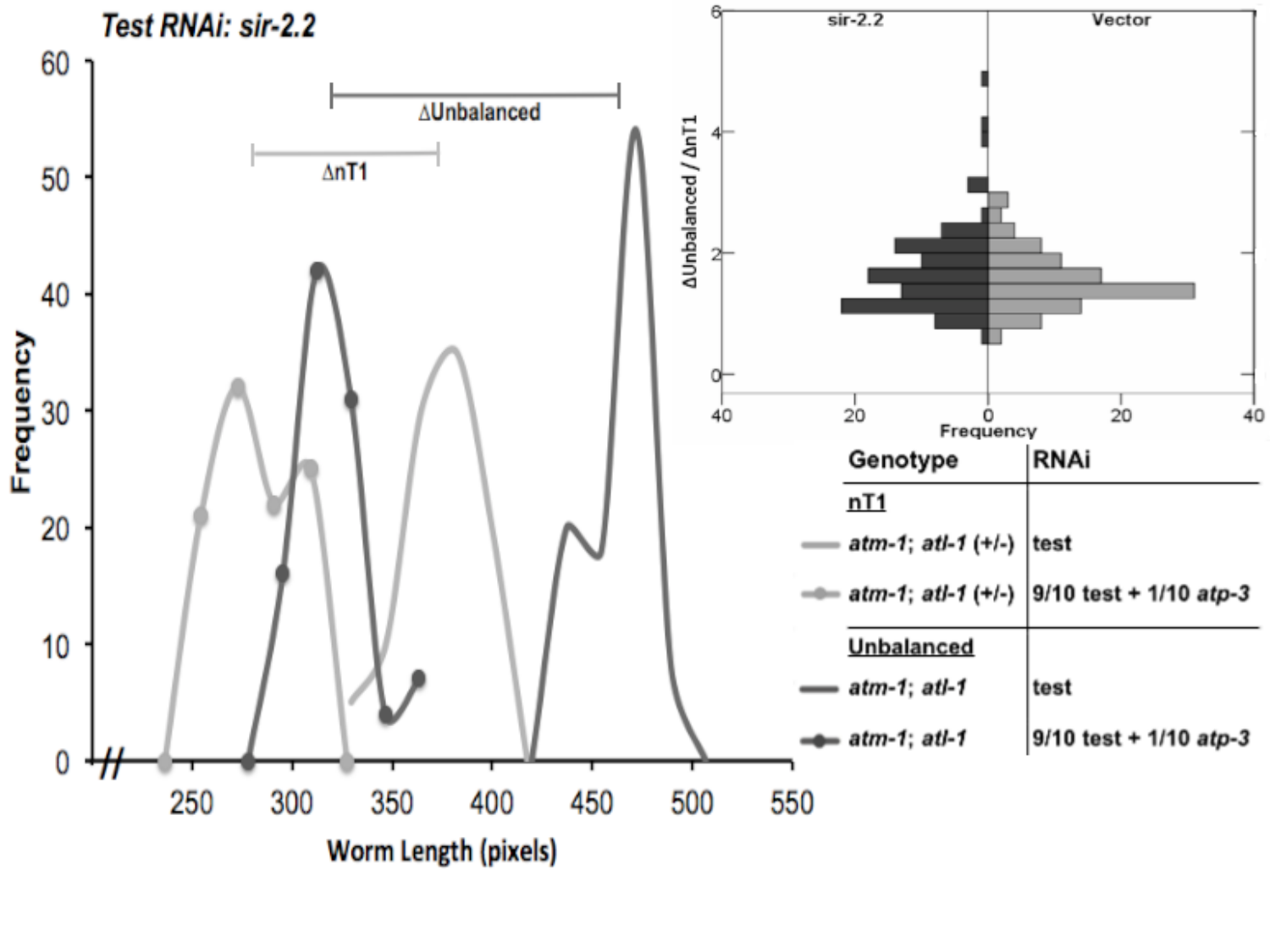

#### Slide 10
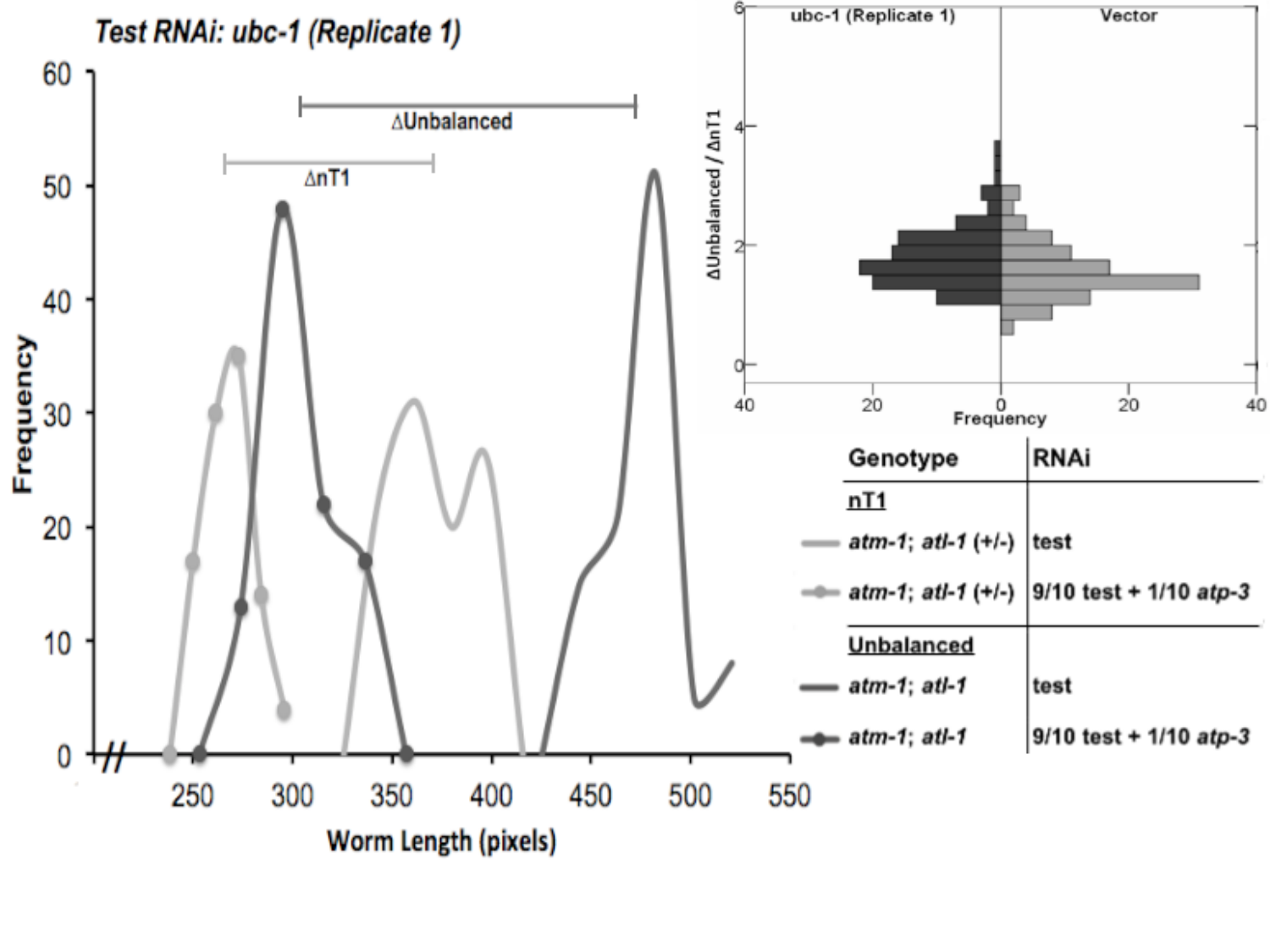

#### Slide 11
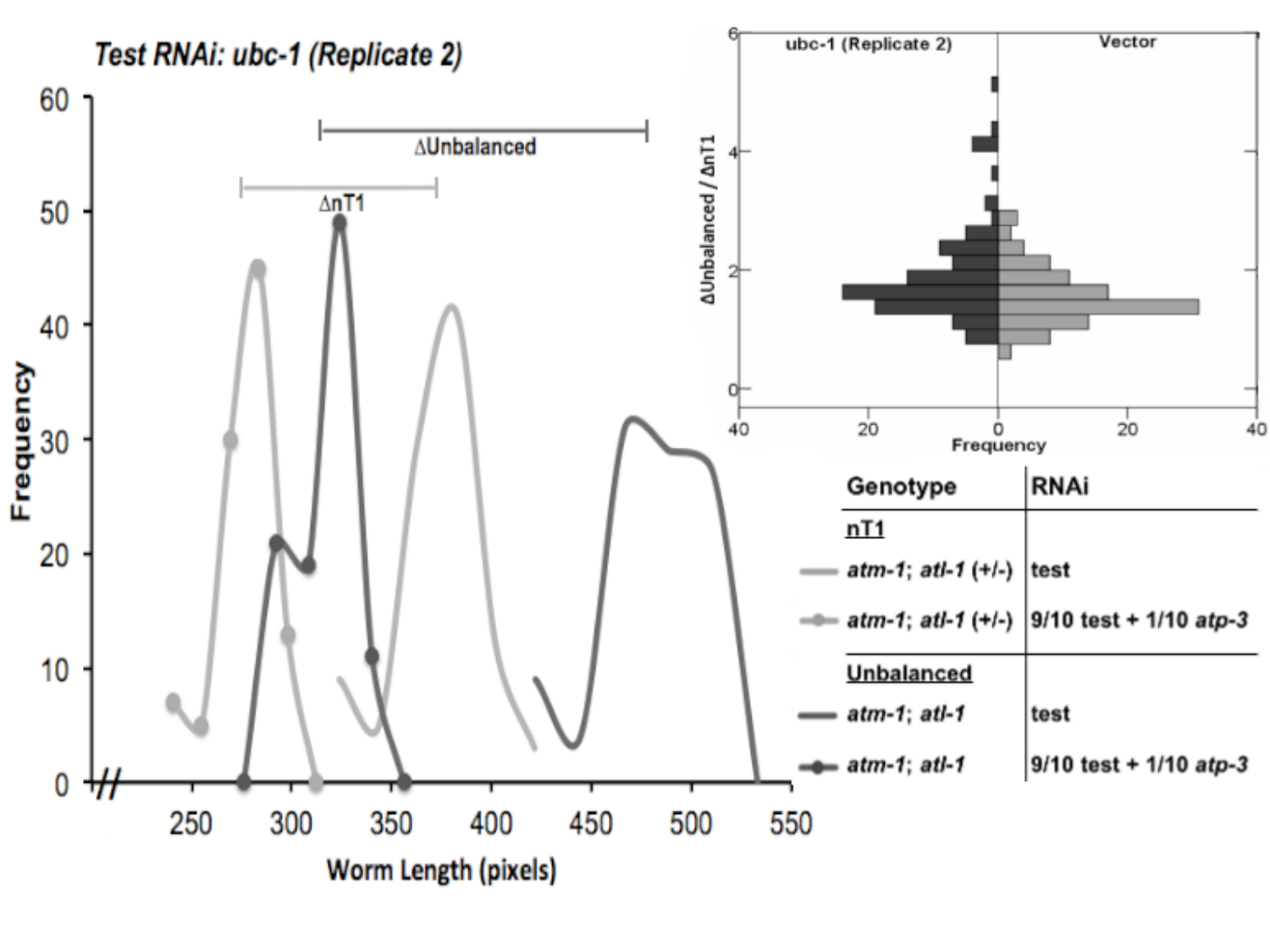

#### Slide 12
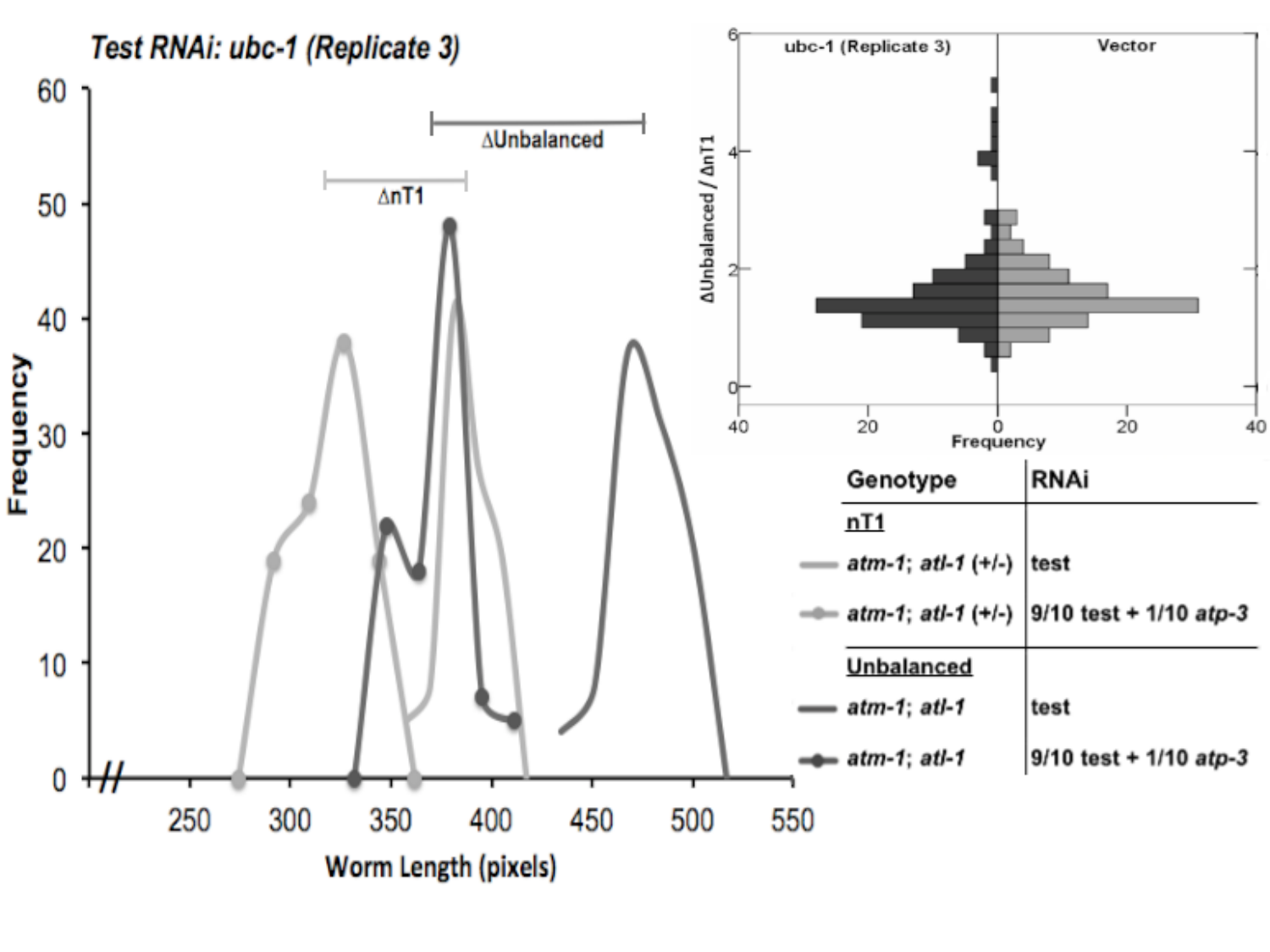

#### Slide 13
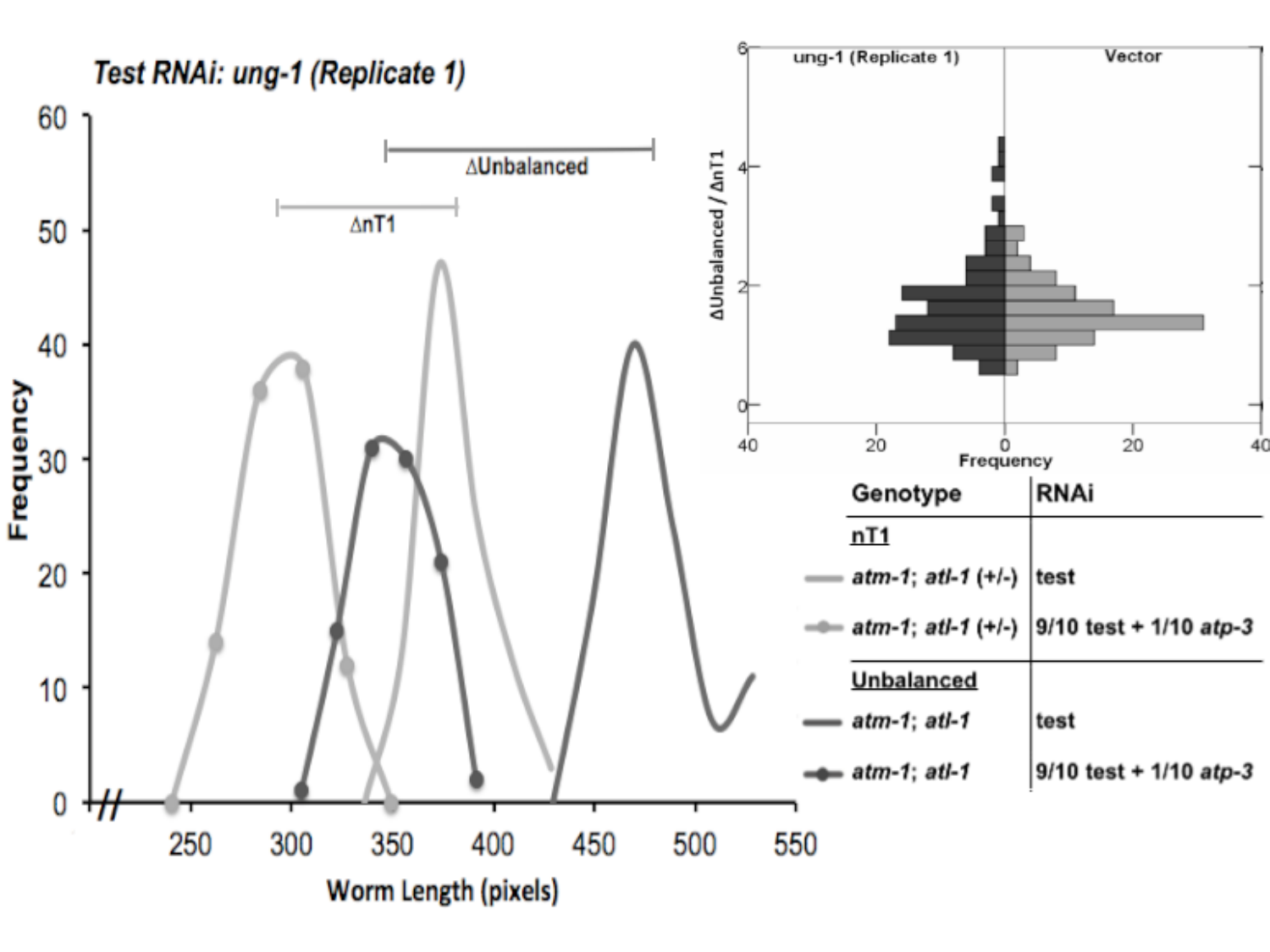

#### Slide 14
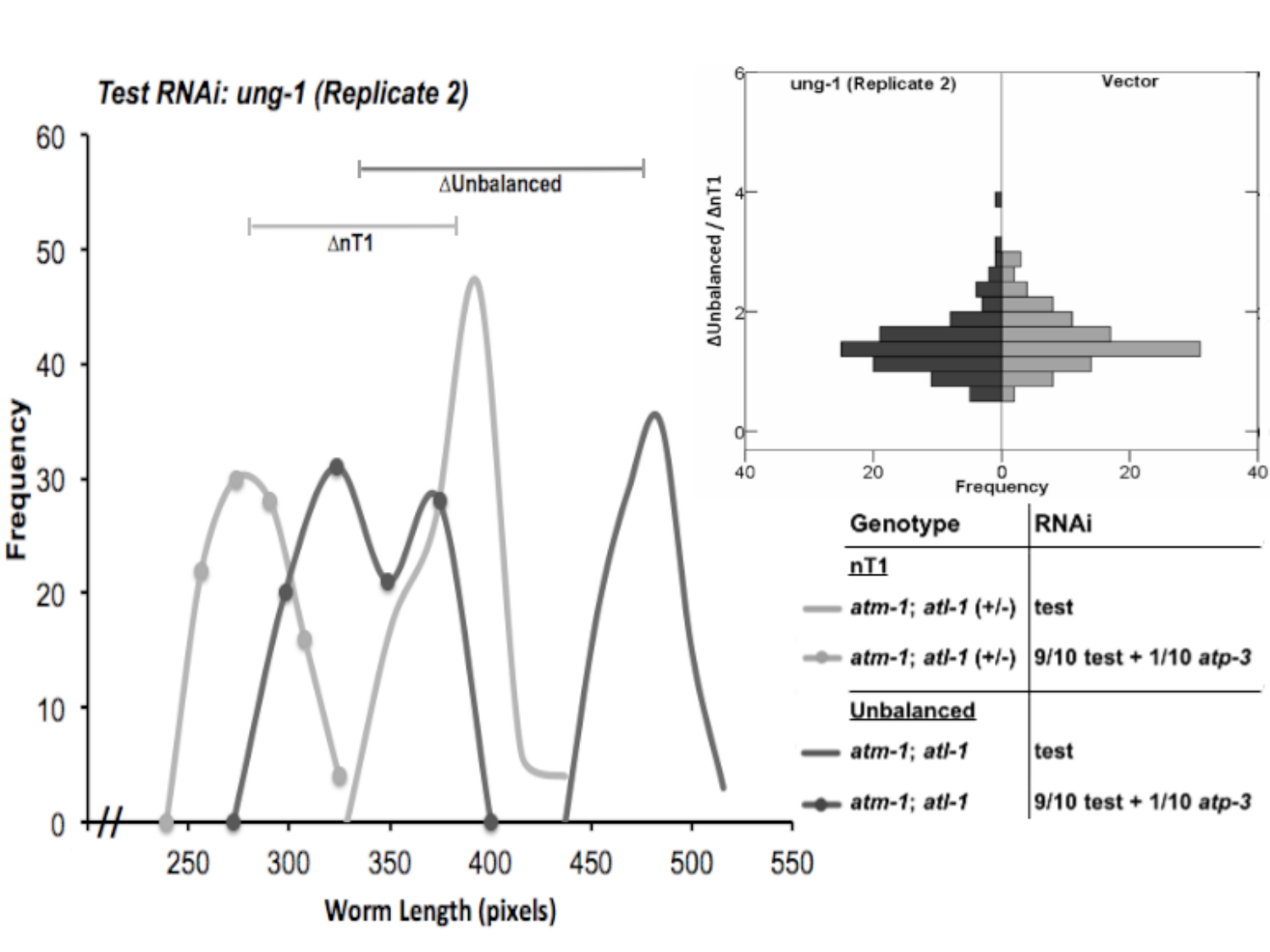

#### Slide 15
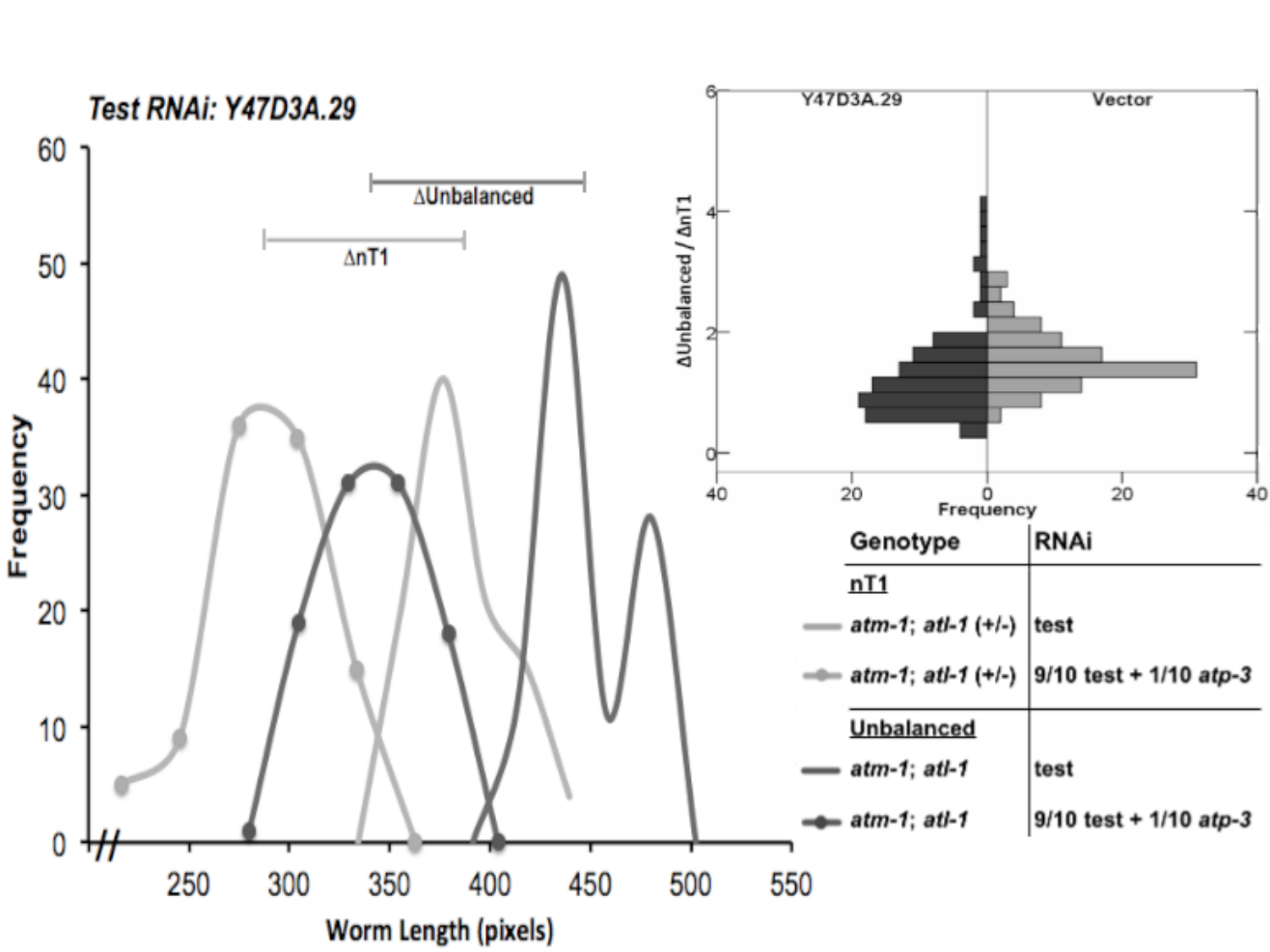
