## Supplemental File S1 for "Inhibition of ATR Reverses a Mitochondrial Respiratory Insufficiency"

### SUPPLEMENTAL METHODS

**Nematode Strains and Maintenance.** Full genotypic information for all strains used in this study is provided in **Table S4**. Lines were maintained at 20˚C on standard NGM agar plates seeded with *E. coli* (OP50) [1]. To investigate a potential role for DNA damage-sensing PIKKs in modulating the mitochondrial threshold effect in *C. elegans*, we tested if worms containing loss-of-function mutations in *atm-1* or *atl-1* responded differently from wild type worms to ETC disruption. *atm-1(gk186)* mutants are homozygous viable and contain a 548 bp deletion that results in two aberrant *atm-1* transcripts, both of which encode non-functional proteins [2]. *atl-1(tm853)* mutants are maternal-effect lethal, meaning only homozygous *atl-1(tm853)* progeny derived from *atl-1(tm853)/+* heterozygotes are viable. The *atl-1(tm853)* mutation contains a 720 bp deletion in the *atl-1* locus that results in no detectable ATL-1 protein by western analysis [3]. Both mutations were moved into the nT1(IV:V) reciprocal chromosomal translocation background. For each new experiment we therefore selected fresh, unbalanced F1 progeny. We also generated two independent *atm-1(gk186); atl-1(tm853)* double mutant nT1 lines. As well, we constructed and maintained a control nT1 line that contained no additional mutations and re-generated the wild type genotype upon loss of the nT1 chromosomal pair. This approach of equally using nT1 to carry all test genotypes, whether required or not, circumvented concerns of a possible untoward maternal effect caused by the nT1 translocation itself.

**Bacterial Feeding RNAi.** All bacterial feeding RNAi constructs were designed in the pL4440 vector backbone and maintained in HT115 bacteria [4]. RNAi clones targeting *atp-3, isp-1, nuo-6* and *skn-1* have been described previously, along with the procedure for feeding RNAi dilution [5, 6]. RNAi targeting *atl-1* was constructed using the following PCR primers and wild type (N2) genomic DNA as template: Forward primer: 5'-CTGTCTAAGCTTATTGAACGGCTGTCTGAATGTGC-3', Reverse primer: 5'-

GATCTCTCGAGCGATCGGGCAAATGACAAGATTCC-3'. Feeding RNAi constructs for DDR screening were obtained from the Ahringer RNAi library [7]. For assays comparing growth rate and life span across strains (**Figs. 1** and **2**, main text), eggs of the same chronological age from all strains were simultaneously seeded onto RNAi lawns that were prepared in unison. RNAi efficacy in *ncl-1* mutants was tested as follows: Worms were fed bacteria containing increasing amounts of *skn-1* RNAi from the arrested L1 stage onward. At the L4/YA boundary, exactly 20 animals were transferred to fresh *skn-1* RNAi plates and then allowed to lay eggs for another 24 hours. The number of dead eggs and hatchlings on each plate was then counted (~1000 progeny per plate).

**Lifespan Analyses**. Lifespan studies were performed as described previously [8], without the use of FUdR. All plates were maintained at 20ºC. The Log-rank test was used to analyze the effect of gene knock-out or RNAi knockdown. The first day of adulthood was designated as day one, unless otherwise noted. Animals that bagged, desiccated, or suffered gonad extrusion were scored as censored individuals. Such animals were included in life-span analyses up to the point of censorship and were weighted by half in mortality calculations.

**Microscopy.** Images of fluorescent worms were captured using an Olympus SZX16 fluorescence microscope connected to an Olympus DP71 CCD camera. Worms were immobilized for imaging by transferring animals to plates at 4ºC. Total worm fluorescence was quantified using ImageJ [9]. All images were collected on the first day of adulthood unless otherwise specified. For strains containing the *Phsp-6::GFP* or *Ptbb-6::GFP* reporter gene, background fluorescence in the head was excluded from signal integration. Six to 10 worms were quantified for each tested condition. For worm length measurements, the Segmented Line function of ImageJ was used in conjunction with worm DIC images [9]. Up to 10 worms were quantified for each tested condition. For confocal imaging, worms containing the HXK2::GFP mitochondrial reporter were mounted on 5% agarose pads in 20 µL of 1 mM levamisole and fluorescent images collected using a Zeiss LSM 780 confocal microscope. In **Fig. 7C**, to determine whether differences existed in the morphology of mitochondria between the four tested conditions, four randomly selected images from each condition were mixed and then grouped by a scorer who was blind to image identity. In a set of 16 images, 10 were co-grouped correctly. The probability that this grouping occurred by chance was modeled using an Excel VBA macro that randomly binned the images into four groups of four, 10,000 times (**File S2**). The final *p-value* was <0.0024 and deemed significant.

**mRNA and mtDNA Quantitation.** For quantitation of mRNA, 1000 first-day adult worms were washed in S-Basal [1], flash frozen in liquid nitrogen, then RNA extracted with an RNeasy kit (Qiagen). cDNA was generated using a High Capacity cDNA Reverse Transcription kit (Applied Biosystems). Quantitative PCR was performed using a 7500 Real Time PCR (Applied Biosystems) in conjunction with $RT^2$ SYBR Green/ROX reagent (Qiagen). Fold change in each mRNA of interest was determined using the ΔΔCt method normalized to the geometric mean of *cdc-42*, *pmp-3*, and *Y45F10D.4* [10]. For quantitation of mitochondrial DNA (mtDNA), PCR primer pairs targeting mtDNA-encoded *ctb-1* and intron 4 of genomic DNA-encoded *ama-1* were synthesized and used as described [11]. The ratio of mitochondrial DNA to nuclear DNA was then calculated using Real Time PCR, using *ama-1* as the normalizing quantity [12]. A complete list of qPCR primers is provided in **Table S5**.

**Mutation Screening Assays:**

<u>Unc Reversion Assays</u>: We used frequency of Unc reversion in *unc-93(e1500)* and *unc-58(e665)* mutants exposed to mitochondrial ETC disruption as one measure of nuclear DNA mutation rate. For each genotype, 50 L1 larvae were seeded individually onto 10 cm bacterial feeding RNAi lawns targeting *atp-3* (1/20$^{th}$ strength); *rpa-1* (/10$^{th}$ strength) or vector-control.

Once each worm population had starved, animals were collected in S-basal then transferred to a 2% peptone-enriched NGM-agar plate (10cm) seeded with wild type *E. coli* (strain RW2) and allowed to starve again. This step was repeated a second time. By this stage, plates containing revertants were easily identifiable and each such plate was scored as a single mutagenic event. Significance testing was undertaken using a Fisher's exact test, comparing number of lethal mutations detected in the *atp-3* population against vector control. Significance was set at $p<0.05$. *rpa-1* treated worms served as a positive control.

*Lethal Mutation Assays:* To score nuclear DNA mutation rate, 50 worms containing the eT1(III;V) (strain BC2200, **Table S4**) or the nT1(IV;V) (strains TJ5013 and SLR004) reciprocal chromosomal translocations were cultured from the L1 larval stage on bacterial feeding RNAi lawns targeting *atp-3* (across a range of dilutions) or vector control. Four hundred F1 progeny containing eT1 or nT1 were subsequently collected and then individually singled onto 6cm NGM/OP50 plates. Absence of Dpy Unc (BC2200) or wild type (TJ5013, SLR004) F2 progeny indicated the presence of a balanced, lethal mutation event in the F1 parent. Significance testing was undertaken using a Fisher's exact test, comparing number of lethal mutations detected in each *atp-3* population against vector control. Significance was set at $p<0.05$.

*Lac-Z Frameshift Assay:* Strain NL3400 contains the P*hsp-16.2::ATG(A)*$_{17}$*::GFP::LacZ* out-of-frame reporter gene. Mutations that disrupt the poly-(A) leader sequence and bring the reporter gene into frame provide a quantitative assessment of nuclear DNA mutation rate. Strain NL3401 contains an out-of frame reporter gene identical to that in NL3400 worms except for the absence of the 17 nucleotide adenine repeat. Differences in the mutation frequency between NL3400 and NL3401 indicate differences in the types of DNA damage mediating reporter frameshifting. To measure nuclear DNA mutation frequency, fifty NL3400 worms were cultivated for one generation on bacterial feeding RNAi targeting *atp-3* (1/20$^{th}$ strength); *rpa-1* (/10$^{th}$ strength) or vector-control. Following a one hour heat shock, β-galactosidase activity was measured [13]. This assay, we discovered, was confounded by the presence of functional β-galactosidase in the HT115 feeding RNAi bacteria and provided no reliable results, which is contrary to previous reports suggesting its successful use under such conditions [13]. Examples of our results are provided in **Supplemental Fig. S2B**. Disruption of DNA damage response genes often results in severe pathology and subsequent tissue invasion by bacteria may have been responsible for many of the apparent positive results in the original screens in which both reporter genes were developed and employed.

*TUNEL Assay:* The 'terminal deoxynucleotidyl transferase dUTP nick end labeling' (TUNEL) assay was used to measure levels of ssDNA and dsDNA breaks in *isp-1(qm150)* mutants relative to wild type (N2) worms. Briefly, animals were fixed in paraffin, sectioned, then stained for DNA breaks using the PromoKine Colorimetric DNA Fragmentation Detection Kit (IHC) (PK-CA577-K403) which utilizes Br-dUTP, a biotin-labeled anti-BrdU antibody, and a HRP-streptavidin conjugate to detect DNA lesions. For fixation and sectioning, washed worms (S-Basal) were immersed in 4% paraformaldehyde for 16 hr at 4°C then dehydrated using increasing concentrations of ethanol (70%, 80%, 95%, 100%) then xylene. Worms were subsequently embedded in paraffin and sectioned (3μm) using a Leica RM2125 microtome. Sections were serially rehydrated in ethanol (100%, 90%, 80%, 70%) and washed in phosphate buffered saline, then the tissue permeabilized with proteinase K. Sections were processed for TUNEL using the DNA Fragmentation Detection Kit and according to the manufacturer's instructions. Nuclei were counterstained with DAPI. Staining was imaged by bright field and fluorescence microscopy (Zeiss Axioskop 2). Controls included γ-irradiated N2 worms, mutant *nuc-1(e1392)* worms (1.4 Gy/min, 16hrs), as well as worms treated with elevated NaCl (350mM), which purportedly results in DNA strand breakage [14]. *nuc-1* encodes a DNase II homolog and is required for DNA degradation during apoptosis [15].

**Nucleotide Extraction and Mass Spectrometry.** Quantitation of deoxyribonucleotide and ribonucleotide species in whole-worm extracts of *isp-1(qm150)* and wild-type (N2) animals was undertaken using HPLC-MS. Briefly, 30,000 stage-synchronized worms were collected and washed in cold S-Basal [1]. Worms were resuspended in HPLC-grade water, then homogenized with a Balch homogenizer, as described previously [16]. Nucleotides were extracted using 6% TCA and water-saturated diethyl ether, following the procedure of Huang and colleagues [17]. Sample pH was neutralized with saturated $NaHCO_3$. LC-MS analyses were conducted using a Thermo Fisher/Dionex Ultimate 3000 HPLC connected in-line with a Thermo Fisher Q Exactive mass spectrometer. HPLC conditions were: Waters XTerra-MS C18 column (3.5 µm, 150 mm x 2.1 mm i.d.); mobile phase A – 5 mM Hexylamine and 0.5% diethylamine in water, pH 10; mobile phase B – 50% acetonitrile in water; flow rate – 400 µL/min; gradient – 1% B to 20% B over 10 minutes followed by 20% B to 30% B over 10 minutes. Full scan mass spectra were acquired on the orbitrap using negative ion detection over a range of m/z 300 – 800 at 70,000 resolution (m/z 300). Nucleotide identification was based on accurate mass (± 5 ppm) and agreement with the HPLC retention time of authentic standards. Quantification was accomplished by integration of extracted ion chromatograms of each metabolite, followed by comparison with corresponding standard curves. Six nucleotides fell below our detection limit using this approach (dGMP, dGDP, dGTP, GMP and the cyclic nucleotides cAMP and cGMP). Tests for significance were undertaken using Graphpad and Q-value as follows: Nucleotides that differed significantly between strains, or within strain but between larval stages, were identified using *Student's t-test* (significance defined as $p<0.05$). Results were then adjusted for multiple comparisons using the False Discovery Rate approach (5% FDR, equal variance assumption). *Student's t-test (*$p<0.05$) was also used to determine whether nucleotide pool sizes differed between strains. For these analyses, mean values of relevant nucleotide species were summed and their individual standard errors were used to calculate a nucleotide pool standard error using routine error propagation techniques.

**R-loop Quantification**. To quantify R-loop formation in worms exposed to mitochondrial ETC disruption, we used an antibody selective for DNA::RNA hybrids [18]. Briefly, 5000 synchronized worms were cultured on bacterial feeding RNAi targeting *isp-1* (1/10$^{th}$ strength), *atl-1* (9/10$^{th}$ strength), both genes combined (1/10$^{th}$ strength *isp-1* + 9/10$^{th}$ strength *atl-1*), or vector control, from the L1 larval stage to the first day of adulthood. Animals were collected in S-Basal, washed thrice in the same medium to remove eggs and bacteria, then flash frozen in liquid $N_2$. Genomic DNA (gDNA) was subsequently extracted using a DNeasy kit (Qiagen), ensuring both proteinase K and RNase A steps were included in the purification protocol. 250 ng of gDNA from each sample was next transferred to a Hybond H+ Nylon membrane (GE Healthcare) using a Hybri-Slot manifold (BRL). The DNA was UV-crosslinked to the membrane (Stratalinker 1800, Stratagene), then blocked in 5% fat-free milk powder dissolved in TBS-T (Tris-buffered saline + 0.1% Tween-20, abbreviated 5% MP). DNA::RNA hybrids were detected using αS9.6 primary antibody (Kerafast, 1:2000 in 5% MP, 16 hours, 4˚C), goat α-mouse secondary antibody (Santa Cruz Biologicals, 1:2000 in TBS-T) and ECL (GE Healthcare). Concurrently, identical gDNA samples were run on a 1% agarose gel to separate gDNA by size. This gel was southern-blotted to a Hybond H+ Nylon membrane then incubated with αS9.6 primary antibody, as described above. All resulting data images were quantified using ImageJ. Significance testing was undertaken using Student's t-test, with $p<0.05$ considered statistically significant.

**Spliceosome Reporter Assays.** To test if worms exposed to mitochondrial ETC disruption have reduced mRNA splicing efficiency, we utilized the *egl-15* and *ret-1* alternate splicing reporter genes engineered by Kuroyanagi and colleagues [19, 20] (strains KH1125, KH2283, KH928, **Table S1**). Briefly, the *egl-15* reporter gene fluoresces green when exon 5B is incorporated but red in its absence. The *ret-1* splicing reporter is comprised of a pair of synthetic genes encoding different fluorescence reporter proteins that have been frame-shifted relative to each other such that they are sensitive to the presence or absence of exon 5 for their correct translation. Synchronized eggs from each reporter strain were placed on relevant bacterial feeding RNAi lawns then cultured until day one of adulthood (20˚C). Worms were immediately imaged using an Olympus SZX16 fluorescence dissecting microscope connected to an Olympus DP71 CCD camera, using filters for GFP and RFP/mCherry.

**Nematode Oxygen Consumption:**

Nematode oxygen consumption measurements using a Seahorse XFe24 Analyzer were undertaken as follows: Synchronized N2 worm populations were cultured on HT115 bacterial feeding RNAi targeting *isp-1* (1/10$^{th}$ strength), *atl-1* (9/10$^{th}$ strength), both genes combined (1/10$^{th}$ strength *isp-1* + 9/10$^{th}$ strength *atl-1*) or vector control, from the L1 larval stage to the first day of adulthood. Animals were collected, washed thrice, then pre-incubated for 30 minutes at RT with rocking to clear the gut of live bacteria, all in S-Basal media. Approximately 30 worms were added to each Seahorse well, with 5 replicate wells employed per test condition. Four test wells were always set aside as worm-free control wells. The Seahorse measurement routine consisted of a 2 minute mix, a 2 minutes wait, then a 2 minute data collection phase, looped 8 times. We followed the procedure of Luz and colleagues [21], and only averaged oxygen consumption rates across the final four measurement phases for each well to provide a single rate value for that well. For each experiment, the 5 biological replicates (wells) were averaged per condition. A total of four independent experiments was performed. Significance testing was undertaken using the Student's t-test, with $p<0.05$ considered significant.

**Polysome Profiling.** Quantitative polysome profiling was undertaken using the procedure of Steffen and colleagues [22] with the following modifications for *C. elegans:* Synchronized nematode populations (50,000 worms per condition) were cultured on bacterial feeding RNAi targeting *isp-1* (1/10$^{th}$ strength), *atl-1* (9/10$^{th}$ strength), both genes combined (1/10$^{th}$ strength *isp-1* + 9/10$^{th}$ strength *atl-1*) or vector control, from the L1 larval stage to the first day of adulthood. After freshly preparing all solutions on the day of the experiment, worms were washed thrice with S-Basal supplemented with 100 µg/mL cycloheximide. During each wash, worms were allowed to settle under gravity (~5 mins). Worms samples were next homogenized in Lysis Buffer (40mM Tris-HCl pH 8.0, 235mM KCl, 8mM $MgCl_2$, 0.8 mM EGTA, 160 ng/mL heparin, 160 ng/mL cycloheximide, 1.95mM PMSF, 160 Units RNase inhibitor, 0.8% Triton X-100, 0.08% sodium deoxycholate) using a Balch homogenizer, as described [16]. Cycloheximide is a translational elongation inhibitor and was added to block ribosomes from running off mRNA transcripts during extraction. Heparin stabilizes mRNA/ribosome complexes. Triton X-100 and sodium deoxycholate were added to collect both membrane bound- and cytosolic ribosomes. Equal quantities of whole-worm lysate (900 µL final volume, corresponding to either 1.5 or 2.5 $OD_{260}$ Units, depending on the replicate) were loaded onto the top of a 7-47% sucrose gradient then centrifuged in a Beckman SWTi40 rotor at 38000 rpm for 2 hours at 4˚C. Sucrose gradients were prepared directly in centrifuge tubes by layering 5.5 ml 7% sucrose on top of the same volume of 47% sucrose, both prepared in Gradient Buffer (1.6 M KCl, 30mM $MgCl_2$, 100mM Tris-HCl pH 7.5, 1mg/ml heparin, 100 µg/ml cycloheximide). Centrifuge tubes were placed horizontally at RT for 3 hours to allow gradient formation. After loading they were used immediately. Following centrifugation, sample gradients were fractionated using a BR-188 Density Gradient Fractionation System (Brandel), employing 55% sucrose in Gradient Buffer as the chase solution (flow rate: 750 µL/min). Ribosomal RNA-containing elution peaks were monitored via UV absorbance (254 nm) until the entire gradient had been fractionated (~ 30 minutes). Two independent experimental replicates were collected.

**Western Analysis.** Synchronous populations of SLR001 and SLR004 worms (**Table S4**) were cultured on bacterial feeding RNAi targeting *atp-3* (1/10$^{th}$ strength and undiluted) or vector control, from the L1 larval stage to the first day of adulthood. Unbalanced individuals, representing homozygous *atl-1(tm853)* and wild type worms, respectively, were then manually selected from each plate and whole-worm extracts prepared for western blotting. Briefly, worms were collected and washed thrice in ice-cold S-basal. Next, a cocktail of protease inhibitors in 1.5% SDS (P2714, Sigma Aldrich) was added to the pellet then whole-worm extracts prepared by boiling for 5 minutes. After centrifugation (17,000xg, 5 mins. 4˚C) the cleared supernatant was retained. For western blotting we used the following reagents and suppliers: 12% NuPAGE gels (Invitrogen); nitrocellulose membrane (Protran, BA83); 5% milk powder in Tris-buffered saline + 0.05% Tween-20 ($TBS\text{-}T_{0.05\%}$) for blocking (1 hour, RT) ; $TBS\text{-}T_{0.05\%}$ for washing steps, α-ICD-1 primary antibody (1:2000 dilution, [23]) incubated at RT for 2 hours in 5% milk powder in $TBS\text{-}T_{0.05\%}$, HRP-coupled goat α-rabbit secondary antibody (ab6721, Abcam) incubated at RT for one hour. α-ICD-1 reactive bands were detected using chemiluminescence (Pierce) and a Typhoon Imaging System (GE Life Sciences).

**DNA Damage Response (DDR) Screen.**

Library Construction: Bacterial feeding RNAi targeting 201 DNA repair-related genes were sub-cloned from the Ahringer *C. elegans* RNAi Collection (Source Bioscience) into 96 well stock plates. Genes related to DDR repair were identified using WormMine (www.wormbase.org). For details see Torgovnick *et. al*. 2018 [24].

*atm-1(gk186); atl-1(tm853)* Suppressor Screen: Genes underlying the recovery in sensitivity of unbalanced *atm-1(gk186); atl-1(tm853)* worms (derived from strain SLR005) to *atp-3* RNAi were identified as follows - bacterial feeding RNAi targeting *atp-3* ($OD_{590}$ 0.39) was mixed 1:7 with each of the 201 DDR-related bacterial feeding RNAi clones (each $OD_{590}$ 0.6) then seeded onto 6-cm RNAi plates (NGM agar containing 1 mM IPTG, 100 µg/ml ampicillin, and 5 µg/ml tetracycline; final *atp-3* concentration = 1/10$^{th}$ strength) and grown overnight at room temperature. A parallel set of plates was also prepared except vector only containing bacteria was used in place of *atp-3* feeding RNAi. Each of the 402 plates was then seeded with 200 arrested L1 larvae (24 hr. arrest) from strain SLR0005. All L1 larvae were the progeny of nT1 parents. After 72 hours, plates were screened for unbalanced animals that were differentially recalcitrant to the size reducing effects of *atp-3* RNAi relative to their nT1 siblings. 16 first-round RNAi hits were obtained, which were subsequently clonally re-isolated, sequence-verified, then re-tested using *atp-3* RNAi at 1/10$^{th}$ and ½ strength. Random images of worms from 1/10$^{th}$ strength *atp-3* RNAi plates were collected using an Olympus SZX16 microscope connected to a DP71 CCD camera, then worm lengths were quantified using ImageJ (n= 20 worms per condition, per genotype, per RNAi).

*Quantitation of RNAi Effect Size and Significance Testing:* In order to quantify the recovery in insensitivity to *atp-3* feeding RNAi by unbalanced *atm-1(gk186); atl-1(tm853)* worms relative to nT1-containing *atm-1(gk186); atl-1(tm853) /+* worms, the worm length measurements collected in the previous section were combined with the following *in silico* sampling approach. For each test RNAi, four columns of 100 worm lengths were generated by randomly picking the lengths from actual worms for each of the following categories:

(i) nT1 worms treated with 9/10$^{th}$ strength test feeding RNAi + 1/10$^{th}$ strength vector feeding RNAi

(ii) nT1 worms treated with 9/10$^{th}$ strength test feeding RNAi + 1/10$^{th}$ strength *atp-3* feeding RNAi

(iii) unbalanced worms treated with 9/10$^{th}$ strength test feeding RNAi + 1/10$^{th}$ strength vector feeding RNAi

(iv) unbalanced worms treated with 9/10$^{th}$ strength test feeding RNAi + 1/10$^{th}$ strength *atp-3* feeding RNAi

The difference in lengths between the unbalanced worms with and without *atp-3* RNAi was divided by the difference in lengths between the nT1worms with and without *atp-3* RNAi, thereby producing a randomly generated population of possible worm length ratios for each test RNAi. The above steps were repeated five times for each test RNAi to create an average ratio population. Because ratios obtained from normal variables are non-normal, non-parametric techniques using medians and median absolute distance (MAD) in place of mean and standard deviation were employed in tests of significance (**Table S1**, and see [25] for a discussion on robust statistics). Data were then transferred to SPSS (IBM) for significance testing. Using the Kruskal-Wallis Test for all samples, a significant effect of test RNAi on worm length was revealed [**Table S2**, chi-squared(15)=288.36, $p < 0.001$]. Three nonparametric two-sample tests were then used as post-hoc tests to compare each test RNAi against vector: Mann-Whitney U Test, Kolmogorov-Smirnov Two-Samples Test, and the Wald-Wolfowitz Runs Test (**Table S3**). Five test RNAi clones yielded significant results for all three post-hoc tests (each $p<0.001$): *vgln-1*, *ubc-1* (independent isolate #2), *rfc-3, scc-3* and Y47D3A.29 (**File S1**). Worms treated with vector in place of test RNAi had a median ratio of 1.61, and a MAD of 0.17. Worms treated with *vgln-1*, *ubc-1* and *rfc-3* had significantly higher ratios than that of the vector (M=1.78, MAD=0.46; M=1.72, MAD=.16; M=1.86, MAD=.35, respectively), while worms treated with *scc-3* and Y47D3A.29 had significantly lower ratios (M=1.44, MAD=.31; M=1.11, MAD=0.37, respectively).

### SUPPLEMENTAL FIGURE LEGENDS

**Figure S1. Loss of ATL-1 desensitizes worms to mitochondrial respiratory chain stress induced by ethidium bromide.**

(**A, B**) Low doses of the potentiometric chemical mutagen ethidium bromide (EtBr) preferentially accumulate in mitochondria, resulting in reduced ETC function as a consequence of disruption to mtDNA integrity. **(A)** Wild type worms cultured from the egg stage to adulthood on either 5 or 25 µg/ml EtBr show a reduction in final adult size. This size reduction is noticeably abrogated by presence of the *atl-1(tm853)* mutation. Representative images were collected randomly. The lengths of the worms shown in each panel are quantified in (**B**). Bars represent mean +/- range.

**Figure S2. DNA mutation detection in *C. elegans* using β-Gal frame-shift- and TUNEL assays.**

(**A**) No evidence for DNA breaks in *isp-1(qm150)* mutants using TUNEL staining. Strain genotype and treatment condition are listed on *left.* Shown are 3 µM sections of paraffin-embedded worms. Positive controls include wild type (N2) and *nuc-1(e1392)* worms treated with gamma (γ) radiation (1344 Gy), mammalian cells undergoing apoptosis, and worms exposed to osmotic stress (350 mM NaCl). *nuc-1* encodes a DNase II required for DNA degradation during apoptosis. (Note that treatment with 350 mM NaCl was previously reported to increase DNA strand breakage [14], but we could find no support for this result in our studies, despite following the same methodology.)

(**B**) P*hsp-16.2::ATG(A)$_{17}$::GFP::LacZ* and P*hsp-16.2::ATG::GFP::LacZ* both encode out-of-frame reporter genes [13]. Mutagenic events that bring the reading frame of either reporter back into frame result in a detectable event (LacZ positive staining). Both germline and somatic events are detectable using this approach. HT115 bacteria contain a functional lac operon (blue worms, *vector-only treatment*) that in our hands precludes use of this reporter with the Ahringer RNAi bacterial feeding RNAi library, contrary to previous reports [13].

**Figure S3. Nucleotide biosynthesis in *C. elegans* highlighting the role of the mitochondrial ETC.**

Schematic depiction of key intermediates involved in the genesis of pyrimidine nucleotides in worms. The mitochondrial inner membrane-associated enzyme dihydroorotate dehydrogenase (DHODH, named DHOD-1 in *C. elegans*) catalyzes the oxidation of dihydroorotate to orotate and delivers its reducing equivalents directly to ubiquinone in the ETC chain.

**Figure S4. mRNA splicing remains unaltered in worms experiencing ETC disruption, both in the absence or presence of *atl-1* knockdown.**

(**A, B**) Schematic of *egl-15* (A) and *ret-1* (B) splicing-detection reporter gene and gene pair respectively [19, 26].

(**C**) Splicing function reported by the *egl-15 (left panel)* and *ret-1 (right panel)* synthetic reporter constructs remains unaltered by 1/10$^{th}$ strength *atp-3, isp-1* or *nuo-6* knockdown in age-matched worms (vector control are 3rd day adults). *atl-1* knockdown alone, or in combination with 1/10$^{th}$ strength *atp-3, isp-1* or NDUFB4*/nuo-6* RNAi, also did not alter expression of reporter constructs. The *egl-15* reporter gene in the *asd-1(yb978)* mutant background is presented as a positive control (*middle panel*). Wild type ASD-1 binds a region in intron 4 that repress selection of E5B-GFP, thereby enhancing E5A-RFP expression[26].

**Figure S5. *atl-1* knockdown does not alter retrograde response pathways normally induced by mitochondrial ETC stress.**

(**A-D**) Reporter genes monitoring the mitochondrial unfolded protein response (*Phsp-6::GFP),* the PMK-3/p38-dependent retrograde response (*Ptbb-6::GFP*) and the SKN-1 dependent oxidative stress response (*Pgst-4::GFP*) following *atp-3, isp-1* and *nuo-6* knockdown (A).

Quantification provided in *panels* (B-D). *atl-1* knockdown does not cause hyperactivation of any reporter gene, except *Ptbb-6::GFP,* which is mildly (1.6 fold) increased in *nuo-6* RNAi-treated worms. (Asterisks indicate significantly different from vector unless otherwise noted by bar over comparison group: *Student's t-test,* * p<0.05, **p<0.01, ***p<0.001, ^ p < 0.01; error bars: SEM.) (**E**) Transcript abundance of endogenous *sod-3, hsp-6* and *ugt-61* in *atl-1(tn853)* mutants relative to wild type worms (data for each gene is normalized to wild type). Transcript abundance for *atm-1(gk186)* and *atl-1(tm853); atm-1(gk186)* (line SLR0003) worms is also provided. All animals are the unbalanced F1 progeny of parents carrying the nT1 reciprocal chromosomal translocation, including wild type control. (Asterisks indicate significantly different from wild type: *Student's t-test*, * p<0.05, **p<0.01, ***p<0.001, Error bars: SEM.)

### SUPPLEMENTAL TABLE LEGENDS

**Table S1. Summary of descriptive statistics for final-round DDR test RNAi screen hits.** (Related to **Fig. 3**).

**Table S2. Mean Ranks of final-round DDR RNAi hits used for Kruskal-Wallis test.** (Related to **Fig. 3**).

**Table S3. Post-hoc tests for each final-round DDR RNAi hit versus vector.** (Related to **Fig. 3**).

**Table S4. List of *C. elegans* strains employed in current study.**

**Table S5. qPCR primer list.**

### OTHER SUPPLEMENTAL FILES

**File S2. Mean difference ratios of all hits from DNA Damage Response (DDR) screen.** (Related to **Fig. 3**)

**File S3. Significance testing for differences in HXK2::GFP fluorescence.** (related to **Fig. 7C**)

### SUPPLEMENTAL REFERENCES


1. Wood WB (ed). *The Nematode Caenorhabditis elegans*. Cold Spring Harbor Laboratory: New York, 1988.

2. Jones MR, Huang JC, Chua SY, Baillie DL, Rose AM. The atm-1 gene is required for genome stability in Caenorhabditis elegans. *Mol Genet Genomics* 2012, **287**(4)**:** 325-335.

3. Garcia-Muse T, Boulton SJ. Distinct modes of ATR activation after replication stress and DNA double-strand breaks in Caenorhabditis elegans. *Embo J* 2005, **24**(24)**:** 4345-4355.

4. Timmons L, Fire A. Specific interference by ingested dsRNA. *Nature* 1998, **395**(6705)**:** 854.

5. Rea SL, Ventura N, Johnson TE. Relationship Between Mitochondrial Electron Transport Chain Dysfunction, Development, and Life Extension in Caenorhabditis elegans. *PLoS Biol* 2007, **5**(10)**:** e259.

6. Munkacsy E, Khan MH, Lane RK, Borror MB, Park JH, Bokov AF*, et al.* DLK-1, SEK-3 and PMK-3 Are Required for the Life Extension Induced by Mitochondrial Bioenergetic Disruption in C. elegans. *PLoS Genet* 2016, **12**(7)**:** e1006133.

7. Kamath RS, Ahringer J. Genome-wide RNAi screening in Caenorhabditis elegans. *Methods* 2003, **30**(4)**:** 313-321.

8. Butler JA, Ventura N, Johnson TE, Rea SL. Long-lived mitochondrial (Mit) mutants of Caenorhabditis elegans utilize a novel metabolism. *FASEB J* 2010, **24**(12)**:** 4977-4988.

9. Schneider CA, Rasband WS, Eliceiri KW. NIH Image to ImageJ: 25 years of image analysis. *Nat Methods* 2012, **9**(7)**:** 671-675.

10. Hoogewijs D, Houthoofd K, Matthijssens F, Vandesompele J, Vanfleteren JR. Selection and validation of a set of reliable reference genes for quantitative sod gene expression analysis in C. elegans. *BMC Mol Biol* 2008, **9:** 9.

11. Sugimoto T, Mori C, Takanami T, Sasagawa Y, Saito R, Ichiishi E*, et al.* Caenorhabditis elegans par2.1/mtssb-1 is essential for mitochondrial DNA replication and its defect causes comprehensive transcriptional alterations including a hypoxia response. *Experimental Cell Research* 2008, **314**(1)**:** 103-114.

12. Rooney JP, Ryde IT, Sanders LH, Howlett EH, Colton MD, Germ KE*, et al.* PCR based determination of mitochondrial DNA copy number in multiple species. *Methods Mol Biol* 2015, **1241:** 23-38.

13. Pothof J, van Haaften G, Thijssen K, Kamath RS, Fraser AG, Ahringer J*, et al.* Identification of genes that protect the C. elegans genome against mutations by genome-wide RNAi. *Genes Dev* 2003, **17**(4)**:** 443-448.

14. Dmitrieva NI, Celeste A, Nussenzweig A, Burg MB. Ku86 preserves chromatin integrity in cells adapted to high NaCl. *Proc Natl Acad Sci U S A* 2005, **102**(30)**:** 10730-10735.

15. Wu Y-C, Stanfield GM, Horvitz HR. NUC-1, a Caenorhabditis elegans DNase II homolog, functions in an intermediate step of DNA degradation during apoptosis. *Genes Dev* 2000, **14**(5)**:** 536-548.

16. Bhaskaran S, Butler JA, Becerra S, Fassio V, Girotti M, Rea SL. Breaking Caenorhabditis elegans the easy way using the Balch homogenizer: an old tool for a new application. *Analytical Biochemistry* 2011, **413**(2)**:** 123-132.

17. Huang D, Zhang Y, Chen X. Analysis of intracellular nucleoside triphosphate levels in normal and tumor cell lines by high-performance liquid chromatography. *J Chromatogr B Analyt Technol Biomed Life Sci* 2003, **784**(1)**:** 101-109.

18. Zhang ZZ, Pannunzio NR, Hsieh CL, Yu K, Lieber MR. Complexities due to single-stranded RNA during antibody detection of genomic rna:dna hybrids. *BMC Res Notes* 2015, **8:** 127.

19. Kuroyanagi H, Watanabe Y, Suzuki Y, Hagiwara M. Position-dependent and neuron-specific splicing regulation by the CELF family RNA-binding protein UNC-75 in Caenorhabditis elegans. *Nucleic Acids Res* 2013, **41**(7)**:** 4015-4025.

20. Kuroyanagi H, Ohno G, Sakane H, Maruoka H, Hagiwara M. Visualization and genetic analysis of alternative splicing regulation in vivo using fluorescence reporters in transgenic Caenorhabditis elegans. *Nat Protoc* 2010, **5**(9)**:** 1495-1517.

21. Luz AL, Rooney JP, Kubik LL, Gonzalez CP, Song DH, Meyer JN. Mitochondrial Morphology and Fundamental Parameters of the Mitochondrial Respiratory Chain Are Altered in Caenorhabditis elegans Strains Deficient in Mitochondrial Dynamics and Homeostasis Processes. *PLoS One* 2015, **10**(6)**:** e0130940.

22. Steffen KK, MacKay VL, Kerr EO, Tsuchiya M, Hu D, Fox LA*, et al.* Yeast life span extension by depletion of 60s ribosomal subunits is mediated by Gcn4. *Cell* 2008, **133**(2)**:** 292-302.

23. Bloss TA, Witze ES, Rothman JH. Suppression of CED-3-independent apoptosis by mitochondrial [beta]NAC in Caenorhabditis elegans. *Nature* 2003, **424**(6952)**:** 1066-1071.

24. Torgovnick A, Schiavi A, Shaik A, Kassahun H, Maglioni S, Rea SL*, et al.* BRCA1 and BARD1 mediate apoptotic resistance but not longevity upon mitochondrial stress in Caenorhabditis elegans. *EMBO Rep* 2018, **19**(12).

25. Hoaglin DC, Mosteller F, Tukey JW. *Understanding robust and exploratory data analysis*. Wiley: New York, 1983.

26. Kuroyanagi H, Kobayashi T, Mitani S, Hagiwara M. Transgenic alternative-splicing reporters reveal tissue-specific expression profiles and regulation mechanisms in vivo. *Nat Methods* 2006, **3**(11)**:** 909-915.
