## Supplemental Table S1 for "Inhibition of ATR Reverses a Mitochondrial Respiratory Insufficiency"

**Table S1**. *Summary of Descriptive Statistics for final-round DDR RNAi Screen Hits*

| Test RNAi | Median | MAD | Variance | Skewness | Kurtosis | Min | Max |
| --- | --- | --- | --- | --- | --- | --- | --- |
| Vector | 1.61 | 0.17 | 0.25 | 1.92 | 7.81 | 1.18 | 3.00 |
| *crn-1* | 1.84 | 0.28 | 0.17 | 0.67 | 0.16 | 1.11 | 3.16 |
| *hpr-17* | 1.69 | 0.34 | 7.57 | 7.78 | 71.75 | -3.67 | 27.44 |
| *ung-1 (isolate 1)* | 1.70 | 0.24 | 0.14 | 0.77 | 1.28 | 0.98 | 3.07 |
| *ung-1 (isolate 2)* | 1.55 | 0.21 | 0.07 | 1.03 | 1.98 | 1.08 | 2.54 |
| *C08H9.2* | 1.94 | 0.26 | 1.03 | 4.77 | 27.71 | 1.36 | 8.99 |
| *ubc-1 (isolate 1)* | 1.72 | 0.16 | 0.05 | 0.34 | -0.46 | 1.33 | 2.35 |
| *ubc-1 (isolate 2)* | 1.81 | 0.21 | 0.11 | 0.91 | 0.79 | 1.31 | 3.09 |
| *ubc-1 (isolate 3)* | 1.71 | 0.30 | 0.16 | 2.73 | 11.40 | 1.15 | 3.94 |
| *rfc-3* | 1.86 | 0.35 | 2.37 | -4.14 | 27.15 | -8.50 | 5.57 |
| *lin-40* | 1.70 | 0.18 | 0.07 | 0.85 | 1.92 | 1.10 | 2.83 |
| *gsp-1* | 1.57 | 0.27 | 0.18 | 0.85 | 0.49 | 0.83 | 2.97 |
| *scc-3* | 1.44 | 0.31 | 0.30 | 0.40 | -0.18 | 0.38 | 2.88 |
| *sir-2.2* | 1.67 | 0.21 | 0.13 | 1.42 | 4.05 | 1.17 | 3.48 |
| *Y47D3A.29* | 1.22 | 0.21 | 0.12 | 1.00 | 1.02 | 0.73 | 2.52 |
| *ape-1* | 1.86 | 0.21 | 0.25 | 3.87 | 23.55 | 1.31 | 5.41 |

*Note. N=100 for all groups; superscript indicates replicate number.*
