## Supplemental Table S2 for "Inhibition of ATR Reverses a Mitochondrial Respiratory Insufficiency"

**Table S2.** *Mean Ranks of Final-round DDR RNAi Hits used for Kruskal-Wallis Test*

| Test RNAi | N | Mean Rank |
| --- | --- | --- |
| Vector | 100 | 692.54 |
| *crn-1* | 100 | 969.4 |
| *hpr-17* | 100 | 837.69 |
| *ung-1* (isolate 1) | 100 | 788.82 |
| *ung-1* (isolate 2) | 100 | 581.33 |
| *C08H9.2* | 100 | 1130.45* |
| *ubc-1* (isolate 1) | 100 | 833.1 |
| *ubc-1* (isolate 2) | 100 | 998.47* |
| *ubc-1* (isolate 3) | 100 | 836.1 |
| *rfc-3* | 100 | 989.36* |
| *lin-40* | 100 | 833.11 |
| *gsp-1* | 100 | 693.31 |
| *scc-3* | 100 | 522.83* |
| *sir-2.2* | 100 | 825.33 |
| *Y47D3A.29* | 100 | 324.45* |
| *ape-1* | 100 | 951.71 |

**marks test RNAis significantly different from the vector as revealed in post-hoc analysis*
