## Supplemental Table S3 for "Inhibition of ATR Reverses a Mitochondrial Respiratory Insufficiency"

**Table S3.** *Post-hoc Tests for Each Final-round DDR RNAi Hit versus Vector*

| Test RNAi | Significance Test^a^ | | Significance | Retain or Reject Null Hypothesis? |
| --- | --- | --- | --- | --- |
| *crn-1* | | Mann – Whitney U Test | .000 | Reject* |
|  | | Kolmogorov – Smirnov Test | .000 | Reject* |
|  | | Wald – Wolfowitz Runs Test | .024 | Retain |
| *hpr-17* | | Mann – Whitney U Test | .159 | Retain |
|  | | Kolmogorov – Smirnov Test | .000 | Reject* |
|  | | Wald – Wolfowitz Runs Test | .024 | Retain |
| *ung-1 (isolate 1)* | | Mann – Whitney U Test | .136 | Retain |
|  | | Kolmogorov – Smirnov Test | .002 | Retain |
|  | | Wald – Wolfowitz Runs Test | .000 | Reject* |
| *ung-1 (isolate 2)* | | Mann – Whitney U Test | .011 | Retain |
|  | | Kolmogorov – Smirnov Test | .024 | Retain |
|  | | Wald – Wolfowitz Runs Test | .008 | Retain |
| *C08H9.2* | | Mann – Whitney U Test | .000 | Reject* |
|  | | Kolmogorov – Smirnov Test | .000 | Reject* |
|  | | Wald – Wolfowitz Runs Test | .000 | Reject* |
| *ubc-1 (isolate 1)* | | Mann – Whitney U Test | .003 | Retain |
|  | | Kolmogorov – Smirnov Test | .002 | Retain |
|  | | Wald – Wolfowitz Runs Test | .012 | Retain |
| *ubc-1 (isolate 2)* | | Mann – Whitney U Test | .000 | Reject* |
|  | | Kolmogorov – Smirnov Test | .000 | Reject* |
|  | | Wald – Wolfowitz Runs Test | .001 | Reject* |
| *ubc-1 (isolate 3)* | | Mann – Whitney U Test | .004 | Retain |
|  | | Kolmogorov – Smirnov Test | .010 | Retain |
|  | | Wald – Wolfowitz Runs Test | .024 | Retain |
| *rfc-3* | | Mann – Whitney U Test | .000 | Reject* |
|  | | Kolmogorov – Smirnov Test | .000 | Reject* |
|  | | Wald – Wolfowitz Runs Test | .000 | Reject* |
| *lin-40* | | Mann – Whitney U Test | .005 | Retain |
|  | | Kolmogorov – Smirnov Test | .010 | Retain |
|  | | Wald – Wolfowitz Runs Test | .033 | Retain |
| *gsp-1* | | Mann – Whitney U Test | .471 | Retain |
|  | | Kolmogorov – Smirnov Test | .006 | Retain |
|  | | Wald – Wolfowitz Runs Test | .004 | Retain |
| *scc-3* | | Mann – Whitney U Test | .000 | Reject* |
|  | | Kolmogorov – Smirnov Test | .000 | Reject* |
|  | | Wald – Wolfowitz Runs Test | .000 | Reject* |
| *sir-2.2* | | Mann – Whitney U Test | .025 | Retain |
|  | | Kolmogorov – Smirnov Test | .006 | Retain |
|  | | Wald – Wolfowitz Runs Test | .197 | Retain |
| *Y47D3A.29* | | Mann – Whitney U Test | .000 | Reject* |
|  | | Kolmogorov – Smirnov Test | .000 | Reject* |
|  | | Wald – Wolfowitz Runs Test | .000 | Reject* |
| *ape-1* | | Mann – Whitney U Test | .000 | Reject* |
|  | | Kolmogorov – Smirnov Test | .000 | Reject* |
|  | | Wald – Wolfowitz Runs Test | .012 | Retain |

*Note. *p<.001*

^a^These are nonparametric significance tests comparing each test RNAi to the vector.
